# Bioprospecting c-di-GMP activated exopolysaccharides in bacteria

**DOI:** 10.1101/2025.09.12.675316

**Authors:** Daniel Perez-Mendoza, Jochen Schmid, Manuel Döring, Broder Rühmann, Miguel Ángel Rodríguez-Carvajal, Manuel Bermudo-Molina, Volker Sieber, Juan Sanjuan

## Abstract

The genetic and physiological diversity of bacteria are critical resources to discover new exopolysaccharides (EPS) as raw materials with biotechnological applications. However, uncovering new EPS is limited by their lack of production in laboratory cultures, as EPS are often cryptic and their biosynthesis only proceed upon unknown environmental cues. The dinucleotide cyclic di-GMP (c-di-GMP) has emerged as a universal second messenger in bacteria and common activator of many EPS. Here, a genetic modification to elevate intracellular c-di-GMP levels and a carbohydrate fingerprinting analysis, were combined for a High-Throughput Screening (HTS) of 330 bacterial strains in search of c-di-GMP activated EPS. Nearly 10% of strains were revealed as promising candidates to overproduce novel EPS composites, in a c-di-GMP dependent manner. In these conditions, *Sphingomonas* sp. C10 massively produced a EPS with an unusual monosaccharide composition, compared to known biotechnologically relevant sphingans.

## Introduction

A large variety of biopolymers (i.e. polymeric substances of natural origin), such as polyesters, polyamides and polysaccharides, are regularly produced by living organisms. Such renewable and biodegradable compounds could replace oil-based commodity materials to provide solutions for a wide range of applications in many industrial sectors [1]. Biobased polymers contribute to the preservation of fossil-based raw materials and the reduction of carbon dioxide released into the atmosphere, while exhibiting enhanced environmental compatibility. Bacteria, in particular, can be considered prime cell factories that are able to efficiently convert nitrogen and carbon sources into valuable biomaterials. Furthermore, bacteria-produced polymers are also a desirable alternative to those of plant and algal origin, owing to the high growth rates of the producing bacteria, the reproducible physicochemical properties of their polymers, and their metabolic flexibility and adaptability to industrial processes. Overall, these features lead to increasing competitiveness and reducing production costs.

Among bacterial polymers, exopolysaccharides (EPS) are secreted macromolecules with different compositions, functions and physical and chemical characteristics. EPS can remain attached to the cell surface forming sheaths, capsules or slimes or, on the contrary, be released into the medium as either a soluble or insoluble material, thereby contributing to the biofilm matrix, protection or signaling functions [2]. In terms of biotechnological applications, bacterial EPS are among the most interesting biopolymers and have proven their usefulness in many different industrial applications such as textiles, detergents, adhesives, oil recovery, wastewater treatment, dredging, brewing, downstream processing, cosmetology, pharmacology and as food additives [3, 4]. Some examples of these biotechnological relevant EPS are xanthan gum, hyaluronic and colanic acid, gellan, alginate or bacterial cellulose [5-7].

EPS are synthesized and exported through several distinct mechanisms that occur in intra- and extracellular ways [8]. However, key conserved components have been identified in many described secretion systems, including an inner-membrane-embedded Glycosyltransferase (GT), periplasmic scaffold proteins, and outer-membrane β-barrel porins [9]. There is also increasing evidence that these biomachinery complexes are usually strictly regulated by environmental cues and often share common regulatory schemes. This is due to the high costs of energy and nutrients that EPS biosynthesis and secretion impose on the producing cells. For instance, up to date more than a dozen EPS are known to be regulated by the bacterial second messenger bis- (3’,5’)-cyclic diguanosine monophosphate (cyclic diguanylate, c-di-GMP) including biotechnological important ones like the widespread bacterial cellulose, poly-β(1→6)-*N*-acetyl-d-glucosamine, curdlan, xanthan, alginate, Psl and Pel [10].

Cyclic-di-GMP is currently considered a universal lifestyle switch molecule in bacteria, with a crucial role in governing the transition from a planktonic/motile to a sessile biofilm mode of growth, but also with a great influence on a wide range of cellular processes, including cell-cell signaling, cell cycle progression and virulence [11, 12]. Cyclic-di-GMP signaling systems are generally composed of four major constituents: (i) diguanylate cyclases (DGCs, synthesize c-di-GMP from two GTP molecules), (ii) phosphodiesterases (PDEs, degrade c-di-GMP), (iii) c-di-GMP binding effectors which interact with (iv) target components to produce a molecular output [13, 14]. Its involvement in the bacterial decision to attach to a surface and form a biofilm community makes this molecule a key activator for the production and secretion of different biofilm matrix components, including many EPS [15]. Many of the biosynthetic machineries of c-di-GMP regulated biopolymers are enzymatic complexes, either repressed or exhibiting inactive conformations, which are turned on by c-di-GMP in combination with often unknown environmental or physiological cues [16]. This is particularly relevant in environmental strains, where EPS production is often cryptic and passes unnoticed for researchers in the laboratory. Based on available bacterial carbohydrate databases, e.g. [17], and considering the limited number of bacterial species and strains that have been explored for EPS production, it can be stated that the natural diversity of bacterial biopolymers remains largely unexplored [10]. The growing demand for sustainable products further increases the need for the discovery of new EPS. Artificially elevating the bacterial c-di-GMP contents may then result in a wise approach to unveil natural EPS, by deregulating its biosynthesis and promoting its massive production under laboratory conditions. Our recently discovered Mixed-Linkage β-Glucan (MLG) in Rhizobiaceae is a paradigmatic example [18]. MLG was detected after artificially increasing the intracellular levels of c-di-GMP in *Sinorhizobium meliloti*, a well-characterised rhizobial model bacterium, whose genome was disclosed more than 20 years ago and which has been carefully studied for the production of different EPS for decades [19-21]. However, MLG biosynthesis is cryptic and its production under laboratory conditions went unadverted for many laboratories worldwide. Raising the intracellular levels of c-di-GMP allowed us to discover this unique bacterial EPS never described in bacteria before. Artificially increasing the intracellular levels of this second messenger rendered also the overproduction of other valuable EPS in diverse environmental bacteria [22-24]. These examples support our genetic approach as a valid strategy to overproduce biotechnological relevant EPS under laboratory conditions. But most importantly, it could be a powerful tool to uncover cryptic EPS that remain to be discovered [25].

The biological diversity present in natural bacterial isolates and culture collections is an excellent source for the exploration of novel EPS, but requires efficient, fast and reliable screening approaches for identifying microorganisms capable of producing EPS in substantial amounts [26]. In this work, we have combined the genetic approach of raising the intracellular levels of c-di-GMP with a carbohydrate fingerprinting analysis using a High Throughput Screening (HTS) method with 330 different environmental bacterial strains for prospecting and exploration for novel EPS.

## Results

### Genetic modification of large collections of bacteria from different biogeographical origins

A total of 330 bacterial strains from different phylogenetic taxa and various geographical and environmental origins were screened for c-di-GMP activated EPS (Table S1). On one hand, there was a set of 86 selected bacteria from our laboratory collection, including: (i) 35 reference strains, (ii) 41 legume nodule isolates from different host plants and habitats, including bean isolates from Granada (Spain) [27], alfalfa symbionts from Argentina [28] and soybean nodule isolates from China [29]; and (iii) 10 peppermint *Pseudomonas* plant growth promoting rhizobacteria (PGPR) from Argentinian soils [30]. All together, these 86 strains represent a heterogeneous group formed manly by plant-interacting rhizobacteria. Rhizosphere represents an interesting reservoir of diverse microorganisms capable of producing numerous EPS playing important roles in the colonization of their host plant roots [31-33]. This bacterial collection was genetically modified by individually transforming each strain with a plasmid (pJBpleD*), harbouring a constitutively active variant of the well-characterised DGC PleD from *Caulobacter crescentus* [23], or with the respective empty vector (pJB3Tc19), following standard conjugation protocols (Table S2). PleD* has been shown to be an active DGC in different genetic backgrounds, capable of rising the intracellular levels of c-di-GMP by more than one order of magnitude in several bacterial strains, including different rhizobacteria [23, 34].

Additionally, a High Throughput conjugation protocol was set up for the simultaneous genetic modification of large numbers of bacterial strains. We adapted previously described protocols for high throughput automated conjugation [35, 36] in order to transfer pJBpleD* or the empty vector pJB3Tc19 into recipient strains growing in 96-deep-well plates (DWP). The protocol was optimized to be performed either manually with multichannel pipettes, or automatized with liquid handling workstations which are especially suitable for HTS approaches. Two different bacterial collections were used: (i) two 96-well plates containing a total of 182 bacterial isolates from different environmental habitats (MTP-I and MTP-II; [37]), and (ii) a 96-well plate with 62 *Sphingomonas* spp. isolates (SPH). The genus *Sphingomonas* has attracted much attention in recent years for its ability to produce high-molecular-weight extracellular polymers called sphingans, including biotechnological interesting EPS like welan, gellan and diutan gum [38, 39]. It must be noted that the 94 bacterial strains in MTP-II plate have been previously analysed under physiological level of c-di-GMP, using the same EPS HTS platform, with the results of 41 of them producing at least one detectable EPS [37]. Each strains from MTP-I, MTP-II and SPH were conjugated by using a handing-liquid station with *E. coli* β2163 donor strains harbouring the pJBpleD* or the empty vector pJB3Tc19, respectively (Table S2).

In summary, 2 variants of each strain were generated in the same 96 well plate: one with pJBpleD* (even columns), expected to express high intracellular levels of c-di-GMP [23, 34], and another one with the empty vector pJB3Tc19 (odd columns), exhibiting physiological levels of c-di-GMP. Blanks at different positions were included in the plates as negative controls (Table S3).

### HTS platform to identify bacterial strains producing c-di-GMP activated EPS

The transformed strains were analysed following a previously described protocol based on a fast carbohydrate analysis via liquid chromatography coupled with ultraviolet and electrospray ionization ion trap detection in 96-well format [37]. The core of the method is the identification of quantitative, but also, qualitative changes (monosaccharide composition) promoted by c-di-GMP on the polysaccharides formed during microbial growth. This approach involves five different steps, all in 96-well format: (i) cultivation of the strains, (ii) centrifugation to remove bacterial cells, (iii) gel-filtration of the supernatants for the complete removal of the remaining carbohydrates from the cultivation media, iv) the polysaccharide hydrolysis and HT-PMP derivatization of the polymer(s) and the (v) final analysis of the monomeric composition by UHPLC–UV–ESI–MS, see Methods for details [37].

Different pre-screening analytical modules are also included during the HTS approach in order to give additional information to preselected candidates, reducing the number of the strains designed for the carbohydrate-fingerprint analysis. This increases the efficiency and reduces consumables and measurement time during the screening process.

The choice of a suitable cultivation medium is also essential for this HTS. Complex media containing poly- and oligomeric carbohydrate compounds as carbon source (e.g. yeast extract), should be avoided as might negatively interfere with the carbohydrate-fingerprint analysis [40]. Strains from MTP-IA &-IB, MTP-IIA &-IIB and SPH plates were grown using EPS media which was found to induce EPS production for many bacterial strains in previous experiments and contain glucose as the sole carbon source [26]. Strains from GRI and GRII were grown in a Minimal Media (MM) previously reported to be able to sustain growth of different rhizobacteria [41]. This MM contains mannitol as a principal carbon source, which is not subjected to the PMP derivatization and thus prevents unwanted interference with the subsequent quantitative analysis of the monosaccharide composition of the EPSs.

One of the unavoidable steps to complete the carbohydrate-fingerprint analysis implies the complete removal of the bacterial cells to prevent interference with the correct determination of the carbohydrates from the polymer(s). In principle, this would remove also cell-associated EPS during the centrifugation step. Thus, to grab the full potential of the different strains on EPS production, we implemented the HTS with a previous growth of the strains in solid media supplemented with 3 different dyes: Congo Red (CR), Aniline Blue (AB) and Calcofluor (CF) (Figure S1). Several bacterial polymers have been shown to specifically interact with dyes, widely used to highlight EPS production in previous EPS screenings and characterization studies [23, 42-44]. AB, for example, shows specificity to β-(1-3)-glucans and other structurally related EPS [45]. CF binds to succinoglycan as well as to different pure and mixed-linkage β-glucans and exhibits a blue-green fluorescence when irradiated by long-wave UV light [18]. CR binds to α-d-glucopyranosyl units, basic or neutral polysaccharides, as well as to some proteins [42, 46]. This step, which also included observation of flocs and biofilm formation in liquid media, was added to the analytical modules described by Rühmann *et al*. [40].

In general, the over-expression of *pleD*\* generates a deep impact on the colony morphotype (morphology and color) of many of the assayed strains (Figure S1). Most frequent phenotypes include *pleD\**-induced wrinkled colonies with increased staining with one or more dyes (see for example Figure S1: E10 *vs*. E9 of GRI plate). Several strains expressing *pleD*\* also switched to an intense fluorescence in CF plates, in contrast to their respective controls (see for example Figure S1: G2 *vs*. G1 or G8 *vs*. G7 of GRI plate). Other strains did not exhibit appreciable changes in binding to any of the dyes but great changes in colony appearance, also suggesting a boost on EPS production (see for example G12 vs. G11 of GRII plate in Figure S1).

In order to decide which strains would be subjected to the complete carbohydrate fingerprint analysis, a key factor taken into account was the total carbohydrate content (TCC) of the *pleD*\* *vs*. the empty vector culture supernatants. All strains were subjected to a fully automated fast detection method of the total sugar content, based on absorbance measurement after a phenol-sulfuric-acid treatment (Table S3). The phenol-sulfuric-acid method is one of the easiest and most reliable methods to measure the total carbohydrate content and it has been previously optimized to be use in a 96-well microplates format [47]. To minimize the effect of the different bacterial growing rates on the amounts of EPS production, the TCC was standardised by two different parameters: (i) the optical density (OD_600nm_) of the culture and (ii) the Total Protein Content (TPC) of the cell pellets.

The TCC ratio of *pleD*\*/WT was >1 for 43% of all the assayed strains, with a TCC average ratio of 1.26 (Figure 1 and Table 1). These values increased when the TCC was standardised by OD, up to 55% of the assayed strains with TCC/OD ratio >1 and an average value of 1.87 (Figure 1 and Table 1). The rise was even higher when the TCC was standardised by TPC, with 59% of the assayed strains showing TCC/TPC ratios >1 and TCC/TPC average ratio of 2.76 (Figure 1 and Table 1). These results showed a general positive impact of *pleD*\* on the TCC of the assayed bacteria, indicating that, as expected, c-di-GMP promoted a generalised increase in EPS production.

**Table 1.** Total Carbohydrate Content (TCC) of the different bacterial subcollections.

| Subcollections | Ratio TCC<sup>+</sup> | Ratio TCC/OD<sup>+</sup> | Ratio TCC/TPC<sup>+</sup> |
| --- | --- | --- | --- |
| Reference Strains | 1.17 (43%) | 1.75 (57%) | 3.00 (62%) |
| Rhizobacteria isolates | 0.69 (16%) | 1.94 (67%) | 2.26 (77%) |
| Environmental Isolates | 1.39 (48%) | 1.52 (51%) | 1.58 (51%) |
| *Sphingomonas* Strains | 1.48 (53%) | 3.69 (59%) | 9.22 (61%) |
| All groups | 1.26 (43%) | 1.87 (55%) | 2.76 (59%) |

+ Mean ratios *pleD*\*/WT; *pleD*\*/WT normalised with the OD (OD<sub>600nm</sub>); *pleD*\*/WT normalised with the Total Protein Content (TPC). Percentage of assayed strains in each group with ratios >1 are indicated in brackets

**Figure 1:**
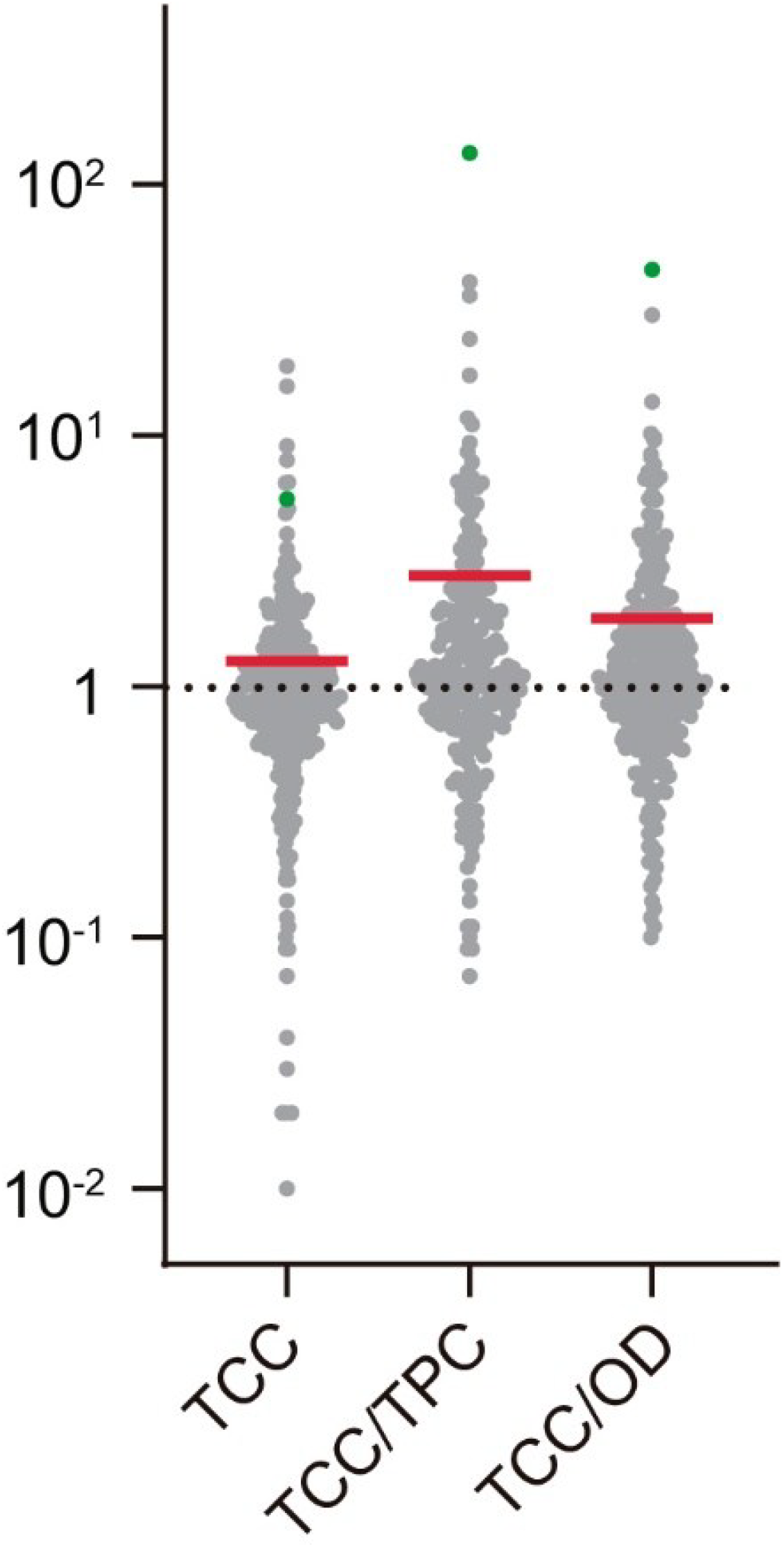
General Impact of c-di-GMP on the EPS production by all strains tested in the screen. Representation the Total Carbohydrate Content (TCC) ratios of the *pleD*\* *vs*. the empty vector culture supernatants of each strain. TCC: TCC Ratios *pleD\**/WT; TCC/OD: TCC ratios *pleD\**/WT normalised with the OD_600nm_; TCC/TPC: TCC ratios *pleD\**/WT normalised with the Total Protein Content are shown. Red lines indicate the mean of different ratios. Green dots indicate the position of *Sphingomonas* strain SphC10.

### Subcollection: Reference Strains

Reference strains assayed in GRI plate displayed a generalised positive effect on EPS production induced by c-di-GMP, with average ratios values (*pleD\**/WT) >1 in all measurements (TCC, TCC/OD and TCC/TPC; Table 1). Eight strains showed a positive effect of *pleD\**on the EPS production in all measurements, with TCC/TPC ratios ranging between 1.0 and 24.2 (Figure S2 and Table S4). In other strains, e.g. *Cupriavidus necator*, despite presenting *pleD*\*/WT ratios <1 (Figure S2), the presence of *pleD\** generated a deep impact on the colony morphotype with wrinkled colonies stained with CR and CF (Figure S1: E10 *vs*. E9 of GRI plate), thus suggesting the c-di-GMP promotion of an insoluble EPS associated to the cellular fraction.

Twelve of these 35 strains were subjected to the full carbohydrate fingerprint analysis by UHPLC–UV–ESI–MS (Figure 2A). Seven of the 12 displayed appreciable qualitative changes in their EPS compositions, with galacturonic acid being the most variable monomer. High c-di-GMP levels seems to have a dual role on the presence of galacturonic acid on the released EPS depending on the strains, reducing its presence in rhizobial strains like *Sinorhizobium meliloti* (GRI/A12 *vs*. GRI/A11 and GRI/B2 *vs*. GRI/B1; Figure 2A) or *Mesorhizobium loti* (GRI/B8 *vs*. GRI/B7; Figure 2A), and promoting it on *Pseudomonas* phytopathogenic strains like *P. syringae* (GRI/C10 *vs*. GRI/C9, GRI/C12 *vs*. GRI/C11; Figure 2A) or *P. savastanoi* (GRI/D2 *vs*. GRI/D1; Figure 2A) or the phylogenetically related *Delftia acidovorans*, in which the presence of *pleD\** increased the galacturonic acid from undetectable levels to more than 50% (GRI/F2 *vs*. GRI/F1; Figure 2A). The presence of *pleD\** in *P. savastanoi* pv. phaseolicola 1448 strain drastically affected the composition of the EPS with the appearance of dimeric sugars like gentiobiose and cellobiose among its components (GRI/D2 *vs*. GRI/D1; Figure 2A).

**Figure 2:**
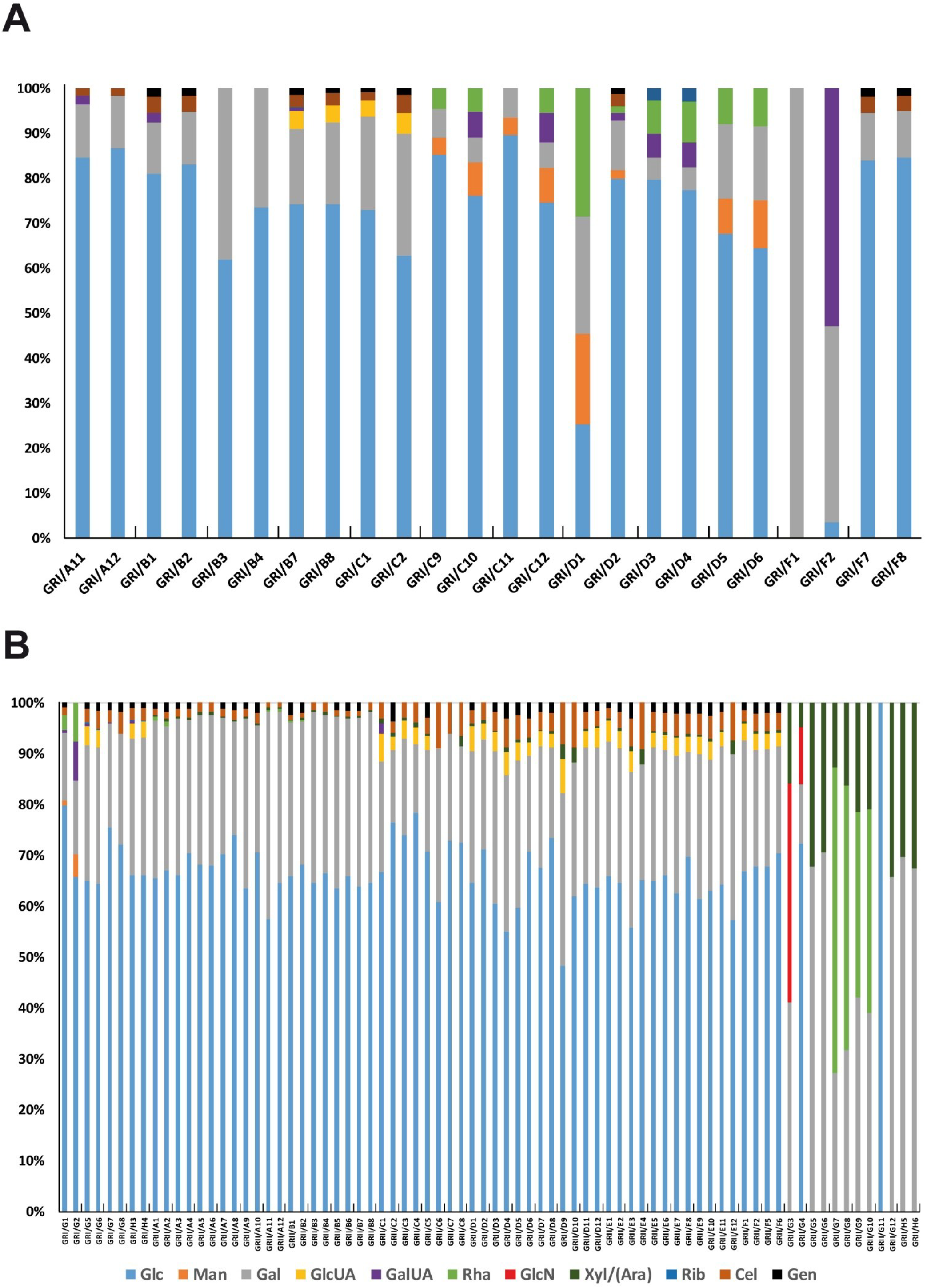
Compositional EPS analysis results from the sugar detection via HT-PMP method of subcollection Reference Strains (A) and subcollection Rhizobacteria Isolates (B), expressed as relative abundance (%). Label indicate the strain position on the plate (Table S3). Abbreviations: Glc, d-glucose; Man, d-mannose; Gal, d-galactose; GlcUA, d-glucuronic acid; GalUA, d-galacturonic acid; Rha, L-rhamnose; Rib, d-ribose; Cel, Cellobiose; Gen, Gentiobiose.

### Subcollection: Rhizobacteria Isolates

This group of 51 root-associated bacteria (GRII and part of GRI plates) coming from different plants, include 41 legume nodule isolates of alfalfa, bean and soybean from Argentina, Spain and China, respectively [27-29], and 10 PGPR *Pseudomonas* from Peppermint rhizosphere of Argentina soils [30]. Overall, the presence of *pleD\** generated discrete quantitative changes on EPS production, with TCC, TCC/OD and TCC/TPC ratios (*pleD*/*WT) of 0.69, 1.94 and 2.26 respectively (Figure S3 and Table 1). However, the presence of *pleD*\* induced strong CR^+^ and CF^+^ phenotypes in several rhizobia isolates: see e.g. GR-03 (Figures S1, G2 *vs*. G1 of GRI plate), GR-42 (Figures S1, G8 *vs*. G7 of GRI plate) or Sfr B33 isolates (Figures S1, C8 *vs*. C7 of GRII plate), likely evidencing the presence of cellulose or other related glucan. In contrast, among PGPR *Pseudomonas* isolates, like *P. spp* SJ07b, the presence of *pleD\** generated a deep impact on the colony morphology displaying wrinkled colonies but without any differential staining in CR or CF, thus suggesting the c-di-GMP promotion of an EPS different from cellulose in those strains (Figure S1, G12 *vs*. G11 of GRII plate).

Thirty-eight of the 51 strains were subjected to the full carbohydrate fingerprint analysis by UHPLC–UV–ESI–MS (Figure 2B). With independence of the c-di-GMP levels, a clear difference between rhizobia and *Pseudomonas* isolates was observed in the composition of their soluble EPS fractions. Glucose was the major component in rhizobia (50-80%) while galactose was most frequent in *Pseudomonas* (Figure 2B). An exception was strain P. spp. SJ08, where the presence of *pleD\** promoted an increase of glucose from none to more than 70% of the total soluble EPS fraction (GRII/G4 *vs*. GRII/G3; Figure 2B). Among the rhizobial strains, there are no significant changes in the composition of their EPS in the presence or absence of *pleD\**. However, a qualitative change is observed in relation to the type of isolate. The presence of glucuronic acid is common in EPS derived from soybean isolates (*Sinorhizobium fredii* from China), while it is not in those from alfalfa plants (*Ensifer meliloti* from Argentina; Figure 2B).

### Subcollection: EPS Environmental Isolates

As detailed above, this group is composed by 182 strains (MTP-I and MTP-II plates) isolated from different environmental habitats [37]. Thirty-six strains were subjected to the full analysis of its monomeric composition by UHPLC–UV–ESI–MS (Figure 3). Overall, the presence of *pleD\** generated discrete quantitative effects on EPS production (Table 1), albeit strong wrinkled phenotypes could be observed in some of the assayed strains (e.g. C6 *vs*. C5 or F10 *vs*. F9 of MTPIIB in Figure S1). This can be partly explained by the fact that *pleD\** generated a widespread negative impact on certain taxonomic groups. For example, different *Xanthomonas* strains displayed TCC ratios <1 (Figure S4 and Table S4, MTP II/H7-H10) which correlated with a reduced mucoid aspect of colonies and changes in the sedimentation pellet generated after their centrifugation (H6 *vs*. H5 or H8 *vs*. H7 of MTPIIB plate in Figure S1 and Figure S5A). The slime reduction induced by *pleD\** in these *Xanthomonas* strains was associated with a decrease in mannose and an increase primarily in fucose and galactose: *Xanthomonas* H7 exhibited a 65% reduction in mannose, alongside a 559% and 161% increase in fucose and galactose, respectively (MTP-IIB/H2 vs. MTP-IIB/H1; Figure 3). Similarly, *Xanthomonas* H9 showed a 67% reduction in mannose, with a 161% and 141% increase in fucose and galactose, respectively (MTP-IIB/H6 vs. MTP-IIB/H5; Figure 3). On the other hand, the presence of high levels of c-di-GMP in this group of bacteria caused significant qualitative changes in the composition of their respective EPS, with the appearance of new sugars such as fucose, glucuronic acid, and rhamnose in many of them (Figure 3).

**Figure 3:**
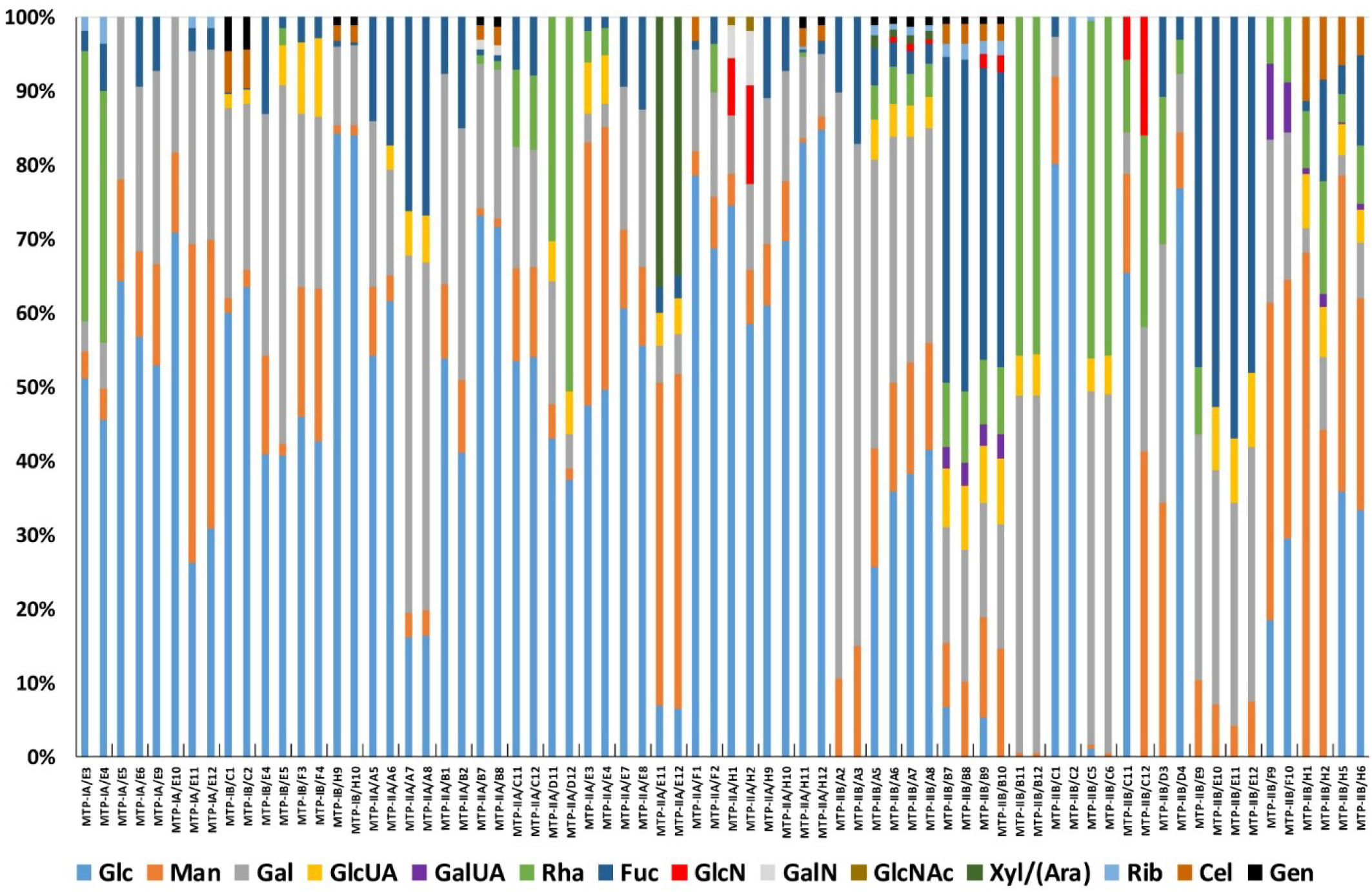
Compositional EPS analysis results from the sugar detection via HT-PMP method of subcollection Environmental Isolates expressed as relative abundance (%). Label indicate the strain position on the plate (Table S3). Abbreviations: Glc, d-glucose; Man, d-mannose; Gal, d-galactose; GlcUA, d-glucuronic acid; GalUA, d-galacturonic acid; Rha, l-rhamnose; Fuc, L-fucose; GlcN, d-glucosamine; GalN, d-glalactosamine; GlcNAc, *N*-acetyl-d-glucosamine; Xyl/(Ara) Xylose/arabinose; Rib, d-ribose; Cel, Cellobiose; Gen, Gentiobiose.

### Subcollection: *Sphingomonas* strains

This turned out to be one of the most interesting groups, where the presence of *pleD*\* had a strong impact on the quantity of soluble EPS with an average TCC ratio (*pleD\**/Vect) = 1.48 (Table 1 and Figure S6). This ratio was even higher when samples were standardised with the optical density (OD) or the total protein content (TPC) of their cultures: 3.69 and 9.22, respectively (Table 1). Indeed, among the 330 assayed strains, three *Sphingomonas* (SphC10, SphD3 and SphE3) were in the top 3 in terms of their TCC/TPC ratio levels (Table S4). The overexpression of *pleD*\* caused a deep impact on the colony morphotype with pronounced wrinkles in some strains: see e.g. E4 *vs*. E3 or H2 *vs*. H1 in SPH plate (Figure S1). Interestingly, contrary to what was observed in other groups, the presence of *pleD*\* in Sphingomonads was not associated with positive interaction with any of the dyes tested (Figure S1), arguing about the possibility of different EPS involved. The impact was even stronger in liquid media, with formation of robust *pleD*\*-dependent biofilms over the walls of the DWP: see e.g. E8 and G8 *vs*. E7 and G7 of SPH plate, respectively (Figure S5B). Furthermore, a strong flocculation associated with the presence of *pleD*\* was also observed (Figure S5B).

From the 34 analysed strains, 15 of them were selected to determine their EPS composition by the HT-PMP method via UHPLC-UV-ESI-MS [37]. Despite some differences concerning uronic acids (GlcUA and GalUA), in most cases the composition of the c-di-GMP induced EPS were similar to their respective controls, arguing for quantitative rather than qualitative changes on EPS production induced by c-di-GMP in this bacterial group (Figure 4).

**Figure 4:**
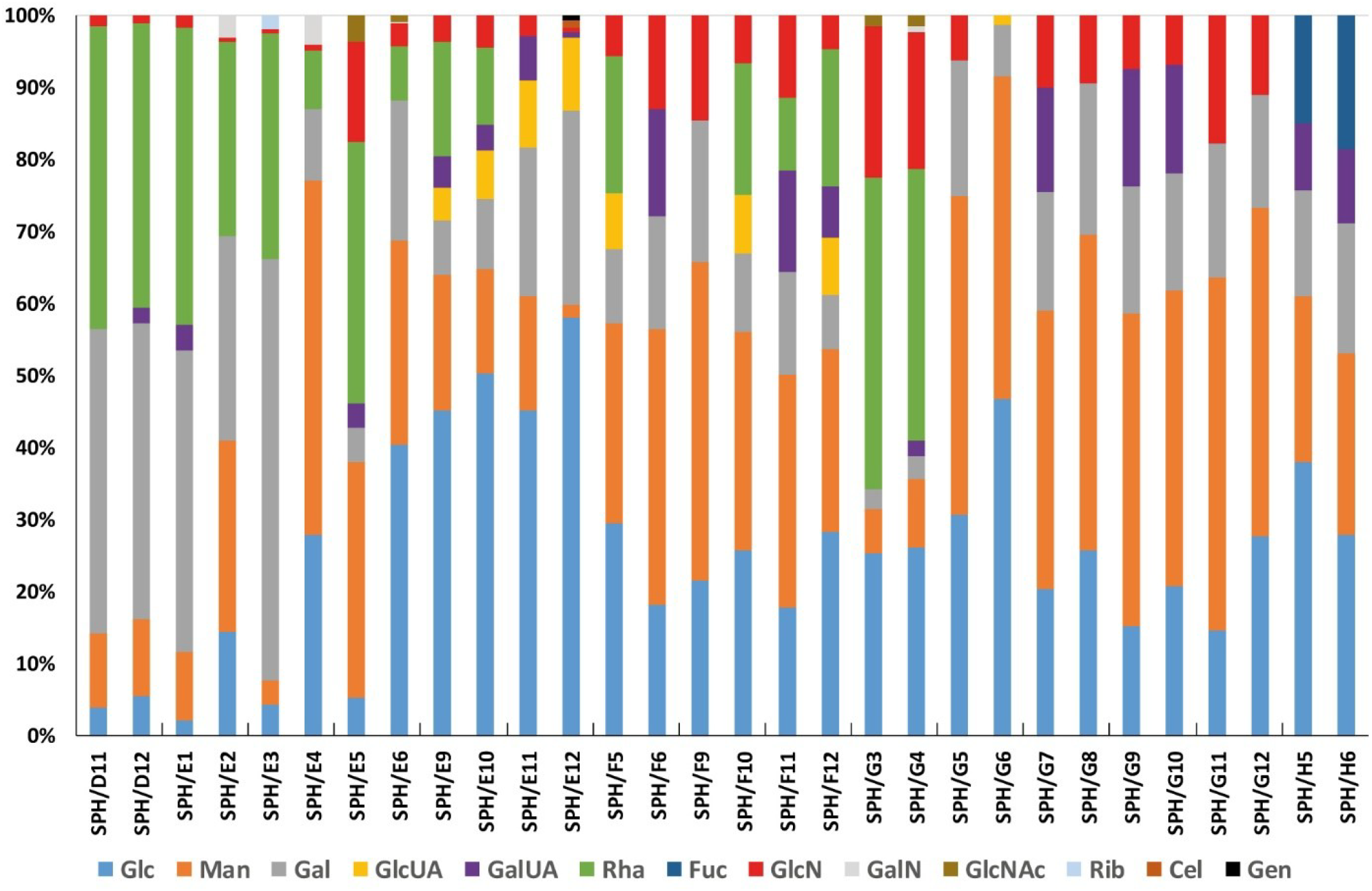
Compositional EPS analysis results from the sugar detection via HT-PMP method of subcollection Sphingomonas Strains expressed as relative abundance (%). Label indicate the strain position on the plate (Table S3). Abbreviations: Glc, d-glucose; Man, d-mannose; Gal, d-galactose; GlcUA, d-glucuronic acid; GalUA, d-galacturonic acid; Rha, L-rhamnose; Fuc, l-fucose; GlcN, d-glucosamine; GalN, d-glalactosamine; GlcNAc, *N*-acetyl-D-glucosamine; Rib, d-ribose; Cel, Cellobiose; Gen, Gentiobiose;

### SphC10 strain produces a novel c-di-GMP-promoted EPS with an unusual sugar composition

Among all the strains screened, SphC10 was selected for further characterization due to different reasons: (i) It was the strain displaying the highest TCC/TPC ratio (pleD*/WT) among all assayed strains = 133 (Figure 1 and Table S4), (ii) the presence of *pleD*\* had a strong impact on its growth, both in solid and liquid media (Figure S7) and (iii) the soluble EPS induced by c-di-GMP has an unusual sugar composition, i.e. presence of galactose (SPH/E12 in Figure 4), which is not common among previously described sphingans [39]. Genomic DNA of SphC10 was extracted and its 16S gene partially sequenced, which showed 100% homology with different strains belonging to *Sphingomonas* genus, including *Sphingomonas yabuuchiae* strain E178 [48] and other *Sphingomonas* sp. strains.

The pJBpleD* and the empty vector pJB3Tc19, were individually reintroduced into the original SphC10 strain. Characteristic donut-shape colonies were formed by SphC10 pJBpleD*, in contrast to the smooth regular colonies produced by the control SphC10 pJB3Tc19 (Figure S7B). A liquid culture in EPS medium set up from a single colony of SphC10 pJBpleD* showed a thick biofilm attached to the flask walls after 48h, in contrast to the vector control (Figure S7C). Samples for c-di-GMP quantification and Scanning Electron Microscopy (SEM) were collected in triplicates from both cultures. The over-expression of *pleD\** generated a strong increase in the intracellular c-di-GMP levels by ≈40 fold in SphC10 pJBpleD* (75.8 ± 3.5 ng of c-di-GMP/mg protein), compared to the control strain containing the empty vector pJB3Tc19 (1.9 ± 0.7 ng of c-di-GMP/mg protein). Microscopy studies revealed that, in contrast to the control, SphC10 pJBpleD* cells grow forming compact flocs where the cells are wrapped in a dense extracellular matrix (Figure 5). After centrifugation, SphC10 pJBpleD* EPS was precipitated from the culture supernatants, centrifuged and desiccated. The monosaccharide composition was this time determined by GC-MS through the formation of per-trimethylsilyl methyl glycosides [49]. Given the presence of uronic acids and the resilience of their glycosidic bonds to hydrolysis, a previous treatment with TFA 2 M was applied. The EPS fraction precipitated was mainly formed by Glc, Gal, GlcUA and Man, in a ratio of 100:33:4:2, respectively. These results closely align with the HT-PMP derivatization analysis obtained from supernatant fraction of the HTS, which identified Glc, Gal, GlcUA and Man as main constituents in a 100:47:17:3 ratio, respectively. The different ratios of GlcUA can be attributed to variations in the degradation rate of the uronic acids under the hydrolysis conditions used [37].

**Figure 5.**
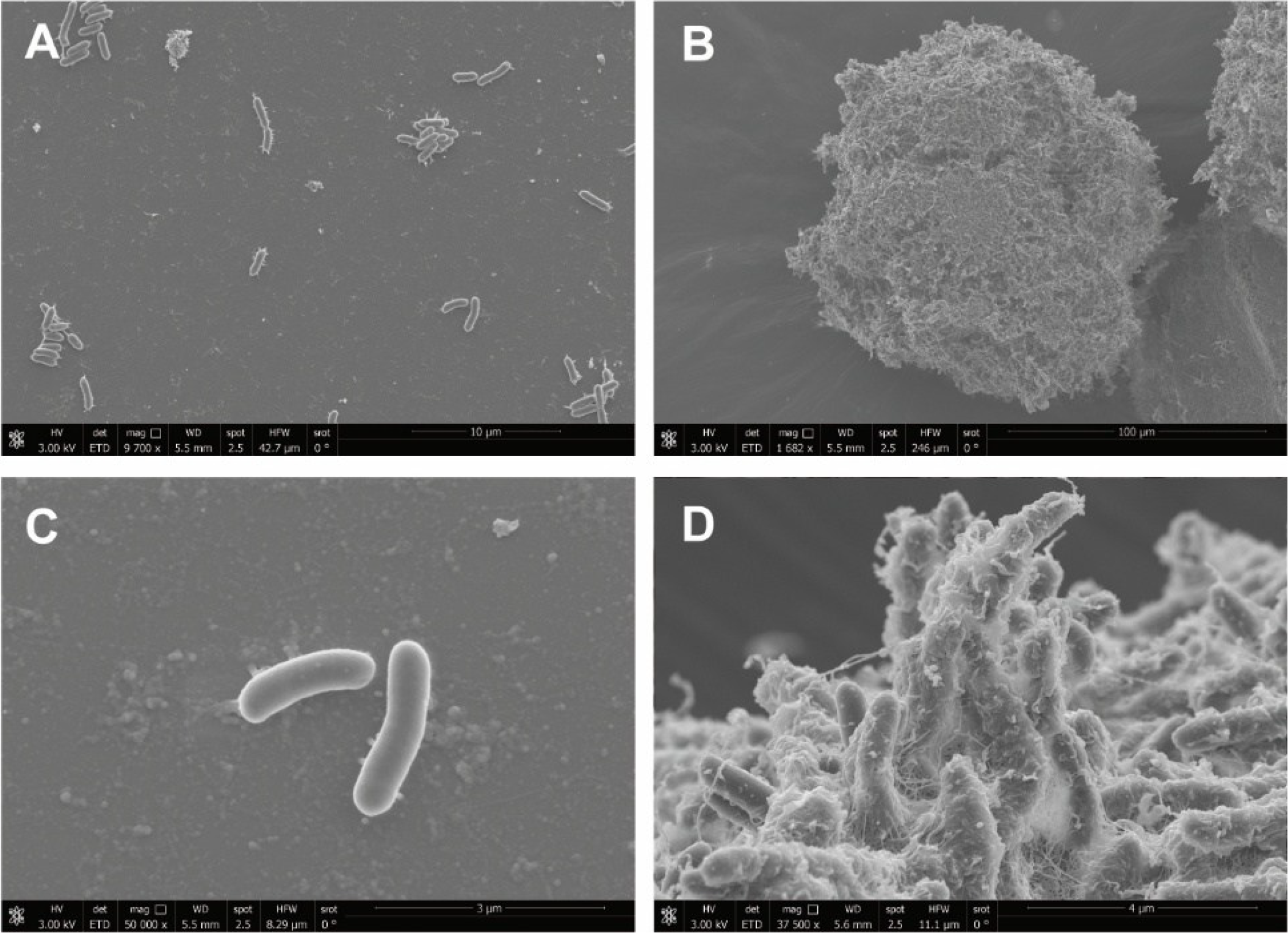
Scanning electron microscopy (SEM) images of SphC10 pJBpleD* at ×1,682 (B) and at ×37,500 (D) and the control SphC10 pJB3Tc19 at ×9,700 (A) and at ×50,000 (C). The images reveal details of the mesh of entangled filaments formed in flocs of SphC10 with high c-di-GMP levels (pJBpleD*).

### Characterization of the c-di-GMP-promoted EPS of SphC10

In summary, our screening revealed the presence of an EPS in the culture supernatants of SphC10 strain whose production was greatly induced by high intracellular levels of c-di-GMP. On the contrary, in microscopy studies, the observed flocculation and the increased volume of the SphC10 pJBpleD* pellets after centrifugation suggested the presence of a cell-associated EPS. To shed light on this, a time course of EPS production was carried out in triplicate flasks with 50 mL EPS media, collecting samples of 15 mL at 24, 48 and 72 hours. Supernatants were separated from cells by centrifugation and the pellets were weighed. An average differential increment of a pellet weight of 213% was observed in the presence of *pleD\** at 24 h, reaching a 2,000% at 48 h (Figure 6). Pellets were subsequently processed following a mild extraction protocol (see M&M). Nearly all of this weight increment disappeared after the extraction was applied, suggesting that most of the c-di-GMP-promoted EPS was detached from the cells and released to the supernatant (Figure 6). EPS from the extracted fractions at 72 h were precipitated, redissolved in water and quantified by the total sugar content, based on absorbance measurement after a phenol-sulfuric-acid treatment. EPS from SphC10 pJBpleD* sample showed 15.1 ± 0.7 mg of glucose equivalents achieved from the 15 mL (1.01 g/L) in contrast to the 0.03 ± 0.002 mg obtained from the 15 mL (0.002 g/L) of SphC10 with the control vector. The composition of the SphC10 pJBpleD* EPS extracted and precipitated (EPS-2) was determined as described before by GC-MS through the formation of per-trimethylsilyl methyl glycosides [49], and compared to the one obtained from the supernatant in the first instance (EPS-1). EPS-2 displays a similar monomeric composition and is manly formed by Glc, Gal, GlcUA and Man, with 100:30:5:2 area ratios, respectively. In addition, the FTIR and ^1^H-NMR spectra of both samples are almost identical (Figure S8), indicating that both polysaccharides are the same.

**Figure 6.**
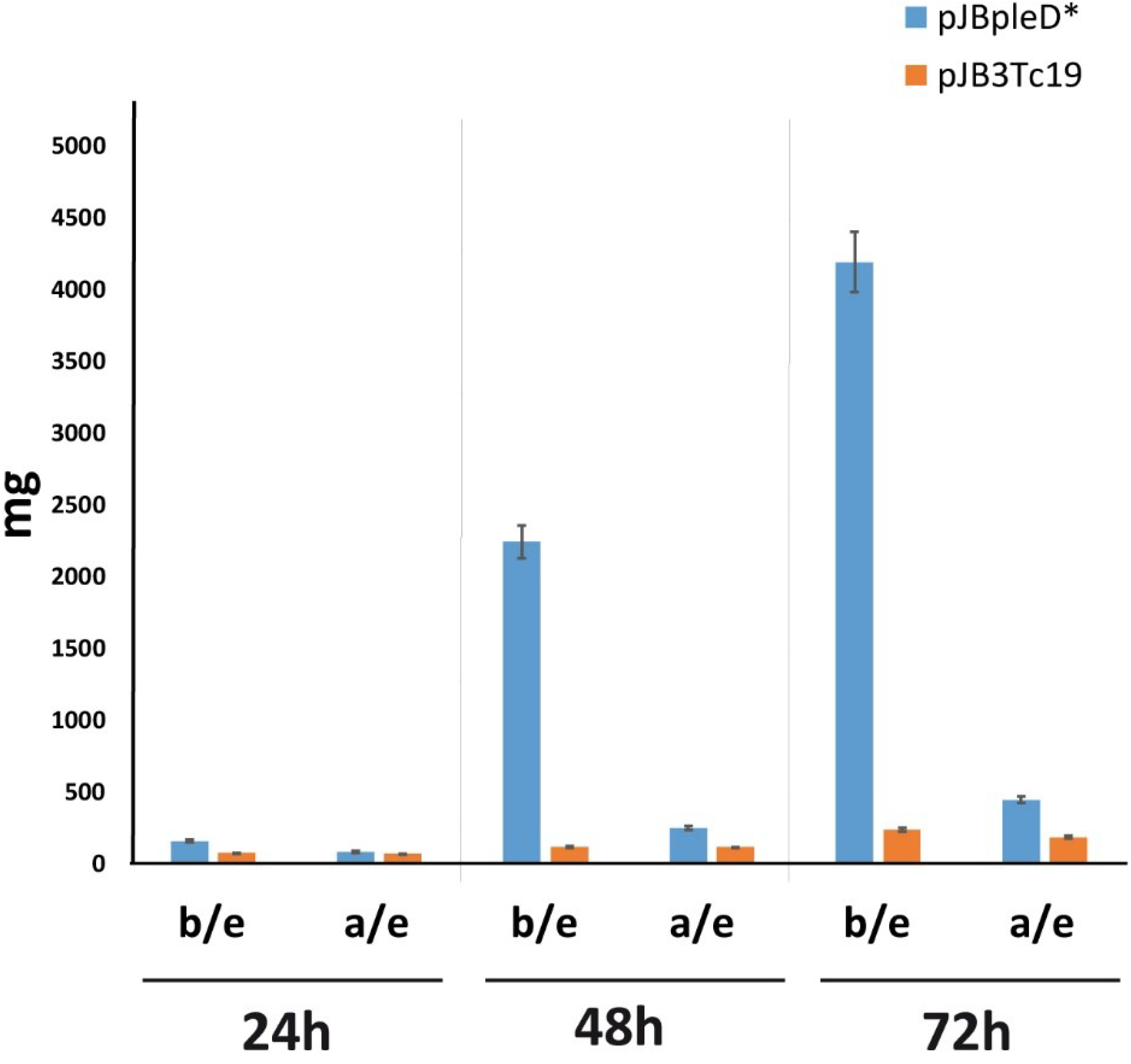
Pellet weight time-course of SphC10 in presence (pJBpleD*, blue bars) or absence (pJB3Tc19, orange bars) of *pleD\**. Supernatants from 15 mL of SphC10 pJBpleD* and SphC10 pJB3Tc19 cultures were separated from cells by centrifugation and their wet pellet were weighed at 24h, 48 h and 72 h before (b/e) and after (a/e) an extraction protocol (boiling remaining pellet in 0,85% of NaCl in distilled water 10 min, plus 5 min of sonication in a bath at 60 °C, followed by centrifugation) was applied.

These results suggest that the second messenger c-di-GMP promoted the production and secretion of an EPS that is loosely associated to the cell surface. As a result, the subsequent study was carried out exclusively on EPS-1. The absolute configuration of its monomers was determined by the formation of their per-trimethylsilylated 2-butyl glycosides [50], indicating that all of them are d-sugars. The glycosidic bond locations were also determined by methylation analysis [51]. The structures of the resulting partially methylated and acetylated alditols (PMMAs), together with a comparison of their retention times with those of standards [52], indicate that the main components are: terminal d-Glc*p*, →4)-d-Glc*p*, →4,6)-d-Glc*p*, and terminal d-Gal*p*, in an approximate ratio of 1:4:1:2, respectively.

The ^1^H-RMN of EPS-1 reveals the presence of a relatively complex polysaccharide, given the number of anomeric signals (between 5.5 and 4.5 ppm; Figure S8). The majority of these anomeric signals are below 5.0 ppm, indicating that the glycosidic linkages are beta in most cases. On the other hand, the signals at 2.2 and 2.1 ppm, which are characteristic of acetyl groups, indicate that the polysaccharide is partially acetylated. Finally, the broad signals at approximately 1.2 and 0.9 ppm probably correspond to fatty acids, being more abundant in EPS-2, which could correspond to remnants of membrane lipids (Figure S8).

## Discussion

The search for natural products is at the pillars of biotechnology, and exploration of microbial diversity is a major approach for discovering new compounds. However, when searching for new products key decisions are where to look as well as how to look. Bacteria represent a vast reservoir of biodiversity, thriving in almost all habitats. These microorganisms are abundant in those capable of producing biomolecules, particularly exopolysaccharides (EPS), which possess unique physical properties such as monosaccharide composition, structural conformation, molecular weight and functional groups, finding numerous applications across a wide range of industrial sectors. They are still constrained by production costs and capacity when considered for bulk markets. However, in recent years high-value markets for specific applications have arisen, where bacterial EPS can be successfully commercialized due to their unique physicochemical and/or bioactive properties.

EPS are produced by microorganisms in response to biotic or abiotic stress factors, providing additional biological protection for the cells. Thus, under stable conditions typically found in laboratory or industrial fermentation processes where the inducing environmental cues are absent, their biosynthesis is usually repressed. This supports the notion that the natural diversity of bacterial EPS remains largely unexplored [10]. As the dinucleotide cyclic di-GMP (c-di-GMP) has emerged as a universal second messenger in bacteria that controls, among other functions, the production of diverse EPS in a limited number of species, we hypothesized that the manipulation of c-di-GMP economy in large bacterial collections can reveal the production of EPS that would otherwise be cryptic under standard culture conditions.

We carried out a protocol for high throughput automated conjugation to introduce and express the DGC PleD* into hundreds of bacterial strains [35, 36], in order to increase their c-di-GMP intracellular levels. 83% of the strains could be transformed with the pJBpleD* plasmid, supporting the use of this HTS approach for the rapid transformation of virtually any large bacterial collection or environmental isolates.

As expected, the presence of *pleD\** led to a generalized increase in the total carbohydrate contents (TCC) of the culture supernatants. Nearly 60% of the strains tested showed a *pleD*\*/WT ratio of TCC/TPC >1, with average increases of 2.76 fold and with 7 strains showing EPS increments above 10-fold. Considering that one of the unavoidable steps in the analysis implies the complete removal of the insoluble cell-associated EPS (e.g. cellulose), the actual positive impact of *pleD\** on EPS production is probably higher. These results fully agree with previous studies establishing c-di-GMP as a general activator of EPS production in bacteria [10, 11, 16, 53]. The presence of *pleD\** not only quantitatively impacted EPS production across different bacterial groups but also qualitatively altered their composition, as indicated by the emergence of new monosaccharides that were not detected under physiological c-di-GMP conditions. Our results support the artificial increase of intracellular c-di-GMP levels as a reliable approach to unveil novel EPS, by deregulating their biosynthesis and promoting their production under laboratory conditions. By that, this approach that can be adapted to analyse large bacterial collections to identify new carbohydrate polymers with multiple purposes: the discovery of novel polysaccharides with potential biotech applications, enhancing our knowledge on the biological diversity of carbohydrate polymers and their biosynthetic pathways, and expanding the molecular and functional targets of c-di-GMP regulation in bacteria. Our study was able to identify a significant number of strains (near 10 %) as promising candidates to produce novel EPS regulated by c-di-GMP.

As a proof of concept, in this study we have uncovered and partially characterized a novel *Sphingomonas* EPS. The presence of the plasmid pJBpleD* led *Sphingomonas* C10 strain to hyperproduce an EPS that is otherwise cryptic. Two different monomeric determination analysis carried out in this study showed glucose, galactose, glucuronic acid and mannose as the main components of this EPS. Although the complete characterization of the SphC10 EPS remains elusive, our analysis suggests a complex polysaccharide structure, with predominant beta-glycosidic bonds and partially acetylated. The presence of galactose and absence of rhamnose in this c-di-GMP-regulated EPS is unusual and, to our knowledge, does not align with any previously reported sphingans. The sphingan family, including biotechnological relevant EPS such as gellan, welan, diutan, conform a closely related group of secreted acid polysaccharides characterized by similar chemical structures. Their main components include a tetrasaccharide building block of glucose, rhamnose, mannose and glucuronic acid. These components share nearly identical backbone structures, consisting of the repeating unit α-l-Rha*p*-(1→4)-β-d-Glc*p*- (1→4)-β-d-Glc*p*UA-(1→4)-β-d-Glc*p*-(1→ [38].

Interestingly, although our strategy would in principle rule out insoluble EPS associated with the cellular fraction, the SphC10 polymer appears to be an EPS weakly associated with the membrane of the producing cells. Our results indicate that under high c-di-GMP conditions, SphC10 massively produces this EPS that keeps cells wrapped in a thick biofilm forming large flocs when grown in liquid medium. Part of this EPS loosely attached to the cells is released into the medium during growth, which made its identification possible during the HTS. This EPS can be released almost entirely from the cellular fraction by physical means, such as vortex combined with heating and/or sonication. FTIR studies and monomeric composition supported that both attached and detached EPS (EPS-1 and EPS-2) are the same.

Although the exact function of sphingans in producing bacteria is not completely clear, they play an important role in cellular protection by forming a protective layer around bacterial cells, helping them to form biofilms and withstand adverse environmental conditions [54]. The second messenger c-di-GMP is a universal biofilm activator in a plethora of bacteria [55, 56]. However, the information about c-di-GMP signalling in *Sphingomonas* is very limited. To the best of our knowledge, only two studies have linked c-di-GMP with EPS production and biofilm formation in *Sphingomonas sp*. Vorholt’s lab described the transcriptional regulator *ctrA* regulates c-di-GMP signalling and EPS synthesis genes in *Sphingomonas melonis* [57]. Li and co-workers recently showed evidence that c-di-GMP is involved in regulation of welan gum biosynthesis in *Sphingomonas* sp. WG [58]. These studies suggest that sphingans may be positively regulated by c-di-GMP at the transcriptional level. The overall pathway of sphingan biosynthesis follows the Wzx/Wzy-dependent route, as described for various capsular polysaccharides in other bacteria [38, 59]. Sequencing the genome of the SphC10 strain along the identification of the biosynthetic and regulatory pathways for this EPS will be a focus of future investigations. Another important research focus will be the elucidation of the EPS structure and the study of its physic-chemical and rheological properties, which can advance its potential biotechnological applications.

### Concluding remarks

The biological diversity found in bacteria represents an excellent resource for the discovery of novel bacterial polymers such as exopolysaccharides (EPS). In the context of a more sustainable future and a more efficient circular economy, bacterial EPS are well-positioned to replace fossil-based materials with sustainable, bio-based alternatives, transforming low-cost carbon and nitrogen sources (e.g., industrial effluents) into high-value products. However, realizing this potential requires efficient, rapid, and reliable methods for identifying microorganisms capable of producing diverse and novel EPS in significant quantities. EPS are valuable to the bacteria that produce them, often being synthesized in response to biotic or abiotic stress. As a result, their production is typically repressed under stable industrial fermentation conditions where such environmental cues are absent. This suggests that the natural diversity of bacterial biopolymers remains largely untapped.

In this study, we present a tailored strategy to search for, characterize, and specifically manipulate the signal transduction pathways involved in EPS repression. We apply this approach as a rational method to unlock the hidden diversity of bacterial polymers by deregulating their biosynthesis and promoting large-scale production under laboratory conditions. Our findings indicate that the universality of the second messenger c-di-GMP makes it an ideal candidate for biotechnological applications as a universal activator of novel bacterial polymers.

### Outstanding questions

- In light of the extensive biosynthetic potential unveiled in bacteria in the post-genomic era, how many bacterial polymers with biotechnological relevance— comparable, for instance, to that of bacterial cellulose—remain to be discovered?
- To what extent can targeted, *ad-hoc* modifications of signal transduction pathways serve as an effective strategy to unveil this hidden polymeric diversity under controlled laboratory conditions?

### Technology readiness

The present study covers the initial stages of technological development, including proof of concept and its validation at the laboratory level (TRL 1–3). The results obtained, however, suggest the potential for implementation in later stages of technological development (TRL 4–7), enabling the industrial-scale overproduction of various novel EPS of biotechnological interest.

## Acknowledgements

DPM was supported by Research Stays for University Academics and Scientists DAAD fellowship (2018). We thank Dr. Mareike Wenning (formerly Technical University of Munich) for providing the 62 *Sphigomonas* strains used in plate SPH in the study. The authors would like to thank the University’s NMR Facility (CITIUS) for their technical support and Ana M. Matia González is acknowledged for her support with the figures. This work was supported by the European Union Next Generation EU/PRTR project TED2021-129640B-I00 and grants BIO2017-83533-P and PID2022-140168NB-I00 by MCIN/AEI/10.13039/501100011033 and by “ERDF A way of making Europe”

## Author contributions

DPM, JS, JSP: conceptualization, investigation. DPM, MD, BR, MARC, MBM: validation. JSP, VS: supervision. DPM: writing – original draft. JS, BR, MARC, JSP: writing – review and editing.

## Materials and Methods

### Culture Media and Growth Conditions

Bacterial strains used in the HTS are listed in Table S1. The strains were stored at −80 °C in a 96-well micro titer plate in their respective media containing 20% of glycerol. Precultures were grown in a 96-deep-well plate (DWP) containing 1 mL of their respective culture media. Inoculation was performed with a 96-pin replicator. After incubation of the preculture for 48 h in their respective temperatures in a microplate shaker (1000 rpm) equipped with an incubator hood (TiMix 5 control and TH 15, Edmund Bühler GmbH), the main culture was inoculated with 10 µL of the preculture in 990 µL of fresh their respective screening media and incubated in the same way as described above. Strains already transformed from GRI and GRII plates were grown at 28 °C in a Minimal Media (MM) previously reported to be able to sustain growth of different rhizobacteria [41] and were subjected directly to the HTS.

Strains from MTP-IA &-IB, MTP-IIA &-IIB and SPH plates were inoculated from backup plates with a replicator into two separated DWP containing EPS media [26] and conjugated by using a handing-liquid station with *E. coli* β2163 donor strains harbouring the pJBpleD* [23] or the empty vector pJB3Tc19 [60], respectively grown up to an exponential phase (OD_600nm_ = 0.6) in LB broth [Luria-Bertani broth for 1 L: 10 g tryptone, 5 g yeast extract, 5 g NaCl supplemented with Tc at 10 µg/mL and diaminopimelate (DAPA) to a final concentration of 0.3 mM]. Resuspension of the matings were inoculated into separated DWP and grown on EPS selective media supplemented with tetracycline (Tc) for 72 h. Two rounds of dilutions and culturing of the strains on EPS media + Tc selective marker were carried out to ensure the enrichment of the transconjugants. Backup plates of transconjugants were generated and stored before subsequent analyses. Detailed information about the automated conjugation process can be found in Schmid *et al*. [35].

### High-Throughput Screening (HTS) analysis of the bacterial strains

The analysis carried out during the HTS was described in detail by Rühmann *et al*. giving further hints for data interpretation [37, 40]. Prior to the analysis itself, transconjugants were inoculated with a 96-pin replicator and grown 72 h at 30 °C on selective solid medium independently supplemented with 3 different dyes: Congo Red (CR) Calcofluor (CF) and Aniline Blue (AB) at 50, 200 and 50 µg/mL, respectively. The different growth steps, both in solid and liquid medium, were documented through pictures taken with a regular camera.

### Total Carbohydrate Content (TCC)

The supernatant of all assayed strains were subjected to a fully automated fast detection method of the total sugar content based on absorbance measurement after a phenol-sulphuric-acid treatment in a 96-well microplates format [40]. To minimize the effect of the different bacterial growing rates on the amounts of EPS production, the TCC was standardised by two different parameters: (i) the optical density (OD_600nm_) of the culture (OD) and (ii) the Total Protein Content (TPC) of the cell pellets. For TPC calculation, the cellular pellets generated during the HTS after the DWP centrifugation were suspended in 900 µL of dH_2_O. 100 µl of Triton X-100 (0,25%) was added and samples boiled during 15 min. 25 µL of sample was mixed with 25 of dH_2_O and 100 µL of Bradford reagent (Bio-Rad) and total protein calculated according to manufacture instructions.

### UHPLC-UV-ESI-MS Analysis

Monosaccharide compositions were determined by a previously developed HT-PMP derivatization method for carbohydrate analysis with UHPLC-UV-ESI-MS/MS [61]. In brief, fermentation supernatants were diluted to about 1 g/L and filtrated. Hydrolysis was performed by incubating 20 μL EPS solution with 20 μL 4 M TFA at 121 °C for 90 min in a 96-well PCR plate. After cooling to room temperature, hydrolysis was stopped by adding 3.2% NH_4_OH solution, increasing the pH to slightly alkaline conditions. Sugar standards were prepared with the neutralized TFA-hydrolysis matrix. Derivatization of samples/standards was performed by mixing the 25 μL hydrolysate with the 75 μL derivatization reagent, containing a mix of 0.1 M PMP in MeOH and 0.4% NH_4_OH solution in ratio 2:1 (v/v). After incubation for 100 min at 70 °C in a PCR-cycler 20 μL derivatized sample was mixed with 130 μL 19.23 mM acetic acid. Samples were filtered into a microtiter plate, which was sealed with a silicon cap mat. Analysis was performed using UHPLC equipped with a reverse phase column tempered to 50 °C and a UV detector (245 nm). Mobile phase A contained a mixture of 5 mM ammonium acetate buffer (pH 5.6) with 15% (w/w) acetonitrile and mobile phase B contained pure acetonitrile following the programmed gradient at a flow rate of 0.6 mL min^−1^. Before accessing ESI-MS, the flow was split 1:20.

### GC-MS

Monosaccharides were analysed by GC-MS separation of their per-*O*-trimethylsilylated *O*-methyl glycosides following the method by Chaplin [49]. Briefly, samples containing polysaccharide (200 µg) together with *myo*-inositol (internal standard, 2 µg) were treated with 1.25 M Hydrochloric acid in methanol at 80 °C for 16 h. After removal of solvent under nitrogen stream, traces of HCl were eliminated by co-evaporation with *tert*-butanol. Trimethylsilylation was carried out by treatment with a mixture of 100 µL of pyridine and 100 µL of *N,O*-bis(trimethylsilyl)trifluoroacetamide (BSTFA, Sigma-Aldrich) for 2 h at room temperature. Finally, samples were evaporated, dissolved in hexane and analysed by GC-MS. A modification of this method, with a previous hydrolysis with 2 M trifluoroacetic acid (TFA) (120 °C, 2 h) followed by evaporation under nitrogen stream and removal of TFA traces with methanol, was also applied. In this case, the methanolysis with HCl/MeOH was carried out for 2 h.

The absolute configuration of each sugar moiety was established through the analysis of the trimethylsilyl *O*-2-butyl glycoside derivatives, as previously reported [50]. The location of glycosidic bonds was determined by formation of partially methylated and acetylated alditols (PMAA) followed by GC–MS analysis [51]. All the GC-MS spectra were acquired on an Agilent 8890GC coupled to an Agilent 5977B GC/MSD equipped with a column Agilent J&W HP-5ms Ultrainert (30 m × 250 µm × 0.25 µm) and He as carrier gas.

### Spectroscopy Methods

ATM-FTIR spectra were acquired on a Jasco FT/IR-4100 system.

^1^H-NMR spectra were recorded in D_2_O at 353 K on a Bruker Avance NEO 500 spectrometer operating a 500.20 MHz, using the residual HDO as reference (δ_H_ 4.22 ppm). Spectral width of 20 ppm was digitized using 32k data points, with 32 scans per spectrum.

### SphC10 Intracellular c-di-GMP quantification

Three biological replicates of SphC10 pJB3Tc19 and pJBpleD* strains were individually grown in flasks with 50 mL of EPS broth inoculated from starting cultures at a final OD_600nm_ = 0.01.

The cells were collected by centrifugation (10,000 rpm) at an OD_600nm_ = 0.6, using the cells from 10 mL of the culture for the extraction of c-di-GMP and other 10 mL for protein quantification via Bradford [62]. Cyclic-di-GMP was extracted using a protocol based on a previous report [23]. Quantification of the c-di-GMP levels of the samples were carried out using an ELISA assay from Cayman^™^ commercial kit (Item No. 501780), following the instruction recommendations including positive and negative controls. Intracellular c-di-GMP levels of each strain were standardized by the amount of total protein content and expressed as the average ± standard deviation of pg of c-di-GMP/mg protein.

### Scanning electron microscopy (SEM)

A High Resolution Field Emission Environmental Scanning Electron Microscope (ESEM) with QuemScan and FEI Cryo Station, mod. QuemScan650F was used. Samples of cells and flocs from liquid cultures in EPS media of *Sphingomonas* SphC10 strain expressing *pleD*\* (by use of pJBpleD*) or not (with pJB3Tc19) were processed and imaged by the technical services of the University of Granada CIC-UGR (https://cic.ugr.es/servicios/servicios-unidades/microscopia/microscopia-electronica-barrido-ambiental-esem).

### Analysis of 16S rRNA of SphC10 strain

Genomic DNA was extracted from stationary phase cultures using a Durviz^™^ RealPure kit. PCR amplified band of 16S rDNA gene using primers: 41F (5′-GCTCAAGATTGAACGCTGGCG-3′) and 1488R (5′-CGGTTACCTTGTTACGACTTCACC-3′) and performed basically as described previously by Herrera-Cervera *et al* [27]. The DNA fragment obtained was purified and sent for Sanger sequencing. Sequence similarity searches were carried out with the BLASTN program from the National Center for Biotechnology Information [63].

### SphC10 EPS Isolation

Starting cultures of SphC10 strains were diluted 1:100 in 500 mL flasks containing 50 mL of EPS broth supplemented with Tc and were grown with shaking (120 rpm) 24, 48 and 72 h at 30 °C. Cell pellets were recovered by centrifugation in Falcon tubes (Fisher Scientific) for 15 minutes at 4,000 rpm and the supernatant fraction was separated in a new tube. The EPS of this fraction was precipitated by adding 2 volumes of ethanol and leaving it overnight at 4 °C. The EPS obtained (EPS-1) was dried, frozen and lyophilized. Cell pellets were resuspended in 5 mL of 0,85% of NaCl in distilled water by vortex and submitted to an extraction protocol consisting in boiling it for 10 minutes followed by sonication in a bath at 60 °C for 5 minutes. Samples were centrifuged in Falcon tubes (Fisher Scientific) for 15 min at 4,000 rpm and supernatant fraction was separated in a new tube. The EPS of this cellular associated fraction (EPS-2) was precipitated dried, frozen and lyophilized as described above.

## Supplemental information titles and legends

**Table S1:** Bacterial strains analysed in this study. Relevant characteristics and plate origin are indicated.

**Table S2:** Deep Well Plates (DWP) processed for conjugation. Number of strains used and transconjugants obtained in this study.

**Table S3:** Plate location of transformed strains with pJBpleD* (even columns) and with the empty vector pJB3Tc19 (odd columns). Optical density (OD_600nm_) of the cultures and (ii) the Total Protein Content (TPC) of the cell pellets of each strain are included and the standardisation of TCC values by OD (TCC/OD) and TPC (TCC/TPC) are calculated. Blanks as negative controls at different positions are indicated.

**Table S4:** Total Carbohydrate Content (TCC) ratios of the *pleD*\* *vs*. the empty vector culture supernatants of each strain. TCC: TCC Ratios *pleD\**/WT; TCC/OD: TCC ratios *pleD\**/WT normalised with the OD_600nm_; TCC/TPC: TCC ratios *pleD\**/WT normalised with the Total Protein Content.

**Figure S1:** Phenotype of the different strains with pJBpleD* (even columns) and with the empty vector pJB3Tc19 (odd columns) after grown 72 hours in EPS solid media supplemented with 3 different dyes: Congo Red (CR), Aniline Blue (AB) and Calcofluor (CF). Plates with CF were imaged under UV light. Square plates containing the solid media with the different dyes were inoculated with 96-well replicator from transconjugants backup plates. Squares of the same colour indicate pairs (empty vector *vs. pleD\**) of relevant strains cited in the text.

**Figure S2:** Impact of c-di-GMP over the EPS production of subcollection Reference Strains. Representation the Total Carbohydrate Content (TCC) ratios of the *pleD*\* *vs*. the empty vector culture supernatants of each strain (Table S4). TCC Ratios PleD*/WT (in blue); TCC ratios PleD*/WT normalised with the OD_600nm_ (TCC/OD; in black); TCC ratios PleD*/WT normalised with the Total Protein Content (TCC/TPC; in orange) are shown.

**Figure S3:** Impact of c-di-GMP over the EPS production of subcollection Rhizobacteria Isolates. Representation the Total Carbohydrate Content (TCC) ratios of the *pleD*\* *vs*. the empty vector culture supernatants of each strain (Table S4). TCC Ratios PleD*/WT (in blue); TCC ratios PleD*/WT normalised with the OD_600nm_ (TCC/OD; in black); TCC ratios PleD*/WT normalised with the Total Protein Content (TCC/TPC; in orange) are shown.

**Figure S4:** Impact of c-di-GMP over the EPS production of subcollection Environmental Isolates. Representation the Total Carbohydrate Content (TCC) ratios of the *pleD*\* *vs* the empty vector culture supernatants of each strain (Table S4). TCC Ratios PleD*/WT (in blue); TCC ratios PleD*/WT normalised with the OD_600nm_ (TCC/OD; in black); TCC ratios PleD*/WT normalised with the Total Protein Content (TCC/TPC; in orange) are shown.

**Figure S5:** Phenotype of different strains after grown in Deep Well Plates (DWP) with EPS liquid media for 48 hours before and after centrifugation of the DWP. A) Two different *Xanthomonas* strains from MTP IIB plate with pJBpleD* (H6 and H8) and with the empty vector pJB3Tc19 (H5 and H7). B) Two different *Sphingomonas* strains from SPH plate with pJBpleD* (E8 and G8) and with the empty vector pJB3Tc19 (E7 and G7).

**Figure S6:** Impact of c-di-GMP over the EPS production of subcollection *Sphingomonas* Strains. Representation the Total Carbohydrate Content (TCC) ratios of the *pleD*\* *vs*. the empty vector culture supernatants of each strain (Table S1). TCC Ratios PleD*/WT (in blue); TCC ratios PleD*/WT normalised with the OD_600nm_ (TCC/OD; in black); TCC ratios PleD*/WT normalised with the Total Protein Content (TCC/TPC; in orange) are shown.

**Figure S7:** Impact of c-di-GMP increment on the growth on solid and liquid media of SphC10 strain. A) Phenotype of SphC10 during the High Throughput Screening (HTS) after grown 42 hours in EPS solid media. Square indicate the pair: SphC10 with the empty vector pJB3Tc19 on the left vs. SphC10 pJBpleD* on the right. B) Colony morphology of SphC10 strain retransformed with pJB3Tc19 and pJBpleD*, after grown 42 hours in EPS solid selective media. C) Phenotype of SphC10 pJB3Tc19 and pJBpleD after grown 42 hours in flasks with EPS liquid selective media.

**Figure S8:** ATM-FTIR (A), and ^1^H-NMR (353 K, 500 MHz) (B) spectra, together with Total ion Monitoring (TIC) GC-MS chromatogram from monosaccharide analysis (C) of samples EPS-1 and EPS-2 isolated from SphC10.

