## Supplemental Material for "Bioprospecting c-di-GMP activated exopolysaccharides in bacteria"

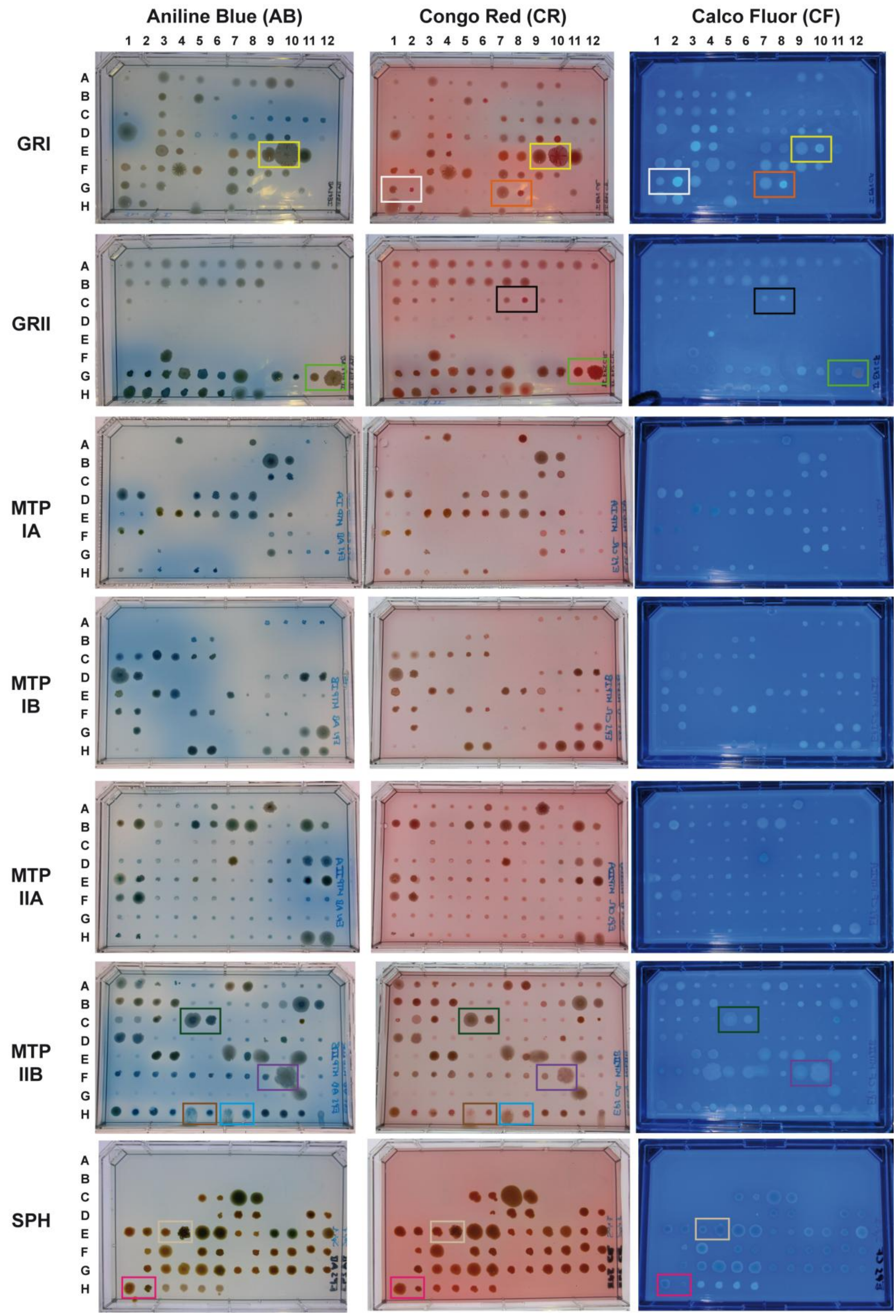


**Figure S1: Morphtype of the different strains.** Phenotype of the different strains with pJBpleD* (even columns) and with the empty vector pJB3Tc19 (odd columns) after grown 72 hours in EPS solid media supplemented with 3 different dyes: Congo Red (CR), Aniline Blue (AB) and Calcofluor (CF). Plates with CF were imaged under UV light. Square plates containing the solid media with the different dyes were inoculated with 96-well replicator from transconjugants backup plates. Squares of the same colour indicate pairs (empty vector *vs. pleD**) of relevant strains cited in the text.

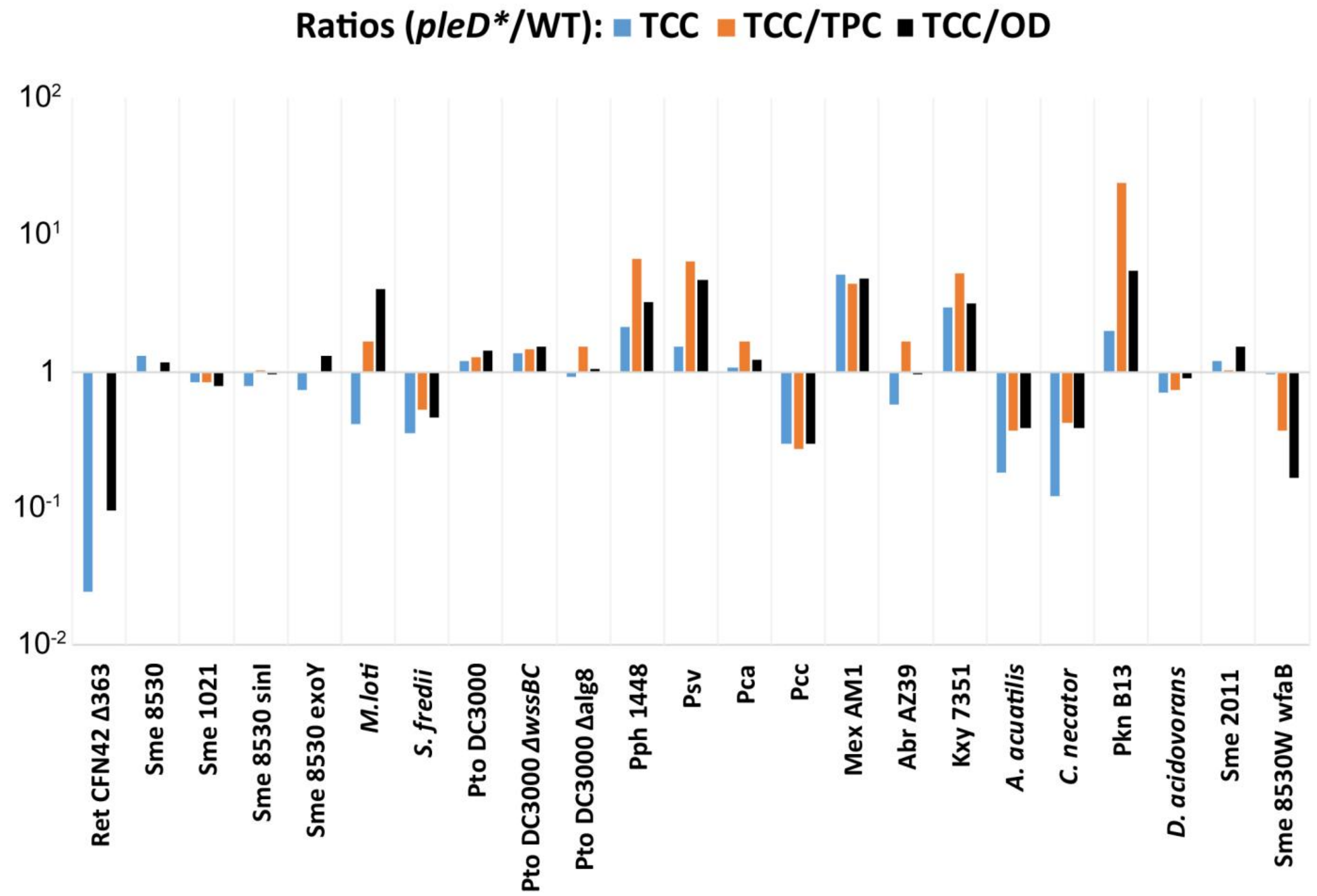


**Figure S2: Impact of c-di-GMP over the EPS production of subcollection Reference Strains.** Representation the Total Carbohydrate Content (TCC) ratios of the *pleD** *vs.* the empty vector culture supernatants of each strain (Table S4). TCC Ratios PleD*/WT (in blue); TCC ratios PleD*/WT normalised with the $OD_{600nm}$ (TCC/OD; in black); TCC ratios PleD*/WT normalised with the Total Protein Content (TCC/TPC; in orange) are shown.

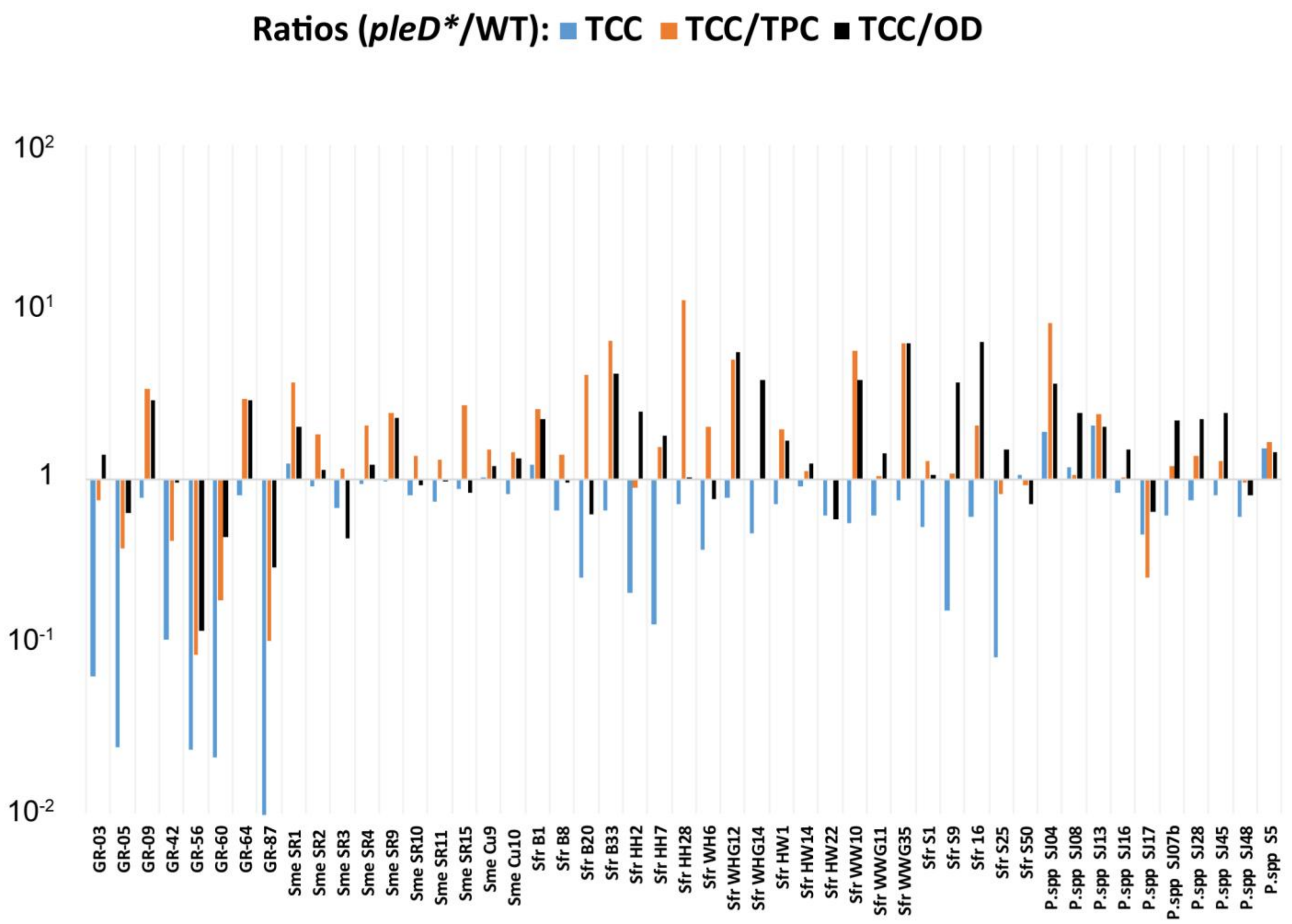


**Figure S3: Impact of c-di-GMP over the EPS production of subcollection Rhizobacteria Isolates.** Representation the Total Carbohydrate Content (TCC) ratios of the *pleD** *vs.* the empty vector culture supernatants of each strain (Table S4). TCC Ratios PleD*/WT (in blue); TCC ratios PleD*/WT normalised with the $OD_{600nm}$ (TCC/OD; in black); TCC ratios PleD*/WT normalised with the Total Protein Content (TCC/TPC; in orange) are shown.

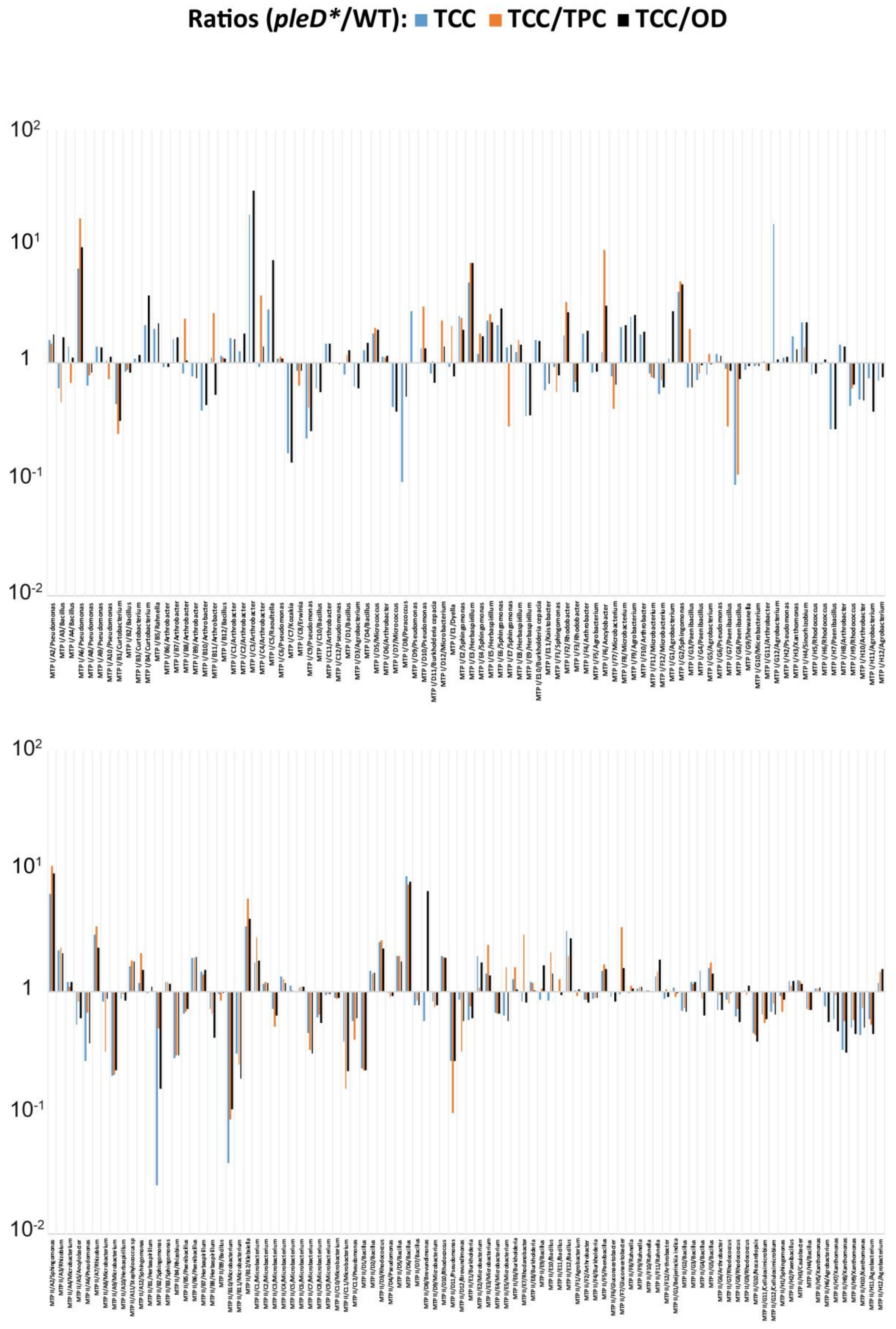


**Figure S4: Impact of c-di-GMP over the EPS production of subcollection Environmental Isolates.** Representation the Total Carbohydrate Content (TCC) ratios of the *pleD** *vs* the empty vector culture supernatants of each strain (Table S4). TCC Ratios PleD*/WT (in blue); TCC ratios PleD*/WT normalised with the $OD_{600nm}$ (TCC/OD; in black); TCC ratios PleD*/WT normalised with the Total Protein Content (TCC/TPC; in orange) are shown.

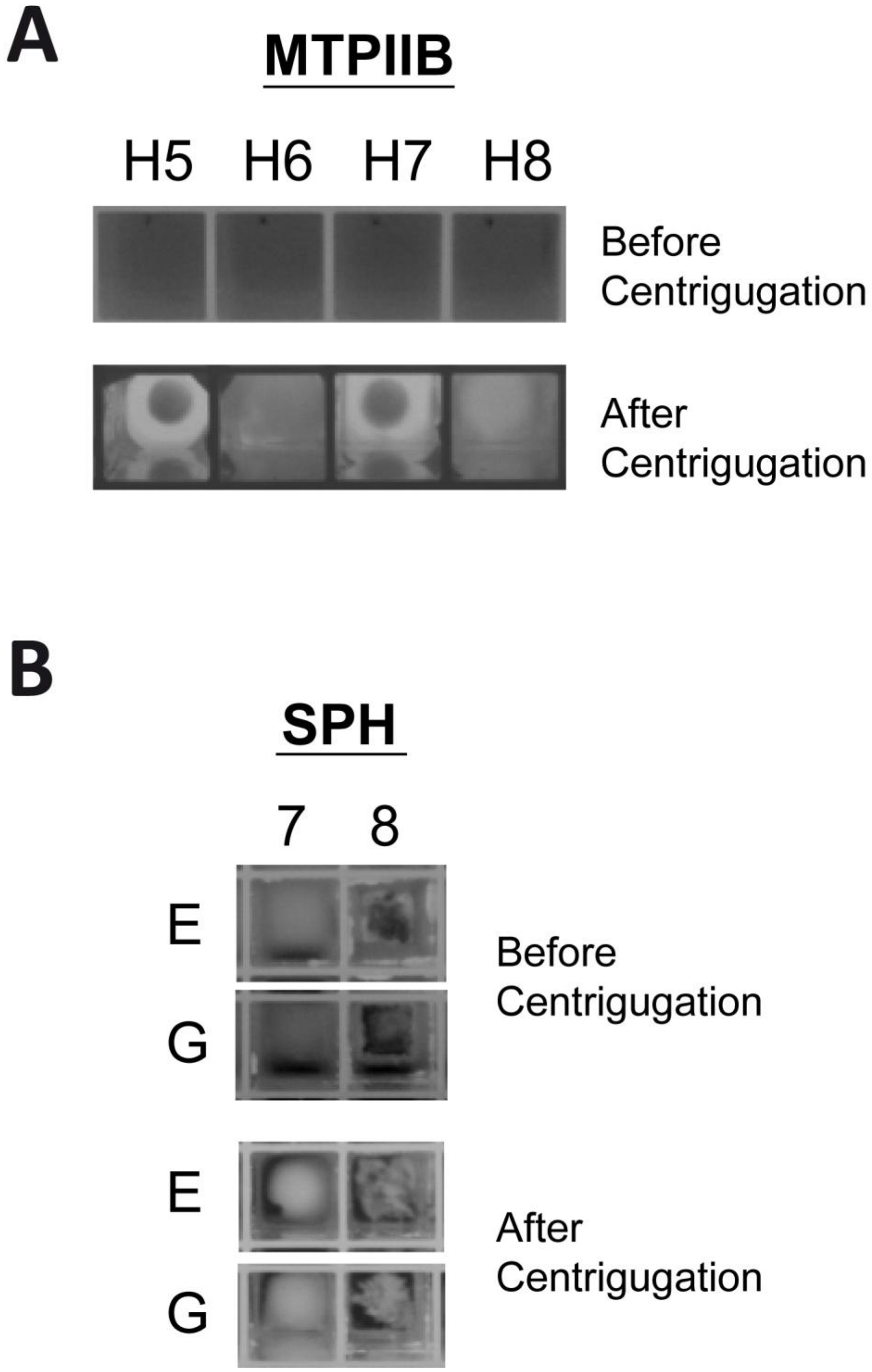


**Figure S5: Phenotype of different strains in liquid cultures.** Phenotype of different strains after grown in Deep Well Plates (DWP) with EPS liquid media for 48 hours before and after centrifugation of the DWP. A) Two different *Xanthomonas* strains from MTP IIB plate with pJBpleD* (H6 and H8) and with the empty vector pJB3Tc19 (H5 and H7). B) Two different *Sphingomonas* strains from SPH plate with pJBpleD* (E8 and G8) and with the empty vector pJB3Tc19 (E7 and G7).

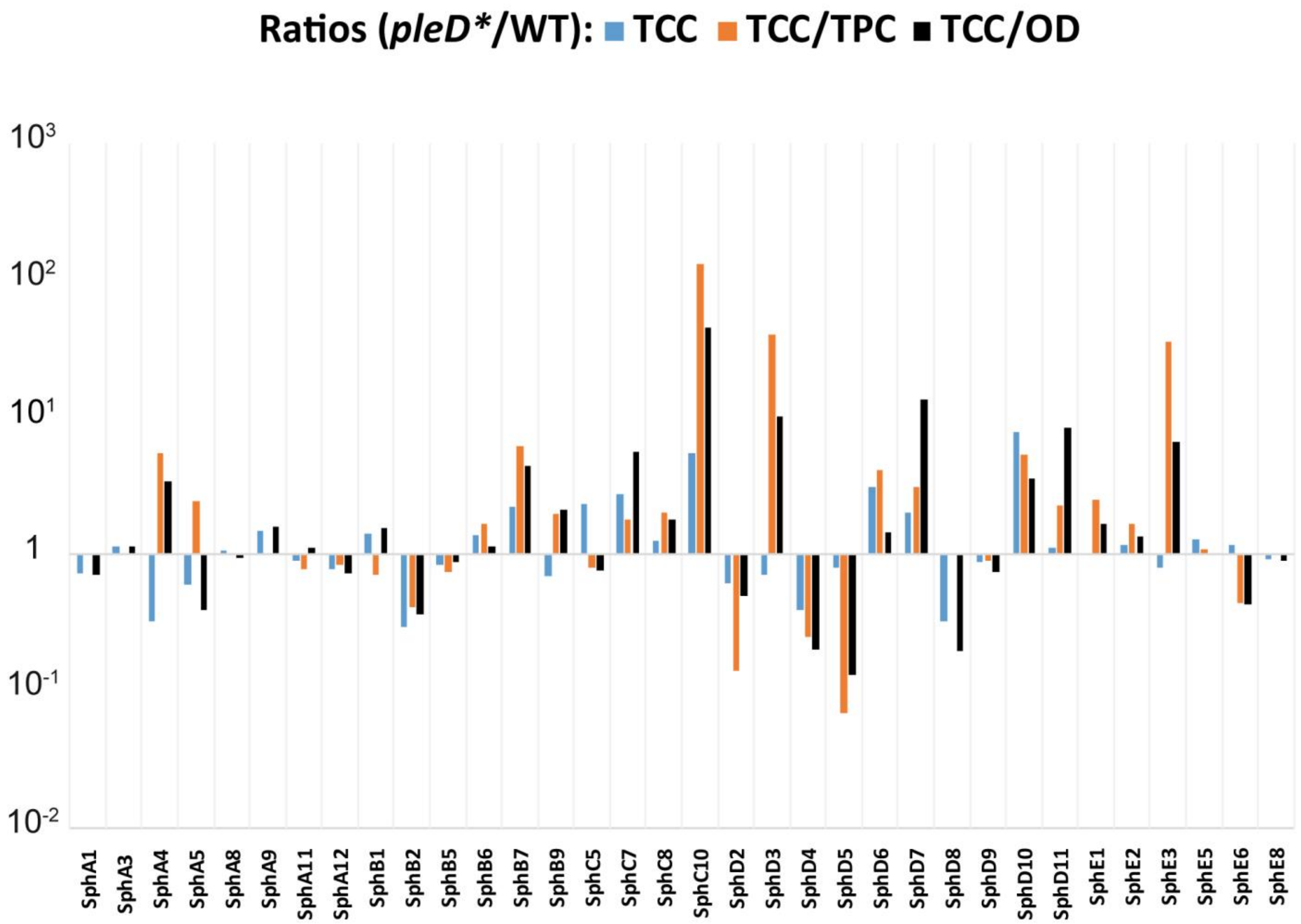


**Figure S6: Impact of c-di-GMP over the EPS production of subcollection *Sphingomonas* Strains.** Representation the Total Carbohydrate Content (TCC) ratios of the *pleD** *vs.* the empty vector culture supernatants of each strain (Table S1). TCC Ratios PleD*/WT (in blue); TCC ratios PleD*/WT normalised with the $OD_{600nm}$ (TCC/OD; in black); TCC ratios PleD*/WT normalised with the Total Protein Content (TCC/TPC; in orange) are shown.

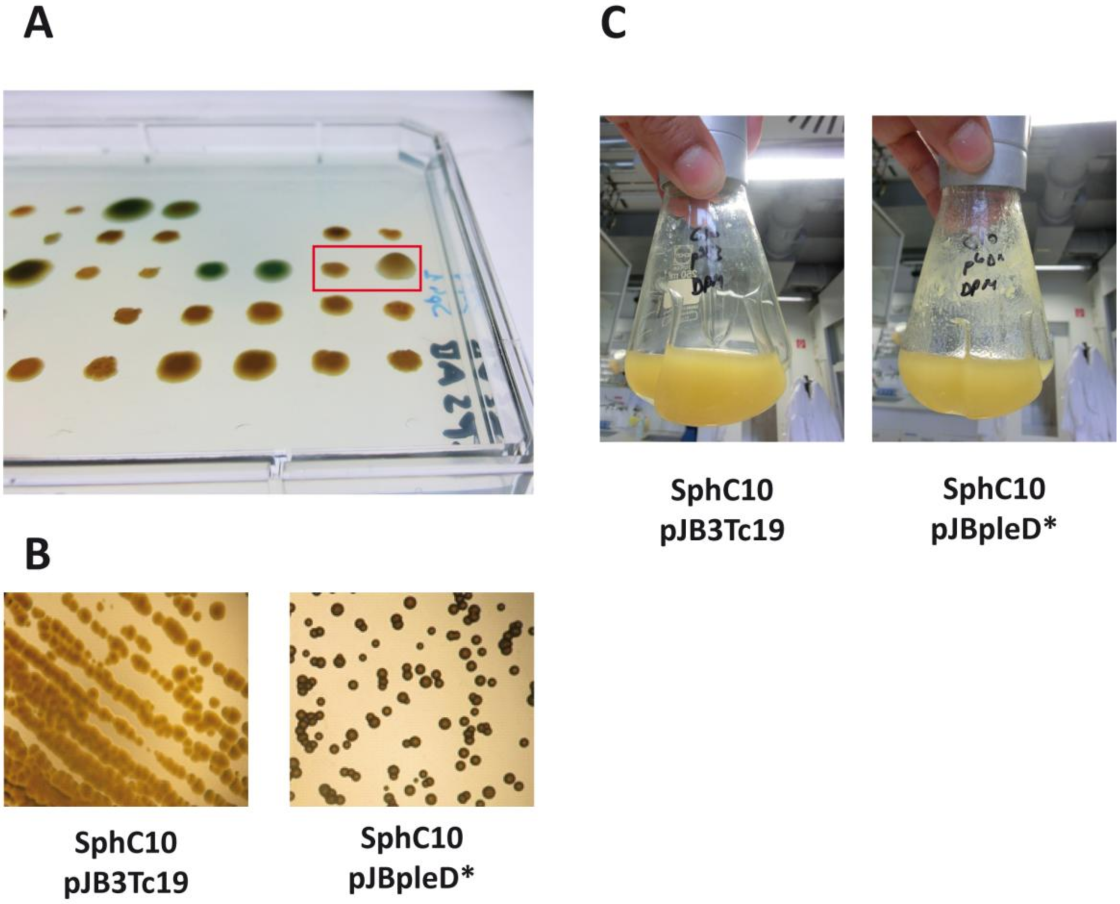


**Figure S7: Impact of c-di-GMP increment on the growth on solid and liquid media of SphC10 strain.** A) Phenotype of SphC10 during the High Throughput Screening (HTS) after grown 42 hours in EPS solid media. Square indicate the pair: SphC10 with the empty vector pJB3Tc19 on the left vs. SphC10 pJBpleD* on the right. B) Colony morphology of SphC10 strain retransformed with pJB3Tc19 and pJBpleD*, after grown 42 hours in EPS solid selective media. C) Phenotype of SphC10 pJB3Tc19 and pJBpleD after grown 42 hours in flasks with EPS liquid selective media.

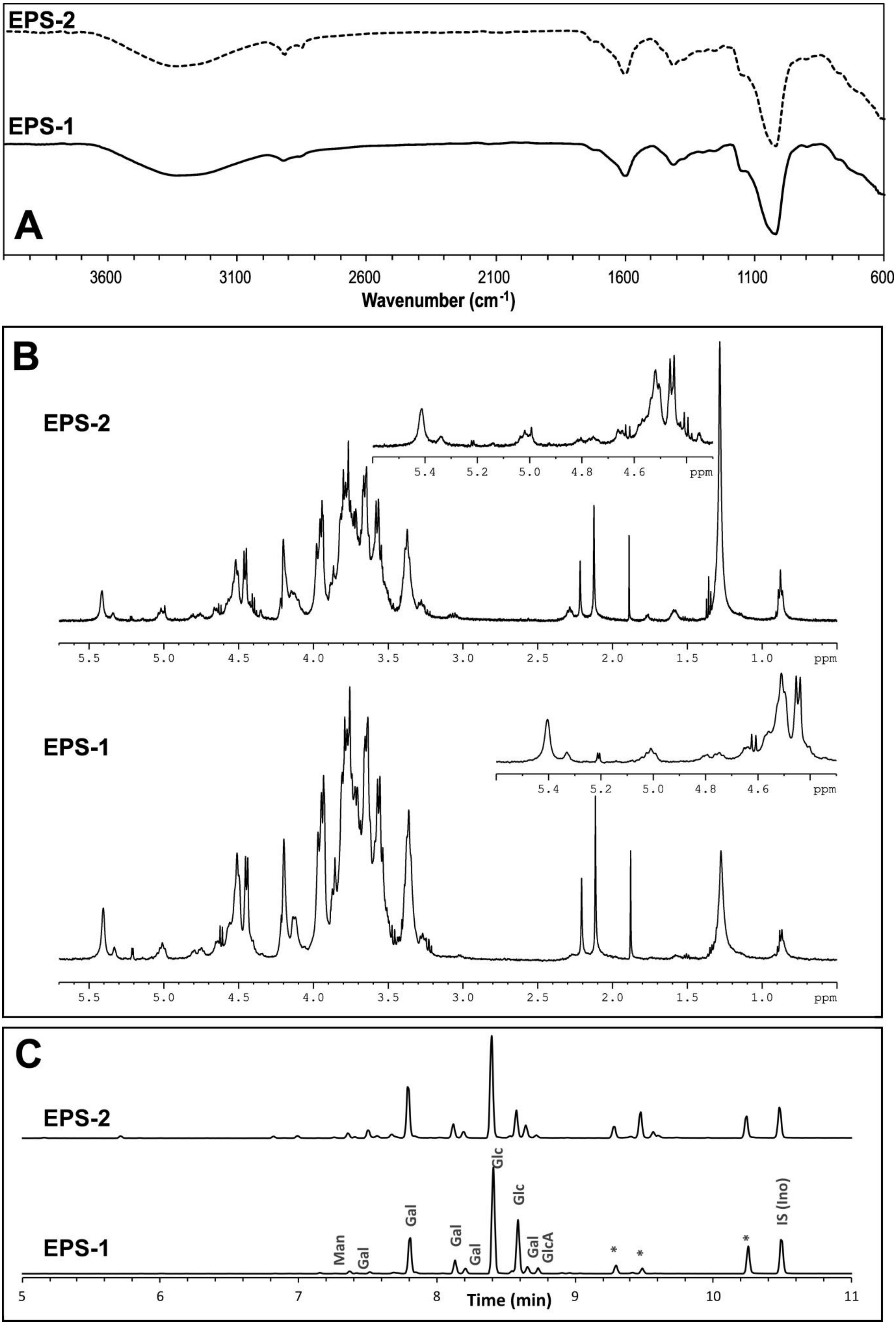


**Figure S8:** ATM-FTIR (A), and $^{1}$H-NMR (353 K, 500 MHz) (B) spectra, together with Total ion Monitoring (TIC) GC-MS chromatogram from monosaccharide analysis (C) of samples EPS-1 and EPS-2 isolated from SphC10.

### Table S1. Bacterial strains analysed

| Strain | Relevant Characteristics | Original Plate |
|---|---|---|
| Ret CFN42 | Rhizobium etli CFN42; Wild-type | GRI |
| RetCFN42 ΔcelAB | Rhizobium etli CFN42; Cel- | GRI |
| Ret CFN42 Δ363 | Rhizobium etli CFN42; MLG- | GRI |
| Ret CFN42 ΔcelABΔ363 | Rhizobium etli CFN42; Cel- MLG- | GRI |
| Sme 8530 | Ensifer meliloti 8530; Wild-type | GRI |
| Sme 1021 | Ensifer meliloti 1021; Wild-type | GRI |
| Sme 8530 *sinI* | Ensifer meliloti 8530; *sinI* mutant | GRI |
| Sme 8530 *exoY* | Ensifer meliloti 8530; EPSI- | GRI |
| Sme GR4 | Ensifer meliloti GR4; Wild-type | GRI |
| M.loti MAFF303099 | Mesorhizobium loti MAFF303099; Wild-type | GRI |
| Rle 8341 | Rhizobium leguminosarum 8341; Wild-type | GRI |
| Rle UPM791 | Rhizobium leguminosarum UPM791; Wild-type | GRI |
| S. fredii B1 | Ensifer fredii B1; Wild-type | GRI |
| S. fredii NGR234 | Ensifer fredii NGR234; Wild-type | GRI |
| S. fredii USDA257 | Ensifer fredii USDA258; Wild-type | GRI |
| Pto DC3000 | Pseudomonas syringae DC3000; Wild-type | GRI |
| Pto DC3000 Δ*wssBC* | Pseudomonas syringae DC3000; Cel- | GRI |
| Pto DC3000 Δ*alg8* | Pseudomonas syringae DC3000; Alg- | GRI |
| Pph 1448 | Pseudomomas phaseolicola 1448; Wild-type | GRI |
| Psv | Pseudomomas savastanoi; pv. savastanoi Wild-type | GRI |
| Pca SCRI1043 | Pectobacterium carotovorum Subsp.atrosepticum SCRI1043; Wild-type | GRI |
| Pcc SCRI193 | Pectobacterium carotovorum Subsp. carotovorum SCRI193 Wild-type | GRI |
| Atu LBA1010 | Agrobacterium tumefaciens LBA1010 Wild-type | GRI |
| Mex PA1 | Methylobacterium extorquens PA1; Wild-type | GRI |
| Mex AM1 | Methylobacterium extorquens AM1; Wild-type | GRI |
| Abr AZ39 | Azospirillum brasilense AZ39; Wild-type | GRI |
| Kxy 7351 | Komagataeibacter xylinus 7351; Wild-type; PRCC collection | GRI |
| A. acuatilis | Alcaligenes aquatilis, PRCC collection | GRI |
| Cupriavidus necator | Cupriavidus necator; PRCC collection | GRI |
| Pkn B13 | Pseudomonas Knackmussii B13; PRCC collection | GRI |
| Delftia acidovorans | Delftia acidovorans; PRCC collection | GRI |
| Ain LnG9092 | Azoarcus indigenes LnG 9092; PRCC collection | GRI |
| Ach denitrificans | Achromobacter denitrificans; PRCC collection | GRI |
| Sme 2011 | Ensifer meliloti 2011; Wild-type | GRI |
| Sme 8530W | Ensifer meliloti 8530; EPSII- | GRI |
| GR-03 | Bacterial isolate from bean nodules in Granada soils (Spain) | GRI |
| GR-05 | Bacterial isolate from bean nodules in Granada soils (Spain) | GRI |
| GR-09 | Bacterial isolate from bean nodules in Granada soils (Spain) | GRI |
| GR-42 | Bacterial isolate from bean nodules in Granada soils (Spain) | GRI |
| GR-45 | Bacterial isolate from bean nodules in Granada soils (Spain) | GRI |
| GR-56 | Bacterial isolate from bean nodules in Granada soils (Spain) | GRI |
| GR-60 | Bacterial isolate from bean nodules in Granada soils (Spain) | GRI |
| GR-64 | Bacterial isolate from bean nodules in Granada soils (Spain) | GRI |
| GR-84 | Bacterial isolate from bean nodules in Granada soils (Spain) | GRI |
| GR-87 | Bacterial isolate from bean nodules in Granada soils (Spain) | GRI |
| Sme SR1 | Bacterial isolate from alfalfa nodules in Angentina soils | GRII |
| Sme SR2 | Bacterial isolate from alfalfa nodules in Angentina soils | GRII |
| Sme SR3 | Bacterial isolate from alfalfa nodules in Angentina soils | GRII |
| Sme SR4 | Bacterial isolate from alfalfa nodules in Angentina soils | GRII |
| Sme SR9 | Bacterial isolate from alfalfa nodules in Angentina soils | GRII |
| Sme SR10 | Bacterial isolate from alfalfa nodules in Angentina soils | GRII |
| Sme SR11 | Bacterial isolate from alfalfa nodules in Angentina soils | GRII |
| Sme SR15 | Bacterial isolate from alfalfa nodules in Angentina soils | GRII |

| | | |
|---|---|---|
| Sme Cu9 | Bacterial isolate from alfalfa nodules in Angentina soils | GRII |
| Sme Cu10 | Bacterial isolate from alfalfa nodules in Angentina soils | GRII |
| Sfr B1 | Bacterial isolate from soybean nodules in China soils | GRII |
| Sfr B8 | Bacterial isolate from soybean nodules in China soils | GRII |
| Sfr B20 | Bacterial isolate from soybean nodules in China soils | GRII |
| Sfr B33 | Bacterial isolate from soybean nodules in China soils | GRII |
| Sfr HH2 | Bacterial isolate from soybean nodules in China soils | GRII |
| Sfr HH7 | Bacterial isolate from soybean nodules in China soils | GRII |
| Sfr HH28 | Bacterial isolate from soybean nodules in China soils | GRII |
| Sfr WH6 | Bacterial isolate from soybean nodules in China soils | GRII |
| Sfr WHG12 | Bacterial isolate from soybean nodules in China soils | GRII |
| Sfr WHG14 | Bacterial isolate from soybean nodules in China soils | GRII |
| Sfr HW1 | Bacterial isolate from soybean nodules in China soils | GRII |
| Sfr HW14 | Bacterial isolate from soybean nodules in China soils | GRII |
| Sfr HW22 | Bacterial isolate from soybean nodules in China soils | GRII |
| Sfr WW10 | Bacterial isolate from soybean nodules in China soils | GRII |
| Sfr WWG11 | Bacterial isolate from soybean nodules in China soils | GRII |
| Sfr WWG35 | Bacterial isolate from soybean nodules in China soils | GRII |
| Sfr S1 | Bacterial isolate from soybean nodules in China soils | GRII |
| Sfr S9 | Bacterial isolate from soybean nodules in China soils | GRII |
| Sfr 16 | Bacterial isolate from soybean nodules in China soils | GRII |
| Sfr S25 | Bacterial isolate from soybean nodules in China soils | GRII |
| Sfr S50 | Bacterial isolate from soybean nodules in China soils | GRII |
| P.spp SJ04 | Bacterial isolate from peppermint rizosphere in Argentina soils | GRII |
| P.spp SJ08 | Bacterial isolate from peppermint rizosphere in Argentina soils | GRII |
| P.spp SJ13 | Bacterial isolate from peppermint rizosphere in Argentina soils | GRII |
| P.spp SJ16 | Bacterial isolate from peppermint rizosphere in Argentina soils | GRII |
| P.spp SJ17 | Bacterial isolate from peppermint rizosphere in Argentina soils | GRII |
| P.spp SJ07b | Bacterial isolate from peppermint rizosphere in Argentina soils | GRII |
| P.spp SJ28 | Bacterial isolate from peppermint rizosphere in Argentina soils | GRII |
| P.spp SJ45 | Bacterial isolate from peppermint rizosphere in Argentina soils | GRII |
| P.spp SJ48 | Bacterial isolate from peppermint rizosphere in Argentina soils | GRII |
| P.spp S5 | Bacterial isolate from peppermint rizosphere in Argentina soils | GRII |
| MTP I/A2/Pseudomonas | plum | MTP I |
| MTP I/A3/Bacillus | windfall, apple | MTP I |
| MTP I/A4/Bacillus | leaf of a quince tree | MTP I |
| MTP I/A6/Pseudomonas | mix of exotic fruits | MTP I |
| MTP I/A8/Pseudomonas | mix of exotic fruits | MTP I |
| MTP I/A9/Pseudomonas | mix of exotic fruits | MTP I |
| MTP I/A10/Pseudomonas | salsify | MTP I |
| MTP I/B1/Curtobacterium | chinese yam | MTP I |
| MTP I/B2/Bacillus | chinese yam | MTP I |
| MTP I/B3/Curtobacterium | chinese yam | MTP I |
| MTP I/B4/Curtobacterium | Sapodilla | MTP I |
| MTP I/B5/Rahnella | Sapodilla | MTP I |
| MTP I/B6/Arthrobacter | Sapodilla | MTP I |
| MTP I/B7/Arthrobacter | contamination | MTP I |
| MTP I/B8/Arthrobacter | contamination | MTP I |
| MTP I/B9/Arthrobacter | contamination | MTP I |
| MTP I/B10/Arthrobacter | contamination | MTP I |
| MTP I/B11/Arthrobacter | contamination | MTP I |
| MTP I/B12/Bacillus | contamination | MTP I |
| MTP I/C1/Arthrobacter | contamination | MTP I |
| MTP I/C2/Arthrobacter | contamination | MTP I |
| MTP I/C3/Arthrobacter | contamination | MTP I |
| MTP I/C4/Arthrobacter | contamination | MTP I |
| MTP I/C5/Raoultella | tulip stipe | MTP I |
| MTP I/C6/Pseudomonas | parsnip | MTP I |
| MTP I/C7/Kozakia | passion fruit | MTP I |

| | | |
|---|---|---|
| MTP I/C8/Erwinia | salacca | MTP I |
| MTP I/C9/Pseudomonas | aquarium alge | MTP I |
| MTP I/C10/Bacillus | activated sludge | MTP I |
| MTP I/C11/Arthrobacter | parsnip | MTP I |
| MTP I/C12/Pseudomonas | plantain | MTP I |
| MTP I/D1/Bacillus | contamination | MTP I |
| MTP I/D3/Agrobacterium | septic tulip stipe | MTP I |
| MTP I/D4/Bacillus | soil | MTP I |
| MTP I/D5/Micrococcus | airborne germ | MTP I |
| MTP I/D6/Arthrobacter | garden mold | MTP I |
| MTP I/D7/Micrococcus | contamination | MTP I |
| MTP I/D8/Paracoccus | strain collection 2 | MTP I |
| MTP I/D9/Pseudomonas | soil | MTP I |
| MTP I/D10/Pseudomonas | soil | MTP I |
| MTP I/D11/Burkholderia cepacia | soil | MTP I |
| MTP I/D12/Microbacterium | soil Holophaga | MTP I |
| MTP I/E1/Dyella | soil Holophaga | MTP I |
| MTP I/E2/Sphingomonas | paper mill | MTP I |
| MTP I/E3/Herbaspirillum | paper mill | MTP I |
| MTP I/E4/Sphingomonas | paper mill | MTP I |
| MTP I/E5/Herbaspirillum | paper mill | MTP I |
| MTP I/E6/Sphingomonas | paper mill | MTP I |
| MTP I/E7/Sphingomonas | paper mill | MTP I |
| MTP I/E8/Herbaspirillum | paper mill | MTP I |
| MTP I/E9/Herbaspirillum | paper mill | MTP I |
| MTP I/E10/Burkholderia cepacia | paper mill | MTP I |
| MTP I/E11/Kaistobacter | paper mill | MTP I |
| MTP I/F1/Sphingomonas | paper mill | MTP I |
| MTP I/F2/Rhodobacter | paper mill | MTP I |
| MTP I/F3/Rhodobacter | paper mill | MTP I |
| MTP I/F4/Arthrobacter | Soil BRAIN | MTP I |
| MTP I/F5/Agrobacterium | Soil BRAIN | MTP I |
| MTP I/F6/Ancylobacter | Soil BRAIN | MTP I |
| MTP I/F7/Microbacterium | Soil BRAIN | MTP I |
| MTP I/F8/Microbacterium | Soil BRAIN | MTP I |
| MTP I/F9/Agrobacterium | soil enrichment | MTP I |
| MTP I/F10/Arthrobacter | soil enrichment | MTP I |
| MTP I/F11/Microbacterium | soil enrichment | MTP I |
| MTP I/F12/Microbacterium | soil enrichment | MTP I |
| MTP I/G1/Agrobacterium | strain collection | MTP I |
| MTP I/G2/Sphingomonas | strain collection | MTP I |
| MTP I/G3/Paenibacillus | strain collection 2 | MTP I |
| MTP I/G4/Paenibacillus | strain collection 2 | MTP I |
| MTP I/G5/Agrobacterium | strain collection 2 | MTP I |
| MTP I/G6/Pseudomonas | strain collection 2 | MTP I |
| MTP I/G7/Paenibacillus | strain collection 2 | MTP I |
| MTP I/G8/Paenibacillus | strain collection 2 | MTP I |
| MTP I/G9/Shewanella | strain collection | MTP I |
| MTP I/G10/Microbacterium | curacao | MTP I |
| MTP I/G11/Arthrobacter | soil | MTP I |
| MTP I/G12/Agrobacterium | strain collection | MTP I |
| MTP I/H2/Pseudomonas | strain collection | MTP I |
| MTP I/H3/Xanthomonas | strain collection | MTP I |
| MTP I/H4/Sinorhizobium | strain collection | MTP I |
| MTP I/H5/Rhodococcus | strain collection | MTP I |
| MTP I/H6/Rhodococcus | strain collection | MTP I |
| MTP I/H7/Paenibacillus | soil | MTP I |
| MTP I/H8/Arthrobacter | alkane containing sample | MTP I |
| MTP I/H9/Rhodococcus | strain collection | MTP I |

| | | |
|---|---|---|
| MTP I/H10/Arthrobacter | strain collection | MTP I |
| MTP I/H11/Agrobacterium | strain collection | MTP I |
| MTP I/H12/Agrobacterium | strain collection | MTP I |
| MTP II/A2/Sphingomonas | soil enrichment | MTP II |
| MTP II/A3/Rhizobium | soil enrichment | MTP II |
| MTP II/A4/Microbacterium | soil enrichment | MTP II |
| MTP II/A5/Ancylobacter | soil enrichment | MTP II |
| MTP II/A6/Pseudomonas | water well | MTP II |
| MTP II/A7/Rhizobium | soil BRAIN | MTP II |
| MTP II/A8/Microbacterium | soil BRAIN | MTP II |
| MTP II/A9/Microbacterium | soil BRAIN | MTP II |
| MTP II/A10/Herbaspirillum | paper mill | MTP II |
| MTP II/A11/Staphylococcus sp | paper mill | MTP II |
| MTP II/A12/Sphingomonas | paper mill | MTP II |
| MTP II/B1/Herbaspirillum | paper mill | MTP II |
| MTP II/B2/Sphingomonas | paper mill | MTP II |
| MTP II/B3/Sphingomonas | paper mill | MTP II |
| MTP II/B4/Rhizobium | contamination | MTP II |
| MTP II/B5/Paenibacillus | soil Aralie | MTP II |
| MTP II/B6/Paenibacillus | soil Aralie | MTP II |
| MTP II/B7/Herbaspirillum | paper mill | MTP II |
| MTP II/B8/Herbaspirillum | paper mill | MTP II |
| MTP II/B9/Bacillus | soil BRAIN | MTP II |
| MTP II/B10/Microbacterium | sugar rich sample | MTP II |
| MTP II/B11/Microbacterium | sugar rich sample | MTP II |
| MTP II/B12/Klebsiella | sugar rich sample | MTP II |
| MTP II/C1/Microbacterium | sugar rich sample | MTP II |
| MTP II/C2/Microbacterium | sugar rich sample | MTP II |
| MTP II/C3/Microbacterium | sugar rich sample | MTP II |
| MTP II/C4/Microbacterium | sugar rich sample | MTP II |
| MTP II/C5/Microbacterium | sugar rich sample | MTP II |
| MTP II/C6/Microbacterium | sugar rich sample | MTP II |
| MTP II/C7/Microbacterium | sugar rich sample | MTP II |
| MTP II/C8/Microbacterium | sugar rich sample | MTP II |
| MTP II/C9/Microbacterium | sugar rich sample | MTP II |
| MTP II/C10/Microbacterium | sugar rich sample | MTP II |
| MTP II/C11/Microbacterium | sugar rich sample | MTP II |
| MTP II/C12/Pseudomonas | sugar rich sample | MTP II |
| MTP II/D1/Bacillus | sugar rich sample | MTP II |
| MTP II/D2/Bacillus | sugar rich sample | MTP II |
| MTP II/D3/Rhodococcus | sugar rich sample | MTP II |
| MTP II/D4/Pseudomonas | sugar rich sample | MTP II |
| MTP II/D5/Bacillus | sugar rich sample | MTP II |
| MTP II/D6/Bacillus | sugar rich sample | MTP II |
| MTP II/D7/Bacillus | sugar rich sample | MTP II |
| MTP II/D8/Brevundimonas | sugar rich sample | MTP II |
| MTP II/D9/Microbacterium | sugar rich sample | MTP II |
| MTP II/D10/Rhodococcus | sugar rich sample | MTP II |
| MTP II/D11/Pseudomonas | cactus | MTP II |
| MTP II/D12/Brevundimonas | cactus | MTP II |
| MTP II/E1/Burkholderia | cactus | MTP II |
| MTP II/E2/Microbacterium | cactus | MTP II |
| MTP II/E3/Microbacterium | cactus | MTP II |
| MTP II/E4/Microbacterium | cactus | MTP II |
| MTP II/E5/Microbacterium | cactus | MTP II |
| MTP II/E6/Burkholderia | cactus | MTP II |
| MTP II/E7/Rhodanobacter | cactus | MTP II |
| MTP II/E8/Burkholderia | cactus | MTP II |
| MTP II/E9/Bacillus | cockchafer dung | MTP II |

| | | |
|---|---|---|
| MTP II/E10/Bacillus | cockchafer dung | MTP II |
| MTP II/E11/Bacillus | contamination | MTP II |
| MTP II/E12/Bacillus | contamination | MTP II |
| MTP II/F1/Agrobacterium | soil | MTP II |
| MTP II/F2/Arthrobacter | soil | MTP II |
| MTP II/F4/Burkholderia | cactus | MTP II |
| MTP II/F5/Paenibacillus | soil | MTP II |
| MTP II/F6/Gluconacetobacter | soil | MTP II |
| MTP II/F7/Gluconacetobacter | soil | MTP II |
| MTP II/F8/Rahnella | carrot | MTP II |
| MTP II/F9/Rahnella | carrot | MTP II |
| MTP II/F10/Rahnella | carrot | MTP II |
| MTP II/F11/Rahnella | carrot | MTP II |
| MTP II/F12/Arthrobacter | soil | MTP II |
| MTP II/G1/Beijerinckia indica | DSMZ 1715 | MTP II |
| MTP II/G2/Bacillus | soil | MTP II |
| MTP II/G3/Bacillus | soil | MTP II |
| MTP II/G4/Bacillus | soil | MTP II |
| MTP II/G5/Bacillus | soil | MTP II |
| MTP II/G6/Arthrobacter | soil | MTP II |
| MTP II/G7/Rhodococcus | strain collection 2 | MTP II |
| MTP II/G8/Rhodococcus | strain collection 2 | MTP II |
| MTP II/G9/Rhodococcus | strain collection | MTP II |
| MTP II/G10/Nocardiopsis | strain collection 2 | MTP II |
| MTP II/G11/Cellulosimicrobium | soil | MTP II |
| MTP II/G12/Cellulosimicrobium | soil | MTP II |
| MTP II/H1/Sphingomonas | compost | MTP II |
| MTP II/H2/Paenibacillus | strain collection 2 | MTP II |
| MTP II/H3/Caulobacter | strain collection | MTP II |
| MTP II/H4/Bacillus | strain collection | MTP II |
| MTP II/H5/Xanthomonas | strain collection | MTP II |
| MTP II/H6/Agrobacterium | strain collection | MTP II |
| MTP II/H7/Xanthomonas | strain collection | MTP II |
| MTP II/H8/Xanthomonas | strain collection | MTP II |
| MTP II/H9/Xanthomonas | strain collection | MTP II |
| MTP II/H10/Xanthomonas | strain collection | MTP II |
| MTP II/H11/Agrobacterium | strain collection | MTP II |
| MTP II/H12/Agrobacterium | strain collection | MTP II |
| SphA1 Sphingomonas alaskensis | on020912a M. Wenning | SPH |
| SphA2 Sphingomonas paucimobilis | st021009 M. Wenning | SPH |
| SphA3 Sphingomonas paucimobilis | no011113 M. Wenning | SPH |
| SphA4 Sphingomonas paucimobilis | mü030313 M. Wenning | SPH |
| SphA5 Sphingomonas yanoikuyae | Kistner(W1) M. Wenning | SPH |
| SphA6 Sphingomonas sp. (wittichii) | mw040317 M. Wenning | SPH |
| SphA7 Sphingomonas subterranea | mw040317 M. Wenning | SPH |
| SphA8 Sphingomonas sp. (wittichii) | mw040526 M. Wenning | SPH |
| SphA9 Sphingomonas sp. | on040804 M. Wenning | SPH |
| SphA10 Sphingomonas sp. (yanoikuyae) | on040902 M. Wenning | SPH |
| SphA11 Sphingomonas sp. (aurantiaca) | sa041011 M. Wenning | SPH |
| SphA12 Sphingomonas yabuuchiae | lvt041012 M. Wenning | SPH |
| SphB1 Sphingomonas echinoides | lvt041012 1-2f M. Wenning | SPH |
| SphB2 Sphingomonas melonis | lvt041012 5-2b M. Wenning | SPH |
| SphB3 Sphingomonas sp. (melonis, asaccharolytica) | Lang-H.(BMIE4) M. Wenning | SPH |
| SphB4 Sphingomonas sp. (aquatilis,melonis) | ga050324 M. Wenning | SPH |
| SphB5 Sphingomonas yanoikuyae | sm050509b 38057-4 M. Wenning | SPH |
| SphB6 Sphingomonas sp. | sa050614 135.2 M. Wenning | SPH |
| SphB7 Sphingomonas sp. | mw050614_29 M. Wenning | SPH |
| SphB8 Sphingomonas melonis | gs050713_2-1 M. Wenning | SPH |

| | | |
|---|---|---|
| SphB9 Sphingomonas pseudosanguinis | sm050809 39536-1 M. Wenning | SPH |
| SphB10 Sphingomonas mucosissima / anadarae | gs060117 3-5 M. Wenning | SPH |
| SphB11 Sphingomonas melonis | gs060202 2-1 M. Wenning | SPH |
| SphB12 Sphingomonas melonis | mw060206 20032a M. Wenning | SPH |
| SphC1 Sphingomonas azotifigens | on060123a 6-1 M. Wenning | SPH |
| SphC2 Sphingomonas melonis | gs060221 1-53MK M. Wenning | SPH |
| SphC3 Sphingomonas yabuuchiae (intermedia) | sm060524a 43477-1 M. Wenning | SPH |
| SphC4 Sphingomonas yabuuchiae | sm060627b 43802-1 M. Wenning | SPH |
| SphC5 Sphingomonas mathurensis | fromG3842 M. Wenning | SPH |
| SphC6 nd | M. Wenning | SPH |
| SphC7 Sphingomonas sp. (melonis) | gs070302 2-1 M. Wenning | SPH |
| SphC8 Sphingomonas yanoikuyae | hok070719 4-1 M. Wenning | SPH |
| SphC9 Sphingomonas melonis | gs071204b 1-1 M. Wenning | SPH |
| SphC10 Sphingomonas yabuuchiae | imd_M 367.3 M. Wenning | SPH |
| SphC11 Sphingomonas koreensis | oe081205 3-1 M. Wenning | SPH |
| SphC12 Sphingomonas paucimobilis | imd 78647-1 M. Wenning | SPH |
| SphD1 nd | M.Wenning | SPH |
| SphD2 Sphingomonas sp. (melonis, aquatilis) | imd 79799_1Keim M. Wenning | SPH |
| SphD3 Sphingomonas paucimobilis | imd IV426 M. Wenning | SPH |
| SphD4 Sphingomonas aquatilis | imd 80569_4Keim_I M. Wenning | SPH |
| SphD5 Sphingomonas melonis | imd 86915I M. Wenning | SPH |
| SphD6 nd | M.Wenning | SPH |
| SphD7 nd | M.Wenning | SPH |
| SphD8 Sphingomonas melonis | imd 95728/2 II M. Wenning | SPH |
| SphD9 Sphingomonas paucimobilis | imd 92312 M. Wenning | SPH |
| SphD10 Sphingomonas paucimobilis | gs100630 Hi2_1Pr.-1 M. Wenning | SPH |
| SphD11 nd | M.Wenning | SPH |
| SphD12 Sphingomonas yabuuchiae | emw101217 1-1 M. Wenning | SPH |
| SphE1 Sphingomonas pseudosanguinis | imd 123787-2 M. Wenning | SPH |
| SphE2 Sphingomonas melonis | imd 1445.7 M. Wenning | SPH |
| SphE3 Sphingomonas paucimobilis | imd water M. Wenning | SPH |
| SphE4 nd | M.Wenning | SPH |
| SphE5 Sphingomonas leidyi | imd water M. Wenning | SPH |
| SphE6 Sphingomonas aurantiaca | imd water M. Wenning | SPH |
| SphE7 nd | M.Wenning | SPH |
| SphE8 nd | M.Wenning | SPH |
| SphE9 nd | M.Wenning | SPH |
| SphE10 nd | M.Wenning | SPH |
| SphE11 nd | M.Wenning | SPH |
| SphE12 nd | M.Wenning | SPH |
| SphF1 nd | M.Wenning | SPH |
| SphF2 Sphingomonas mucosissima | imd IVP516 M. Wenning | SPH |

**Table S2. Deep Well Plates (DWP) processed for conjugation**

| Plate | Nº of Strains | Grown (OD≥0.1) | Transformed Strains Grown in Tc (OD≥0.1) | Plates (nº of transformed strains) | |
|---|---|---|---|---|---|
| GRI | 45 | - | pJB3Tc19: 45 | GRI:90 | |
| | | | pJBpleD*: 45 | | |
| GRII | 41 | - | pJB3Tc19: 41 | GRII:82 | |
| | | | pJBpleD*: 41 | | |
| MTP-I | 88 | 55 | pJB3Tc19: 44 | MTP-IA 88 | MTP-IB 88 |
| | | | pJBpleD*: 39 | | |
| MTP-II | 94 | 50 | pJB3Tc19: 44 | MTP-IIA 92 | MTP-IIB 96 |
| | | | pJBpleD*: 44 | | |
| SPH | 62 | 37 | pJB3Tc19: 37 | SPH: 68 | |
| | | | pJBpleD*: 35 | | |

OD ≥ 0.1: Optical Density at 600 nm > 0,1 was established as a threshold to consider that one particular strain was grown.

Table S3

| Strain | Plasmid | Plate | Well | TCC | OD | TPC [prot] µg/mL | TCC/TPC (µg/mL$^{-1}$ | TCC/OD |
|---|---|---|---|---|---|---|---|---|
| Ret CFN42 | pJB3Tc19 | GRI | A1 | nd | 0,04 | nd | 0,0 | nd |
| Ret CFN42 | pJBpleD* | GRI | A2 | nd | 0,04 | nd | 0,0 | nd |
| Ret CFN42 ΔcelAB | pJB3Tc19 | GRI | A3 | nd | 0,04 | 6 | nd | nd |
| Ret CFN42 ΔcelAB | pJBpleD* | GRI | A4 | nd | 0,04 | 2 | nd | nd |
| Ret CFN42 Δ363 | pJB3Tc19 | GRI | A5 | 611 | 0,18 | 14 | 45,2 | 3415 |
| Ret CFN42 Δ363 | pJBpleD* | GRI | A6 | 15 | 0,05 | nd | nd | 332 |
| Ret CFN42 ΔcelABΔ363 | pJB3Tc19 | GRI | A7 | 574 | 0,18 | 16 | 36,3 | 3152 |
| Ret CFN42 ΔcelABΔ363 | pJBpleD* | GRI | A8 | nd | 0,05 | 2 | nd | nd |
| Sme 8530 | pJB3Tc19 | GRI | A9 | 1180 | 0,45 | 16 | 72,7 | 2652 |
| Sme 8530 | pJBpleD* | GRI | A10 | 1554 | 0,50 | 22 | 71,6 | 3104 |
| Sme 1021 | pJB3Tc19 | GRI | A11 | 766 | 0,75 | 62 | 12,3 | 1028 |
| Sme 1021 | pJBpleD* | GRI | A12 | 653 | 0,80 | 64 | 10,3 | 815 |
| Sme 8530 *sinI* | pJB3Tc19 | GRI | B1 | 600 | 0,83 | 68 | 8,9 | 727 |
| Sme 8530 *sinI* | pJBpleD* | GRI | B2 | 472 | 0,67 | 52 | 9,1 | 706 |
| Sme 8530 *exoY* | pJB3Tc19 | GRI | B3 | 997 | 0,50 | 20 | 50,1 | 2008 |
| Sme 8530 *exoY* | pJBpleD* | GRI | B4 | 729 | 0,27 | nd | nd | 2653 |
| Sme GR4 | pJB3Tc19 | GRI | B5 | nd | 0,18 | 21 | nd | nd |
| Sme GR4 | pJBpleD* | GRI | B6 | nd | 0,10 | 10 | nd | nd |
| M.loti MAFF303099 | pJB3Tc19 | GRI | B7 | 5766 | 0,50 | 49 | 117,8 | 11491 |
| M.loti MAFF303099 | pJBpleD* | GRI | B8 | 2399 | 0,05 | 12 | 195,6 | 46497 |
| Rle 8341 | pJB3Tc19 | GRI | B9 | 2462 | 1,09 | 25 | 99,4 | 2268 |
| Rle 8341 | pJBpleD* | GRI | B10 | nd | 0,04 | 7 | nd | nd |
| Rle UPM791 | pJB3Tc19 | GRI | B11 | 2953 | 0,45 | 58 | 50,6 | 6616 |
| Rle UPM791 | pJBpleD* | GRI | B12 | nd | 0,04 | 2 | nd | nd |
| S. fredii B1 | pJB3Tc19 | GRI | C1 | 1520 | 0,85 | 104 | 14,6 | 1794 |
| S. fredii B1 | pJBpleD* | GRI | C2 | 544 | 0,66 | 70 | 7,7 | 826 |
| S. fredii NGR234 | pJB3Tc19 | GRI | C3 | 2201 | 1,02 | 140 | 15,7 | 2165 |
| S. fredii NGR234 | pJBpleD* | GRI | C4 | nd | 0,06 | 8 | nd | nd |
| S. fredii USDA257 | pJB3Tc19 | GRI | C5 | 829 | 1,26 | 128 | 6,5 | 658 |
| S. fredii USDA257 | pJBpleD* | GRI | C6 | nd | 0,04 | 5 | nd | nd |
| Pto DC3000 | pJB3Tc19 | GRI | C7 | 207 | 0,49 | 142 | 1,5 | 423 |

| | | | | | | | | |
|---|---|---|---|---|---|---|---|---|
| Pto DC3000 | pJBpleD* | GRI | C8 | 250 | 0,41 | 135 | 1,9 | 605 |
| Pto DC3000 Δ*wssBC* | pJB3Tc19 | GRI | C9 | 227 | 0,60 | 191 | 1,2 | 375 |
| Pto DC3000 Δ*wssBC* | pJBpleD* | GRI | C10 | 309 | 0,54 | 180 | 1,7 | 569 |
| Pto DC3000 Δ*alg8* | pJB3Tc19 | GRI | C11 | 325 | 0,46 | 176 | 1,9 | 701 |
| Pto DC3000 Δ*alg8* | pJBpleD* | GRI | C12 | 300 | 0,41 | 106 | 2,8 | 732 |
| Pph 1448 | pJB3Tc19 | GRI | D1 | 101 | 0,13 | 49 | 2,1 | 750 |
| Pph 1448 | pJBpleD* | GRI | D2 | 213 | 0,09 | 15 | 13,8 | 2416 |
| Psv | pJB3Tc19 | GRI | D3 | 817 | 0,37 | 100 | 8,2 | 2214 |
| Psv | pJBpleD* | GRI | D4 | 1247 | 0,12 | 23 | 53,1 | 10504 |
| Pca SCRI1043 | pJB3Tc19 | GRI | D5 | 330 | 0,47 | 90 | 3,7 | 704 |
| Pca SCRI1043 | pJBpleD* | GRI | D6 | 354 | 0,41 | 57 | 6,2 | 864 |
| Pcc SCRI193 | pJB3Tc19 | GRI | D7 | 57 | 0,22 | 57 | 1,0 | 262 |
| Pcc SCRI193 | pJBpleD* | GRI | D8 | 17 | 0,22 | 63 | 0,3 | 79 |
| Atu LBA1010 | pJB3Tc19 | GRI | D9 | 465 | 0,30 | 36 | 12,8 | 1576 |
| Atu LBA1010 | pJBpleD* | GRI | D10 | nd | 0,04 | 6 | nd | nd |
| Mex PA1 | pJB3Tc19 | GRI | D11 | nd | 0,13 | 21 | nd | nd |
| Mex PA1 | pJBpleD* | GRI | D12 | nd | 0,08 | 12 | nd | nd |
| Mex AM1 | pJB3Tc19 | GRI | E1 | 16 | 0,13 | 16 | 1,0 | 120 |
| Mex AM1 | pJBpleD* | GRI | E2 | 81 | 0,14 | 19 | 4,3 | 582 |
| Abr AZ39 | pJB3Tc19 | GRI | E3 | 63 | 0,21 | 102 | 0,6 | 303 |
| Abr AZ39 | pJBpleD* | GRI | E4 | 36 | 0,12 | 35 | 1,0 | 293 |
| Kxy 7351 | pJB3Tc19 | GRI | E5 | 30 | 0,04 | 13 | 2,2 | 727 |
| Kxy 7351 | pJBpleD* | GRI | E6 | 89 | 0,04 | 8 | 11,7 | 2291 |
| A. acuatilis | pJB3Tc19 | GRI | E7 | 40 | 0,29 | 93 | 0,4 | 138 |
| A. acuatilis | pJBpleD* | GRI | E8 | 7 | 0,14 | 46 | 0,2 | 53 |
| Cupriavidus necator | pJB3Tc19 | GRI | E9 | 548 | 0,55 | 121 | 4,5 | 989 |
| Cupriavidus necator | pJBpleD* | GRI | E10 | 69 | 0,18 | 35 | 1,9 | 381 |
| Pkn B13 | pJB3Tc19 | GRI | E11 | 90 | 0,11 | 72 | 1,3 | 834 |
| Pkn B13 | pJBpleD* | GRI | E12 | 180 | 0,04 | 6 | 30,6 | 4622 |
| Delftia acidovorans | pJB3Tc19 | GRI | F1 | 287 | 0,22 | 65 | 4,4 | 1282 |
| Delftia acidovorans | pJBpleD* | GRI | F2 | 202 | 0,17 | 62 | 3,3 | 1162 |
| Ain LnG9092 | pJB3Tc19 | GRI | F3 | 102 | 0,11 | nd | nd | 959 |
| Ain LnG9092 | pJBpleD* | GRI | F4 | nd | 0,09 | 12 | nd | nd |

| | | | | | | | | |
|---|---|---|---|---|---|---|---|---|
| Ach denitrificans | pJB3Tc19 | GRI | F5 | nd | 0,26 | 51 | nd | nd |
| Ach denitrificans | pJBpleD* | GRI | F6 | nd | 0,08 | 10 | nd | nd |
| Sme 2011 | pJB3Tc19 | GRI | F7 | 583 | 0,51 | 69 | 8,5 | 1145 |
| Sme 2011 | pJBpleD* | GRI | F8 | 705 | 0,40 | 82 | 8,6 | 1771 |
| Sme 8530W (wfaB) | pJB3Tc19 | GRI | F9 | 1022 | 0,11 | 21 | 49,6 | 9406 |
| Sme 8530W (wfaB) | pJBpleD* | GRI | F10 | 986 | 0,62 | 54 | 18,3 | 1579 |
| Blank | | GRI | F11 | | | | | |
| Blank | | GRI | F12 | | | | | |
| GR-03 | pJB3Tc19 | GRI | G1 | 3053 | 1,00 | 136 | 22,5 | 3065 |
| GR-03 | pJBpleD* | GRI | G2 | 200 | 0,05 | 12 | 16,8 | 4301 |
| GR-05 | pJB3Tc19 | GRI | G3 | 2872 | 0,96 | 100 | 28,7 | 2992 |
| GR-05 | pJBpleD* | GRI | G4 | 71 | 0,04 | 6 | 11,0 | 1885 |
| GR-09 | pJB3Tc19 | GRI | G5 | 4060 | 1,33 | 157 | 25,8 | 3063 |
| GR-09 | pJBpleD* | GRI | G6 | 3171 | 0,35 | 35 | 90,1 | 9143 |
| GR-42 | pJB3Tc19 | GRI | G7 | 4786 | 0,59 | 55 | 86,3 | 8146 |
| GR-42 | pJBpleD* | GRI | G8 | 526 | 0,07 | 14 | 37,2 | 7759 |
| GR-45 | pJB3Tc19 | GRI | G9 | 2763 | 0,95 | 92 | 30,1 | 2907 |
| GR-45 | pJBpleD* | GRI | G10 | nd | 0,04 | 10 | nd | nd |
| GR-56 | pJB3Tc19 | GRI | G11 | 938 | 0,19 | 25 | 37,0 | 4817 |
| GR-56 | pJBpleD* | GRI | G12 | 23 | 0,04 | 7 | 3,3 | 597 |
| GR-60 | pJB3Tc19 | GRI | H1 | 4849 | 0,76 | 63 | 76,4 | 6358 |
| GR-60 | pJBpleD* | GRI | H2 | 105 | 0,04 | 7 | 14,4 | 2872 |
| GR-64 | pJB3Tc19 | GRI | H3 | 4284 | 1,14 | 135 | 31,7 | 3753 |
| GR-64 | pJBpleD* | GRI | H4 | 3437 | 0,31 | 36 | 96,1 | 11140 |
| GR-84 | pJB3Tc19 | GRI | H5 | 3425 | 0,77 | 68 | 50,7 | 4467 |
| GR-84 | pJBpleD* | GRI | H6 | nd | 0,04 | 7 | nd | nd |
| GR-87 | pJB3Tc19 | GRI | H7 | 2540 | 1,20 | 83 | 30,7 | 2115 |
| GR-87 | pJBpleD* | GRI | H8 | 24 | 0,04 | 7 | 3,3 | 623 |
| Blank | | GRI | H9 | | | | | |
| Blank | | GRI | H10 | | | | | |
| Blank | | GRI | H11 | | | | | |
| Blank | | GRI | H12 | | | | | |
| Sme SR1 | pJB3Tc19 | GRII | A1 | 1588 | 0,45 | 67 | 23,8 | 3501 |

| | | | | | | | | |
|---|---|---|---|---|---|---|---|---|
| Sme SR1 | pJBpleD* | GRII | A2 | 1990 | 0,28 | 22 | 90,6 | 7159 |
| Sme SR2 | pJB3Tc19 | GRII | A3 | 1267 | 0,43 | 49 | 26,1 | 2956 |
| Sme SR2 | pJBpleD* | GRII | A4 | 1151 | 0,34 | 24 | 48,4 | 3381 |
| Sme SR3 | pJB3Tc19 | GRII | A5 | 1218 | 0,44 | 78 | 15,5 | 2780 |
| Sme SR3 | pJBpleD* | GRII | A6 | 816 | 0,66 | 46 | 17,9 | 1239 |
| Sme SR4 | pJB3Tc19 | GRII | A7 | 1165 | 0,44 | 43 | 27,3 | 2631 |
| Sme SR4 | pJBpleD* | GRII | A8 | 1101 | 0,34 | 19 | 57,0 | 3202 |
| Sme SR9 | pJB3Tc19 | GRII | A9 | 1012 | 0,47 | 37 | 27,2 | 2167 |
| Sme SR9 | pJBpleD* | GRII | A10 | 993 | 0,20 | 15 | 67,5 | 5073 |
| Sme SR10 | pJB3Tc19 | GRII | A11 | 1573 | 0,46 | 73 | 21,5 | 3410 |
| Sme SR10 | pJBpleD* | GRII | A12 | 1273 | 0,40 | 43 | 29,8 | 3150 |
| Sme SR11 | pJB3Tc19 | GRII | B1 | 2299 | 0,46 | 192 | 12,0 | 5017 |
| Sme SR11 | pJBpleD* | GRII | B2 | 1703 | 0,35 | 108 | 15,8 | 4874 |
| Sme SR15 | pJB3Tc19 | GRII | B3 | 989 | 0,45 | 100 | 9,9 | 2178 |
| Sme SR15 | pJBpleD* | GRII | B4 | 860 | 0,47 | 32 | 27,3 | 1816 |
| Sme Cu9 | pJB3Tc19 | GRII | B5 | 1115 | 0,48 | 134 | 8,3 | 2304 |
| Sme Cu9 | pJBpleD* | GRII | B6 | 1129 | 0,41 | 91 | 12,4 | 2751 |
| Sme Cu10 | pJB3Tc19 | GRII | B7 | 1135 | 0,54 | 178 | 6,4 | 2118 |
| Sme Cu10 | pJBpleD* | GRII | B8 | 929 | 0,33 | 100 | 9,3 | 2819 |
| Blank | pJB3Tc19 | GRII | B9 | | | | | |
| Blank | pJBpleD* | GRII | B10 | | | | | |
| Blank | pJB3Tc19 | GRII | B11 | | | | | |
| Blank | pJBpleD* | GRII | B12 | | | | | |
| Sfr B1 | pJB3Tc19 | GRII | C1 | 774 | 0,74 | 212 | 3,7 | 1045 |
| Sfr B1 | pJBpleD* | GRII | C2 | 947 | 0,40 | 99 | 9,6 | 2379 |
| Sfr B8 | pJB3Tc19 | GRII | C3 | 718 | 0,68 | 167 | 4,3 | 1061 |
| Sfr B8 | pJBpleD* | GRII | C4 | 466 | 0,46 | 78 | 6,0 | 1022 |
| Sfr B20 | pJB3Tc19 | GRII | C5 | 1343 | 0,67 | 213 | 6,3 | 2002 |
| Sfr B20 | pJBpleD* | GRII | C6 | 350 | 0,28 | 13 | 26,7 | 1240 |
| Sfr B33 | pJB3Tc19 | GRII | C7 | 388 | 0,66 | 149 | 2,6 | 589 |
| Sfr B33 | pJBpleD* | GRII | C8 | 254 | 0,10 | 14 | 17,7 | 2525 |
| Sfr HH2 | pJB3Tc19 | GRII | C9 | 481 | 0,77 | 183 | 2,6 | 625 |
| Sfr HH2 | pJBpleD* | GRII | C10 | 101 | 0,06 | 43 | 2,4 | 1577 |

| | | | | | | | | |
|---|---|---|---|---|---|---|---|---|
| Sfr HH7 | pJB3Tc19 | GRII | C11 | 2324 | 0,51 | 167 | 13,9 | 4598 |
| Sfr HH7 | pJBpleD* | GRII | C12 | 317 | 0,04 | 15 | 21,7 | 8365 |
| Sfr HH28 | pJB3Tc19 | GRII | D1 | 1046 | 0,75 | 182 | 5,7 | 1395 |
| Sfr HH28 | pJBpleD* | GRII | D2 | 748 | 0,53 | 11 | 67,2 | 1416 |
| Sfr WH6 | pJB3Tc19 | GRII | D3 | 1820 | 0,78 | 232 | 7,9 | 2338 |
| Sfr WH6 | pJBpleD* | GRII | D4 | 691 | 0,39 | 43 | 16,1 | 1775 |
| Sfr WHG12 | pJB3Tc19 | GRII | D5 | 902 | 0,80 | 150 | 6,0 | 1123 |
| Sfr WHG12 | pJBpleD* | GRII | D6 | 703 | 0,11 | 23 | 31,2 | 6509 |
| Sfr WHG14 | pJB3Tc19 | GRII | D7 | 1397 | 1,44 | 91 | 15,4 | 970 |
| Sfr WHG14 | pJBpleD* | GRII | D8 | 664 | 0,17 | nd | nd | 3849 |
| Sfr HW1 | pJB3Tc19 | GRII | D9 | 423 | 0,93 | 150 | 2,8 | 457 |
| Sfr HW1 | pJBpleD* | GRII | D10 | 302 | 0,39 | 54 | 5,6 | 774 |
| Sfr HW14 | pJB3Tc19 | GRII | D11 | 1691 | 0,65 | 142 | 11,9 | 2599 |
| Sfr HW14 | pJBpleD* | GRII | D12 | 1536 | 0,47 | 115 | 13,4 | 3248 |
| Sfr HW22 | pJB3Tc19 | GRII | E1 | 2031 | 1,07 | 286 | 7,1 | 1893 |
| Sfr HW22 | pJBpleD* | GRII | E2 | 1242 | 1,13 | 175 | 7,1 | 1095 |
| Sfr WW10 | pJB3Tc19 | GRII | E3 | 615 | 0,92 | 208 | 3,0 | 671 |
| Sfr WW10 | pJBpleD* | GRII | E4 | 338 | 0,13 | 20 | 17,3 | 2635 |
| Sfr WWG11 | pJB3Tc19 | GRII | E5 | 1847 | 1,47 | 230 | 8,0 | 1260 |
| Sfr WWG11 | pJBpleD* | GRII | E6 | 1127 | 0,63 | 133 | 8,4 | 1791 |
| Sfr WWG35 | pJB3Tc19 | GRII | E7 | 1213 | 1,26 | 199 | 6,1 | 960 |
| Sfr WWG35 | pJBpleD* | GRII | E8 | 909 | 0,15 | 23 | 39,9 | 6249 |
| Sfr S1 | pJB3Tc19 | GRII | E9 | 1386 | 0,89 | 192 | 7,2 | 1558 |
| Sfr S1 | pJBpleD* | GRII | E10 | 720 | 0,43 | 78 | 9,3 | 1666 |
| Sfr S9 | pJB3Tc19 | GRII | E11 | 2001 | 0,89 | 202 | 9,9 | 2253 |
| Sfr S9 | pJBpleD* | GRII | E12 | 330 | 0,04 | 31 | 10,7 | 8598 |
| Sfr 16 | pJB3Tc19 | GRII | F1 | 1445 | 0,66 | 148 | 9,8 | 2200 |
| Sfr 16 | pJBpleD* | GRII | F2 | 868 | 0,06 | 42 | 20,5 | 14717 |
| Sfr S25 | pJB3Tc19 | GRII | F3 | 618 | 1,35 | 179 | 3,5 | 458 |
| Sfr S25 | pJBpleD* | GRII | F4 | 53 | 0,08 | 19 | 2,8 | 694 |
| Sfr S50 | pJB3Tc19 | GRII | F5 | 1008 | 0,61 | 72 | 13,9 | 1664 |
| Sfr S50 | pJBpleD* | GRII | F6 | 1075 | 0,91 | 84 | 12,9 | 1180 |
| Blank | | GRII | F7 | | | | | |

| | | | | | | | | |
|---|---|---|---|---|---|---|---|---|
| Blank | | GRII | F8 | | | | | |
| Blank | | GRII | F9 | | | | | |
| Blank | | GRII | F10 | | | | | |
| Blank | | GRII | F11 | | | | | |
| Blank | | GRII | F12 | | | | | |
| P.spp SJ04 | pJB3Tc19 | GRII | G1 | 64 | 0,64 | 160 | 0,4 | 99 |
| P.spp SJ04 | pJBpleD* | GRII | G2 | 122 | 0,33 | 36 | 3,4 | 372 |
| P.spp SJ08 | pJB3Tc19 | GRII | G3 | 183 | 0,75 | 104 | 1,8 | 246 |
| P.spp SJ08 | pJBpleD* | GRII | G4 | 216 | 0,35 | 115 | 1,9 | 618 |
| P.spp SJ13 | pJB3Tc19 | GRII | G5 | 75 | 0,19 | 139 | 0,5 | 406 |
| P.spp SJ13 | pJBpleD* | GRII | G6 | 159 | 0,19 | 119 | 1,3 | 836 |
| P.spp SJ16 | pJB3Tc19 | GRII | G7 | 207 | 0,15 | 51 | 4,1 | 1375 |
| P.spp SJ16 | pJBpleD* | GRII | G8 | 173 | 0,08 | 41 | 4,2 | 2079 |
| P.spp SJ17 | pJB3Tc19 | GRII | G9 | 256 | 0,19 | 12 | 20,6 | 1367 |
| P.spp SJ17 | pJBpleD* | GRII | G10 | 120 | 0,14 | 22 | 5,4 | 870 |
| P.spp SJ07b | pJB3Tc19 | GRII | G11 | 255 | 0,61 | 87 | 2,9 | 419 |
| P.spp SJ07b | pJBpleD* | GRII | G12 | 154 | 0,16 | 44 | 3,5 | 951 |
| P.spp SJ28 | pJB3Tc19 | GRII | H1 | 162 | 0,14 | 103 | 1,6 | 1138 |
| P.spp SJ28 | pJBpleD* | GRII | H2 | 122 | 0,05 | 56 | 2,2 | 2627 |
| P.spp SJ45 | pJB3Tc19 | GRII | H3 | 78 | 0,15 | 31 | 2,5 | 504 |
| P.spp SJ45 | pJBpleD* | GRII | H4 | 62 | 0,05 | 19 | 3,2 | 1251 |
| P.spp SJ48 | pJB3Tc19 | GRII | H5 | 106 | 0,18 | 210 | 0,5 | 601 |
| P.spp SJ48 | pJBpleD* | GRII | H6 | 63 | 0,13 | 131 | 0,5 | 481 |
| P.spp S5 | pJB3Tc19 | GRII | H7 | 271 | 0,44 | 291 | 0,9 | 610 |
| P.spp S5 | pJBpleD* | GRII | H8 | 417 | 0,47 | 266 | 1,6 | 884 |
| Blank | | GRII | H9 | | | | | |
| Blank | | GRII | H10 | | | | | |
| Blank | | GRII | H11 | | | | | |
| Blank | | GRII | H12 | | | | | |
| Blank | | MTP-IA | A1 | | | | | |
| Blank | | MTP-IA | A2 | | | | | |
| MTP I/A2/Pseudomonas | pJB3Tc19 | MTP-IA | A3 | 264 | 0,48 | 236 | 1,1 | 552 |
| MTP I/A2/Pseudomonas | pJBpleD* | MTP-IA | A4 | 418 | 0,43 | 255 | 1,6 | 975 |

| | | | | | | | | |
|---|---|---|---|---|---|---|---|---|
| MTP I/A3/Bacillus | pJB3Tc19 | MTP-IA | A5 | 387 | 0,38 | 140 | 2,8 | 1030 |
| MTP I/A3/Bacillus | pJBpleD* | MTP-IA | A6 | 236 | 0,14 | 186 | 1,3 | 1702 |
| MTP I/A4/Bacillus | pJB3Tc19 | MTP-IA | A7 | 1092 | 0,20 | 107 | 10,2 | 5398 |
| MTP I/A4/Bacillus | pJBpleD* | MTP-IA | A8 | 1508 | 0,25 | 221 | 6,8 | 5982 |
| Blank | | MTP-IA | A9 | | | | | |
| Blank | | MTP-IA | A10 | | | | | |
| MTP I/A6/Pseudomonas | pJB3Tc19 | MTP-IA | A11 | 56 | 0,38 | 144 | 0,4 | 147 |
| MTP I/A6/Pseudomonas | pJBpleD* | MTP-IA | A12 | 358 | 0,25 | 54 | 6,7 | 1441 |
| MTP I/B1/Curtobacterium | pJB3Tc19 | MTP-IA | B1 | 321 | 0,25 | 34 | 9,5 | 1268 |
| MTP I/B1/Curtobacterium | pJBpleD* | MTP-IA | B2 | 204 | 0,19 | 28 | 7,4 | 1057 |
| MTP I/B2/Bacillus | pJB3Tc19 | MTP-IA | B3 | 150 | 0,04 | nd | nd | 3477 |
| MTP I/B2/Bacillus | pJBpleD* | MTP-IA | B4 | 208 | 0,04 | nd | nd | 4723 |
| MTP I/B3/Curtobacterium | pJB3Tc19 | MTP-IA | B5 | 257 | 0,28 | 71 | 3,6 | 934 |
| MTP I/B3/Curtobacterium | pJBpleD* | MTP-IA | B6 | 268 | 0,25 | 101 | 2,6 | 1058 |
| MTP I/B4/Curtobacterium | pJB3Tc19 | MTP-IA | B7 | 356 | 0,09 | 11 | 31,3 | 3749 |
| MTP I/B4/Curtobacterium | pJBpleD* | MTP-IA | B8 | 158 | 0,13 | 20 | 7,8 | 1193 |
| MTP I/B5/Rahnella | pJB3Tc19 | MTP-IA | B9 | 3992 | 0,40 | 319 | 12,5 | 9901 |
| MTP I/B5/Rahnella | pJBpleD* | MTP-IA | B10 | 3364 | 0,41 | 310 | 10,9 | 8195 |
| MTP I/B6/Arthrobacter | pJB3Tc19 | MTP-IA | B11 | 262 | 0,04 | nd | nd | 5845 |
| MTP I/B6/Arthrobacter | pJBpleD* | MTP-IA | B12 | 283 | 0,04 | nd | nd | 6863 |
| MTP I/C1/Arthrobacter | pJB3Tc19 | MTP-IA | C1 | 132 | 0,09 | nd | nd | 1511 |
| MTP I/C1/Arthrobacter | pJBpleD* | MTP-IA | C2 | 275 | 0,05 | nd | nd | 5728 |
| MTP I/C2/Arthrobacter | pJB3Tc19 | MTP-IA | C3 | 99 | 0,06 | nd | nd | 1661 |
| MTP I/C2/Arthrobacter | pJBpleD* | MTP-IA | C4 | 194 | 0,05 | nd | nd | 3656 |
| MTP I/C3/Arthrobacter | pJB3Tc19 | MTP-IA | C5 | 103 | 0,04 | nd | nd | 2434 |
| MTP I/C3/Arthrobacter | pJBpleD* | MTP-IA | C6 | 95 | 0,04 | nd | nd | 2235 |
| MTP I/C4/Arthrobacter | pJB3Tc19 | MTP-IA | C7 | 240 | 0,04 | nd | nd | 5453 |
| MTP I/C4/Arthrobacter | pJBpleD* | MTP-IA | C8 | 385 | 0,04 | nd | nd | 9021 |
| MTP I/C5/Raoultella | pJB3Tc19 | MTP-IA | C9 | 2654 | 0,27 | 363 | 7,3 | 9926 |
| MTP I/C5/Raoultella | pJBpleD* | MTP-IA | C10 | 2142 | 0,21 | 123 | 17,4 | 10368 |
| MTP I/C6/Pseudomonas | pJB3Tc19 | MTP-IA | C11 | 325 | 0,04 | nd | nd | 7877 |
| MTP I/C6/Pseudomonas | pJBpleD* | MTP-IA | C12 | 248 | 0,04 | nd | nd | 5886 |
| MTP I/D1/Bacillus | pJB3Tc19 | MTP-IA | D1 | 573 | 0,26 | 52 | 11,0 | 2225 |

| | | | | | | | | |
|---|---|---|---|---|---|---|---|---|
| MTP I/D1/Bacillus | pJBpleD* | MTP-IA | D2 | 225 | 0,23 | nd | nd | 971 |
| Blank | | MTP-IA | D3 | | | | | |
| Blank | | MTP-IA | D4 | | | | | |
| MTP I/D3/Agrobacterium | pJB3Tc19 | MTP-IA | D5 | 2189 | 0,22 | 278 | 7,9 | 10118 |
| MTP I/D3/Agrobacterium | pJBpleD* | MTP-IA | D6 | 2413 | 0,45 | 114 | 21,2 | 5373 |
| MTP I/D4/Bacillus | pJB3Tc19 | MTP-IA | D7 | 2980 | 0,42 | 219 | 13,6 | 7175 |
| MTP I/D4/Bacillus | pJBpleD* | MTP-IA | D8 | 3456 | 0,44 | 230 | 15,0 | 7770 |
| MTP I/D5/Micrococcus | pJB3Tc19 | MTP-IA | D9 | 215 | 0,04 | nd | nd | 4850 |
| MTP I/D5/Micrococcus | pJBpleD* | MTP-IA | D10 | 351 | 0,05 | nd | nd | 7795 |
| MTP I/D6/Arthrobacter | pJB3Tc19 | MTP-IA | D11 | 224 | 0,07 | nd | nd | 3096 |
| MTP I/D6/Arthrobacter | pJBpleD* | MTP-IA | D12 | 283 | 0,05 | nd | nd | 5571 |
| MTP I/E1/Dyella | pJB3Tc19 | MTP-IA | E1 | 11 | 0,28 | nd | nd | 40 |
| MTP I/E1/Dyella | pJBpleD* | MTP-IA | E2 | 210 | 0,18 | 9 | 23,5 | 1195 |
| MTP I/E2/Sphingomonas | pJB3Tc19 | MTP-IA | E3 | 2068 | 1,18 | 691 | 3,0 | 1754 |
| MTP I/E2/Sphingomonas | pJBpleD* | MTP-IA | E4 | 1895 | 0,78 | 168 | 11,3 | 2440 |
| MTP I/E3/Herbaspirillum | pJB3Tc19 | MTP-IA | E5 | 57 | 0,30 | 131 | 0,4 | 188 |
| MTP I/E3/Herbaspirillum | pJBpleD* | MTP-IA | E6 | 165 | 0,11 | nd | nd | 1439 |
| MTP I/E4/Sphingomonas | pJB3Tc19 | MTP-IA | E7 | 3585 | 0,43 | 255 | 14,1 | 8418 |
| MTP I/E4/Sphingomonas | pJBpleD* | MTP-IA | E8 | 3886 | 0,42 | 245 | 15,9 | 9214 |
| MTP I/E5/Herbaspirillum | pJB3Tc19 | MTP-IA | E9 | 185 | 0,28 | nd | nd | 648 |
| MTP I/E5/Herbaspirillum | pJBpleD* | MTP-IA | E10 | 31 | 0,34 | nd | nd | 91 |
| MTP I/E6/Sphingomonas | pJB3Tc19 | MTP-IA | E11 | 1904 | 0,51 | 134 | 14,2 | 3728 |
| MTP I/E6/Sphingomonas | pJBpleD* | MTP-IA | E12 | 1622 | 0,51 | 178 | 9,1 | 3185 |
| MTP I/F1/Sphingomonas | pJB3Tc19 | MTP-IA | F1 | 1551 | 1,11 | 486 | 3,2 | 1396 |
| MTP I/F1/Sphingomonas | pJBpleD* | MTP-IA | F2 | 346 | 0,96 | 264 | 1,3 | 361 |
| MTP I/F2/Rhodobacter | pJB3Tc19 | MTP-IA | F3 | 417 | 0,04 | nd | nd | 9929 |
| MTP I/F2/Rhodobacter | pJBpleD* | MTP-IA | F4 | 252 | 0,05 | nd | nd | 5577 |
| MTP I/F3/Rhodobacter | pJB3Tc19 | MTP-IA | F5 | 49 | 0,04 | nd | nd | 1223 |
| MTP I/F3/Rhodobacter | pJBpleD* | MTP-IA | F6 | 71 | 0,04 | nd | nd | 1782 |
| MTP I/F4/Arthrobacter | pJB3Tc19 | MTP-IA | F7 | 69 | 0,04 | nd | nd | 1569 |
| MTP I/F4/Arthrobacter | pJBpleD* | MTP-IA | F8 | 69 | 0,04 | nd | nd | 1536 |
| MTP I/F5/Agrobacterium | pJB3Tc19 | MTP-IA | F9 | 1286 | 0,31 | 35 | 36,6 | 4094 |
| MTP I/F5/Agrobacterium | pJBpleD* | MTP-IA | F10 | 1033 | 0,20 | 24 | 43,1 | 5288 |

| | | | | | | | | |
|---|---|---|---|---|---|---|---|---|
| MTP I/F6/Ancylobacter | pJB3Tc19 | MTP-IA | F11 | 143 | 0,04 | nd | nd | 3320 |
| MTP I/F6/Ancylobacter | pJBpleD* | MTP-IA | F12 | 91 | 0,05 | nd | nd | 2005 |
| MTP I/G1/Agrobacterium | pJB3Tc19 | MTP-IA | G1 | 302 | 0,05 | nd | nd | 6297 |
| MTP I/G1/Agrobacterium | pJBpleD* | MTP-IA | G2 | 389 | 0,04 | nd | nd | 9295 |
| MTP I/G2/Sphingomonas | pJB3Tc19 | MTP-IA | G3 | 144 | 0,36 | 159 | 0,9 | 399 |
| MTP I/G2/Sphingomonas | pJBpleD* | MTP-IA | G4 | 255 | 0,33 | 141 | 1,8 | 763 |
| MTP I/G3/Paenibacillus | pJB3Tc19 | MTP-IA | G5 | 299 | 0,22 | 116 | 2,6 | 1380 |
| MTP I/G3/Paenibacillus | pJBpleD* | MTP-IA | G6 | 338 | 0,21 | 119 | 2,8 | 1586 |
| MTP I/G4/Paenibacillus | pJB3Tc19 | MTP-IA | G7 | 249 | 0,04 | nd | nd | 6186 |
| MTP I/G4/Paenibacillus | pJBpleD* | MTP-IA | G8 | 103 | 0,04 | nd | nd | 2384 |
| MTP I/G5/Agrobacterium | pJB3Tc19 | MTP-IA | G9 | 1815 | 0,29 | 15 | 124,6 | 6329 |
| MTP I/G5/Agrobacterium | pJBpleD* | MTP-IA | G10 | 173 | 0,05 | nd | nd | 3235 |
| MTP I/G6/Pseudomonas | pJB3Tc19 | MTP-IA | G11 | 112 | 0,07 | nd | nd | 1599 |
| MTP I/G6/Pseudomonas | pJBpleD* | MTP-IA | G12 | 314 | 0,20 | 59 | 5,3 | 1599 |
| Blank | | MTP-IA | H1 | | | | | |
| Blank | | MTP-IA | H2 | | | | | |
| MTP I/H2/Pseudomonas | pJB3Tc19 | MTP-IA | H3 | 231 | 0,32 | 128 | 1,8 | 713 |
| MTP I/H2/Pseudomonas | pJBpleD* | MTP-IA | H4 | 307 | 0,32 | 56 | 5,5 | 958 |
| MTP I/H3/Xanthomonas | pJB3Tc19 | MTP-IA | H5 | 2377 | 0,53 | 563 | 4,2 | 4474 |
| MTP I/H3/Xanthomonas | pJBpleD* | MTP-IA | H6 | 1942 | 0,64 | 447 | 4,3 | 3027 |
| MTP I/H4/Sinorhizobium | pJB3Tc19 | MTP-IA | H7 | 300 | 0,59 | 71 | 4,3 | 507 |
| MTP I/H4/Sinorhizobium | pJBpleD* | MTP-IA | H8 | 306 | 0,44 | 31 | 9,9 | 699 |
| MTP I/H5/Rhodococcus | pJB3Tc19 | MTP-IA | H9 | 328 | 0,26 | 32 | 10,2 | 1259 |
| MTP I/H5/Rhodococcus | pJBpleD* | MTP-IA | H10 | 306 | 0,31 | 15 | 20,9 | 974 |
| MTP I/H6/Rhodococcus | pJB3Tc19 | MTP-IA | H11 | 66 | 0,31 | 43 | 1,5 | 215 |
| MTP I/H6/Rhodococcus | pJBpleD* | MTP-IA | H12 | 168 | 0,41 | 46 | 3,7 | 412 |
| Blank | | MTP-IB | A1 | | | | | |
| Blank | | MTP-IB | A2 | | | | | |
| MTP I/A8/Pseudomonas | pJB3Tc19 | MTP-IB | A3 | 48 | 0,27 | 241 | 0,2 | 181 |
| MTP I/A8/Pseudomonas | pJBpleD* | MTP-IB | A4 | 237 | 0,18 | 165 | 1,4 | 1298 |
| MTP I/A9/Pseudomonas | pJB3Tc19 | MTP-IB | A5 | 246 | 0,25 | 225 | 1,1 | 970 |
| MTP I/A9/Pseudomonas | pJBpleD* | MTP-IB | A6 | 296 | 0,18 | 153 | 1,9 | 1632 |
| MTP I/A10/Pseudomonas | pJB3Tc19 | MTP-IB | A7 | 70 | 0,18 | 149 | 0,5 | 391 |

| | | | | | | | | | |
|---|---|---|---|---|---|---|---|---|---|
| MTP I/A10/Pseudomonas | pJBpleD* | MTP-IB | A8 | 159 | 0,18 | 130 | | 1,2 | 870 |
| Blank | | MTP-IB | A9 | | | | | | |
| Blank | | MTP-IB | A10 | | | | | | |
| Blank | | MTP-IB | A11 | | | | | | |
| Blank | | MTP-IB | A12 | | | | | | |
| MTP I/B7/Arthrobacter | pJB3Tc19 | MTP-IB | B1 | 113 | 0,06 | nd | nd | | 1788 |
| MTP I/B7/Arthrobacter | pJBpleD* | MTP-IB | B2 | 239 | 0,05 | nd | nd | | 5243 |
| MTP I/B8/Arthrobacter | pJB3Tc19 | MTP-IB | B3 | 205 | 0,04 | 40 | | 5,1 | 4701 |
| MTP I/B8/Arthrobacter | pJBpleD* | MTP-IB | B4 | 281 | 0,04 | 195 | | 1,4 | 6718 |
| MTP I/B9/Arthrobacter | pJB3Tc19 | MTP-IB | B5 | 311 | 0,45 | 385 | | 0,8 | 688 |
| MTP I/B9/Arthrobacter | pJBpleD* | MTP-IB | B6 | 386 | 0,39 | 306 | | 1,3 | 979 |
| MTP I/B10/Arthrobacter | pJB3Tc19 | MTP-IB | B7 | 326 | 0,04 | nd | nd | | 7826 |
| MTP I/B10/Arthrobacter | pJBpleD* | MTP-IB | B8 | 114 | 0,04 | nd | nd | | 2785 |
| MTP I/B11/Arthrobacter | pJB3Tc19 | MTP-IB | B9 | 98 | 0,04 | nd | nd | | 2326 |
| MTP I/B11/Arthrobacter | pJBpleD* | MTP-IB | B10 | 154 | 0,04 | nd | nd | | 3595 |
| MTP I/B12/Bacillus | pJB3Tc19 | MTP-IB | B11 | 113 | 0,05 | nd | nd | | 2254 |
| MTP I/B12/Bacillus | pJBpleD* | MTP-IB | B12 | 66 | 0,04 | nd | nd | | 1494 |
| MTP I/C7/Kozakia | pJB3Tc19 | MTP-IB | C1 | 3782 | 0,56 | 302 | | 12,5 | 6803 |
| MTP I/C7/Kozakia | pJBpleD* | MTP-IB | C2 | 3511 | 0,65 | 502 | | 7,0 | 5369 |
| MTP I/C8/Erwinia | pJB3Tc19 | MTP-IB | C3 | 578 | 0,32 | 314 | | 1,8 | 1809 |
| MTP I/C8/Erwinia | pJBpleD* | MTP-IB | C4 | 994 | 0,20 | 163 | | 6,1 | 4938 |
| MTP I/C9/Pseudomonas | pJB3Tc19 | MTP-IB | C5 | 372 | 0,43 | 362 | | 1,0 | 874 |
| MTP I/C9/Pseudomonas | pJBpleD* | MTP-IB | C6 | 210 | 0,43 | 297 | | 0,7 | 492 |
| MTP I/C10/Bacillus | pJB3Tc19 | MTP-IB | C7 | 219 | 0,04 | nd | nd | | 4954 |
| MTP I/C10/Bacillus | pJBpleD* | MTP-IB | C8 | 389 | 0,04 | nd | nd | | 9302 |
| MTP I/C11/Arthrobacter | pJB3Tc19 | MTP-IB | C9 | 427 | 0,04 | nd | nd | | 10248 |
| MTP I/C11/Arthrobacter | pJBpleD* | MTP-IB | C10 | 352 | 0,04 | nd | nd | | 8597 |
| MTP I/C12/Pseudomonas | pJB3Tc19 | MTP-IB | C11 | 358 | 0,40 | 431 | | 0,8 | 885 |
| MTP I/C12/Pseudomonas | pJBpleD* | MTP-IB | C12 | 440 | 0,16 | 57 | | 7,8 | 2760 |
| MTP I/D7/Micrococcus | pJB3Tc19 | MTP-IB | D1 | 451 | 0,34 | 105 | | 4,3 | 1328 |
| MTP I/D7/Micrococcus | pJBpleD* | MTP-IB | D2 | 349 | 0,40 | 198 | | 1,8 | 871 |
| MTP I/D8/Paracoccus | pJB3Tc19 | MTP-IB | D3 | 316 | 0,04 | nd | nd | | 7352 |
| MTP I/D8/Paracoccus | pJBpleD* | MTP-IB | D4 | 639 | 0,04 | 45 | | 14,2 | 15364 |

| | | | | | | | | |
|---|---|---|---|---|---|---|---|---|
| MTP I/D9/Pseudomonas | pJB3Tc19 | MTP-IB | D5 | 135 | 0,04 | nd | nd | 3180 |
| MTP I/D9/Pseudomonas | pJBpleD* | MTP-IB | D6 | 334 | 0,04 | nd | nd | 8191 |
| MTP I/D10/Pseudomonas | pJB3Tc19 | MTP-IB | D7 | 176 | 0,04 | nd | nd | 4132 |
| MTP I/D10/Pseudomonas | pJBpleD* | MTP-IB | D8 | 307 | 0,04 | nd | nd | 7605 |
| MTP I/D11/Burkholderia cepacia | pJB3Tc19 | MTP-IB | D9 | 475 | 0,58 | 601 | 0,8 | 813 |
| MTP I/D11/Burkholderia cepacia | pJBpleD* | MTP-IB | D10 | 382 | 0,63 | 637 | 0,6 | 606 |
| MTP I/D12/Microbacterium | pJB3Tc19 | MTP-IB | D11 | 371 | 0,29 | 200 | 1,9 | 1259 |
| MTP I/D12/Microbacterium | pJBpleD* | MTP-IB | D12 | 200 | 0,26 | 151 | 1,3 | 778 |
| MTP I/E7/Sphingomonas | pJB3Tc19 | MTP-IB | E1 | 432 | 0,50 | nd | nd | 860 |
| MTP I/E7/Sphingomonas | pJBpleD* | MTP-IB | E2 | 471 | 0,20 | nd | nd | 2387 |
| MTP I/E8/Herbaspirillum | pJB3Tc19 | MTP-IB | E3 | 359 | 0,27 | 204 | 1,8 | 1342 |
| MTP I/E8/Herbaspirillum | pJBpleD* | MTP-IB | E4 | 1452 | 0,23 | 166 | 8,7 | 6339 |
| MTP I/E9/Herbaspirillum | pJB3Tc19 | MTP-IB | E5 | 376 | 0,31 | 69 | 5,5 | 1212 |
| MTP I/E9/Herbaspirillum | pJBpleD* | MTP-IB | E6 | 231 | 0,31 | 22 | 10,7 | 745 |
| MTP I/E10/Burkholderia cepacia | pJB3Tc19 | MTP-IB | E7 | 323 | 0,31 | 145 | 2,2 | 1045 |
| MTP I/E10/Burkholderia cepacia | pJBpleD* | MTP-IB | E8 | 233 | 0,23 | 128 | 1,8 | 1006 |
| MTP I/E11/Kaistobacter | pJB3Tc19 | MTP-IB | E9 | 588 | 0,26 | 138 | 4,3 | 2299 |
| MTP I/E11/Kaistobacter | pJBpleD* | MTP-IB | E10 | 472 | 0,21 | 92 | 5,1 | 2298 |
| Blank | | MTP-IB | E11 | | | | | |
| Blank | | MTP-IB | E12 | | | | | |
| MTP I/F7/Microbacterium | pJB3Tc19 | MTP-IB | F1 | 625 | 0,64 | 223 | 2,8 | 971 |
| MTP I/F7/Microbacterium | pJBpleD* | MTP-IB | F2 | 739 | 0,66 | 253 | 2,9 | 1118 |
| MTP I/F8/Microbacterium | pJB3Tc19 | MTP-IB | F3 | 902 | 0,57 | 63 | 14,3 | 1576 |
| MTP I/F8/Microbacterium | pJBpleD* | MTP-IB | F4 | 801 | 0,59 | 198 | 4,0 | 1359 |
| MTP I/F9/Agrobacterium | pJB3Tc19 | MTP-IB | F5 | 1535 | 0,34 | 34 | 45,0 | 4518 |
| MTP I/F9/Agrobacterium | pJBpleD* | MTP-IB | F6 | 138 | 0,04 | 28 | 4,9 | 3288 |
| MTP I/F10/Arthrobacter | pJB3Tc19 | MTP-IB | F7 | 344 | 0,04 | nd | nd | 8185 |
| MTP I/F10/Arthrobacter | pJBpleD* | MTP-IB | F8 | 303 | 0,04 | nd | nd | 7641 |
| MTP I/F11/Microbacterium | pJB3Tc19 | MTP-IB | F9 | 6937 | 1,22 | 116 | 59,7 | 5702 |
| MTP I/F11/Microbacterium | pJBpleD* | MTP-IB | F10 | 6539 | 1,22 | 110 | 59,4 | 5377 |
| MTP I/F12/Microbacterium | pJB3Tc19 | MTP-IB | F11 | 355 | 0,23 | 178 | 2,0 | 1537 |
| MTP I/F12/Microbacterium | pJBpleD* | MTP-IB | F12 | 367 | 0,28 | 214 | 1,7 | 1319 |
| MTP I/G7/Paenibacillus | pJB3Tc19 | MTP-IB | G1 | 35 | 0,04 | nd | nd | 872 |

| | | | | | | | | |
|---|---|---|---|---|---|---|---|---|
| MTP I/G7/Paenibacillus | pJBpleD* | MTP-IB | G2 | 551 | 0,59 | 273 | 2,0 | 927 |
| MTP I/G8/Paenibacillus | pJB3Tc19 | MTP-IB | G3 | 190 | 0,04 | nd | nd | 4726 |
| MTP I/G8/Paenibacillus | pJBpleD* | MTP-IB | G4 | 212 | 0,04 | nd | nd | 5367 |
| MTP I/G9/Shewanella | pJB3Tc19 | MTP-IB | G5 | 196 | 0,04 | nd | nd | 4672 |
| MTP I/G9/Shewanella | pJBpleD* | MTP-IB | G6 | 330 | 0,05 | nd | nd | 6060 |
| MTP I/G10/Microbacterium | pJB3Tc19 | MTP-IB | G7 | 226 | 1,09 | 190 | 1,2 | 207 |
| MTP I/G10/Microbacterium | pJBpleD* | MTP-IB | G8 | 500 | 1,08 | 311 | 1,6 | 462 |
| MTP I/G11/Arthrobacter | pJB3Tc19 | MTP-IB | G9 | 129 | 0,04 | nd | nd | 3121 |
| MTP I/G11/Arthrobacter | pJBpleD* | MTP-IB | G10 | 103 | 0,04 | nd | nd | 2526 |
| MTP I/G12/Agrobacterium | pJB3Tc19 | MTP-IB | G11 | 2644 | 0,29 | 33 | 79,6 | 9165 |
| MTP I/G12/Agrobacterium | pJBpleD* | MTP-IB | G12 | 2571 | 0,26 | nd | nd | 9764 |
| MTP I/H7/Paenibacillus | pJB3Tc19 | MTP-IB | H1 | 402 | 0,41 | nd | nd | 985 |
| MTP I/H7/Paenibacillus | pJBpleD* | MTP-IB | H2 | 108 | 0,40 | nd | nd | 267 |
| MTP I/H8/Arthrobacter | pJB3Tc19 | MTP-IB | H3 | 289 | 0,05 | nd | nd | 6414 |
| MTP I/H8/Arthrobacter | pJBpleD* | MTP-IB | H4 | 416 | 0,05 | nd | nd | 8853 |
| MTP I/H9/Rhodococcus | pJB3Tc19 | MTP-IB | H5 | 357 | 0,59 | 435 | 0,8 | 608 |
| MTP I/H9/Rhodococcus | pJBpleD* | MTP-IB | H6 | 151 | 0,38 | 307 | 0,5 | 399 |
| MTP I/H10/Arthrobacter | pJB3Tc19 | MTP-IB | H7 | 330 | 0,04 | nd | nd | 7665 |
| MTP I/H10/Arthrobacter | pJBpleD* | MTP-IB | H8 | 160 | 0,04 | nd | nd | 3651 |
| MTP I/H11/Agrobacterium | pJB3Tc19 | MTP-IB | H9 | 1361 | 0,31 | 21 | 65,0 | 4417 |
| MTP I/H11/Agrobacterium | pJBpleD* | MTP-IB | H10 | 1011 | 0,60 | nd | nd | 1698 |
| MTP I/H12/Agrobacterium | pJB3Tc19 | MTP-IB | H11 | 2628 | 0,26 | 8 | 334,5 | 10183 |
| MTP I/H12/Agrobacterium | pJBpleD* | MTP-IB | H12 | 1835 | 0,24 | nd | nd | 7691 |
| Blank | | MTP-IIA | A1 | | | | | |
| Blank | | MTP-IIA | A2 | | | | | |
| MTP II/A2/Sphingomonas | pJB3Tc19 | MTP-IIA | A3 | 23 | 0,57 | 486 | 0,0 | 41 |
| MTP II/A2/Sphingomonas | pJBpleD* | MTP-IIA | A4 | 150 | 0,38 | 287 | 0,5 | 390 |
| MTP II/A3/Rhizobium | pJB3Tc19 | MTP-IIA | A5 | 148 | 0,51 | 428 | 0,3 | 288 |
| MTP II/A3/Rhizobium | pJBpleD* | MTP-IIA | A6 | 330 | 0,54 | 410 | 0,8 | 605 |
| MTP II/A4/Microbacterium | pJB3Tc19 | MTP-IIA | A7 | 626 | 0,74 | 443 | 1,4 | 842 |
| MTP II/A4/Microbacterium | pJBpleD* | MTP-IIA | A8 | 763 | 0,75 | 487 | 1,6 | 1024 |
| MTP II/A5/Ancylobacter | pJB3Tc19 | MTP-IIA | A9 | 353 | 0,48 | 159 | 2,2 | 741 |
| MTP II/A5/Ancylobacter | pJBpleD* | MTP-IIA | A10 | 189 | 0,42 | 101 | 1,9 | 448 |

| | | | | | | | | |
|---|---|---|---|---|---|---|---|---|
| MTP II/A6/Pseudomonas | pJB3Tc19 | MTP-IIA | A11 | 348 | 0,47 | 52 | 6,7 | 737 |
| MTP II/A6/Pseudomonas | pJBpleD* | MTP-IIA | A12 | 93 | 0,33 | 21 | 4,5 | 280 |
| MTP II/B1/Herbaspirillum | pJB3Tc19 | MTP-IIA | B1 | 104 | 0,52 | 227 | 0,5 | 198 |
| MTP II/B1/Herbaspirillum | pJBpleD* | MTP-IIA | B2 | 309 | 0,67 | 192 | 1,6 | 463 |
| MTP II/B2/Sphingomonas | pJB3Tc19 | MTP-IIA | B3 | 248 | 0,28 | 92 | 2,7 | 897 |
| MTP II/B2/Sphingomonas | pJBpleD* | MTP-IIA | B4 | 206 | 0,26 | 238 | 0,9 | 799 |
| MTP II/B3/Sphingomonas | pJB3Tc19 | MTP-IIA | B5 | 122 | 0,26 | 122 | 1,0 | 472 |
| MTP II/B3/Sphingomonas | pJBpleD* | MTP-IIA | B6 | 25 | 0,23 | 120 | 0,2 | 106 |
| MTP II/B4/Rhizobium | pJB3Tc19 | MTP-IIA | B7 | 1111 | 0,71 | 397 | 2,8 | 1560 |
| MTP II/B4/Rhizobium | pJBpleD* | MTP-IIA | B8 | 982 | 0,74 | 361 | 2,7 | 1333 |
| MTP II/B5/Paenibacillus | pJB3Tc19 | MTP-IIA | B9 | 188 | 0,56 | 468 | 0,4 | 337 |
| MTP II/B5/Paenibacillus | pJBpleD* | MTP-IIA | B10 | 307 | 0,51 | 421 | 0,7 | 606 |
| MTP II/B6/Paenibacillus | pJB3Tc19 | MTP-IIA | B11 | 227 | 0,48 | 244 | 0,9 | 470 |
| MTP II/B6/Paenibacillus | pJBpleD* | MTP-IIA | B12 | 273 | 0,38 | 139 | 2,0 | 721 |
| MTP II/C1/Microbacterium | pJB3Tc19 | MTP-IIA | C1 | 120 | 0,05 | nd | nd | 2403 |
| MTP II/C1/Microbacterium | pJBpleD* | MTP-IIA | C2 | 120 | 0,05 | 2 | 50,7 | 2650 |
| MTP II/C2/Microbacterium | pJB3Tc19 | MTP-IIA | C3 | 39 | 0,34 | 299 | 0,1 | 115 |
| MTP II/C2/Microbacterium | pJBpleD* | MTP-IIA | C4 | 1 | 0,05 | 15 | 0,1 | 18 |
| MTP II/C3/Microbacterium | pJB3Tc19 | MTP-IIA | C5 | 131 | 1,22 | 737 | 0,2 | 107 |
| MTP II/C3/Microbacterium | pJBpleD* | MTP-IIA | C6 | 159 | 1,26 | 736 | 0,2 | 126 |
| MTP II/C4/Microbacterium | pJB3Tc19 | MTP-IIA | C7 | 288 | 0,49 | 444 | 0,6 | 588 |
| MTP II/C4/Microbacterium | pJBpleD* | MTP-IIA | C8 | 82 | 0,46 | 424 | 0,2 | 178 |
| MTP II/C5/Microbacterium | pJB3Tc19 | MTP-IIA | C9 | 494 | 0,70 | 566 | 0,9 | 707 |
| MTP II/C5/Microbacterium | pJBpleD* | MTP-IIA | C10 | 327 | 0,64 | 540 | 0,6 | 513 |
| MTP II/C6/Microbacterium | pJB3Tc19 | MTP-IIA | C11 | 212 | 0,86 | 562 | 0,4 | 246 |
| MTP II/C6/Microbacterium | pJBpleD* | MTP-IIA | C12 | 406 | 0,85 | 563 | 0,7 | 480 |
| MTP II/D1/Bacillus | pJB3Tc19 | MTP-IIA | D1 | 236 | 0,58 | 504 | 0,5 | 409 |
| MTP II/D1/Bacillus | pJBpleD* | MTP-IIA | D2 | 351 | 0,56 | 536 | 0,7 | 629 |
| MTP II/D2/Bacillus | pJB3Tc19 | MTP-IIA | D3 | 204 | 0,53 | 519 | 0,4 | 386 |
| MTP II/D2/Bacillus | pJBpleD* | MTP-IIA | D4 | 150 | 0,92 | 566 | 0,3 | 163 |
| MTP II/D3/Rhodococcus | pJB3Tc19 | MTP-IIA | D5 | 332 | 0,70 | 437 | 0,8 | 474 |
| MTP II/D3/Rhodococcus | pJBpleD* | MTP-IIA | D6 | 318 | 0,68 | 490 | 0,6 | 466 |
| MTP II/D4/Pseudomonas | pJB3Tc19 | MTP-IIA | D7 | 275 | 0,35 | 295 | 0,9 | 787 |

| | | | | | | | | |
|---|---|---|---|---|---|---|---|---|
| MTP II/D4/Pseudomonas | pJBpleD* | MTP-IIA | D8 | 11 | 0,13 | 130 | 0,1 | 85 |
| MTP II/D5/Bacillus | pJB3Tc19 | MTP-IIA | D9 | 446 | 0,52 | 457 | 1,0 | 862 |
| MTP II/D5/Bacillus | pJBpleD* | MTP-IIA | D10 | 139 | 0,84 | 565 | 0,2 | 166 |
| MTP II/D6/Bacillus | pJB3Tc19 | MTP-IIA | D11 | 722 | 0,25 | 327 | 2,2 | 2900 |
| MTP II/D6/Bacillus | pJBpleD* | MTP-IIA | D12 | 2543 | 0,22 | 194 | 13,1 | 11644 |
| MTP II/E1/Burkholderia | pJB3Tc19 | MTP-IIA | E1 | 191 | 0,75 | 667 | 0,3 | 253 |
| MTP II/E1/Burkholderia | pJBpleD* | MTP-IIA | E2 | 337 | 0,73 | 416 | 0,8 | 460 |
| MTP II/E2/Microbacterium | pJB3Tc19 | MTP-IIA | E3 | 1164 | 0,92 | 541 | 2,2 | 1272 |
| MTP II/E2/Microbacterium | pJBpleD* | MTP-IIA | E4 | 1364 | 0,89 | 526 | 2,6 | 1526 |
| MTP II/E3/Microbacterium | pJB3Tc19 | MTP-IIA | E5 | 250 | 0,96 | 417 | 0,6 | 262 |
| MTP II/E3/Microbacterium | pJBpleD* | MTP-IIA | E6 | 181 | 1,06 | 582 | 0,3 | 170 |
| MTP II/E4/Microbacterium | pJB3Tc19 | MTP-IIA | E7 | 275 | 1,12 | 497 | 0,6 | 245 |
| MTP II/E4/Microbacterium | pJBpleD* | MTP-IIA | E8 | 374 | 1,28 | 524 | 0,7 | 293 |
| MTP II/E5/Microbacterium | pJB3Tc19 | MTP-IIA | E9 | 198 | 1,12 | 564 | 0,4 | 176 |
| MTP II/E5/Microbacterium | pJBpleD* | MTP-IIA | E10 | 225 | 1,26 | 615 | 0,4 | 179 |
| MTP II/E6/Burkholderia | pJB3Tc19 | MTP-IIA | E11 | 908 | 1,43 | 697 | 1,3 | 636 |
| MTP II/E6/Burkholderia | pJBpleD* | MTP-IIA | E12 | 995 | 1,41 | 683 | 1,5 | 703 |
| MTP II/F1/Agrobacterium | pJB3Tc19 | MTP-IIA | F1 | 722 | 0,22 | 174 | 4,1 | 3227 |
| MTP II/F1/Agrobacterium | pJBpleD* | MTP-IIA | F2 | 335 | 0,34 | 241 | 1,4 | 997 |
| MTP II/F2/Arthrobacter | pJB3Tc19 | MTP-IIA | F3 | 238 | 0,47 | 478 | 0,5 | 506 |
| MTP II/F2/Arthrobacter | pJBpleD* | MTP-IIA | F4 | 146 | 0,52 | 447 | 0,3 | 283 |
| Blank | | MTP-IIA | F5 | | | | | |
| Blank | | MTP-IIA | F6 | | | | | |
| MTP II/F4/Burkholderia | pJB3Tc19 | MTP-IIA | F7 | 813 | 0,78 | 470 | 1,7 | 1037 |
| MTP II/F4/Burkholderia | pJBpleD* | MTP-IIA | F8 | 771 | 0,77 | 464 | 1,7 | 1003 |
| MTP II/F5/Paenibacillus | pJB3Tc19 | MTP-IIA | F9 | 401 | 0,60 | 501 | 0,8 | 673 |
| MTP II/F5/Paenibacillus | pJBpleD* | MTP-IIA | F10 | 360 | 0,59 | 507 | 0,7 | 608 |
| MTP II/F6/Gluconacetobacter | pJB3Tc19 | MTP-IIA | F11 | 418 | 0,15 | 58 | 7,2 | 2822 |
| MTP II/F6/Gluconacetobacter | pJBpleD* | MTP-IIA | F12 | 165 | 0,26 | 145 | 1,1 | 630 |
| MTP II/G1/Beijerinckia indica | pJB3Tc19 | MTP-IIA | G1 | 350 | 0,58 | 377 | 0,9 | 609 |
| MTP II/G1/Beijerinckia indica | pJBpleD* | MTP-IIA | G2 | 201 | 0,54 | 536 | 0,4 | 371 |
| MTP II/G2/Bacillus | pJB3Tc19 | MTP-IIA | G3 | 154 | 0,49 | 477 | 0,3 | 313 |
| MTP II/G2/Bacillus | pJBpleD* | MTP-IIA | G4 | 36 | 0,51 | 495 | 0,1 | 71 |

| | | | | | | | | |
|---|---|---|---|---|---|---|---|---|
| MTP II/G3/Bacillus | pJB3Tc19 | MTP-IIA | G5 | 150 | 0,52 | 480 | 0,3 | 290 |
| MTP II/G3/Bacillus | pJBpleD* | MTP-IIA | G6 | 226 | 0,54 | 507 | 0,4 | 417 |
| MTP II/G4/Bacillus | pJB3Tc19 | MTP-IIA | G7 | 112 | 0,48 | 505 | 0,2 | 233 |
| MTP II/G4/Bacillus | pJBpleD* | MTP-IIA | G8 | 290 | 0,54 | 484 | 0,6 | 533 |
| MTP II/G5/Bacillus | pJB3Tc19 | MTP-IIA | G9 | 251 | 0,55 | 477 | 0,5 | 456 |
| MTP II/G5/Bacillus | pJBpleD* | MTP-IIA | G10 | 245 | 0,57 | 510 | 0,5 | 427 |
| MTP II/G6/Arthrobacter | pJB3Tc19 | MTP-IIA | G11 | 150 | 0,51 | 524 | 0,3 | 291 |
| MTP II/G6/Arthrobacter | pJBpleD* | MTP-IIA | G12 | 300 | 0,57 | 525 | 0,6 | 525 |
| MTP II/H1/Sphingomonas | pJB3Tc19 | MTP-IIA | H1 | 15 | 0,55 | 238 | 0,1 | 26 |
| MTP II/H1/Sphingomonas | pJBpleD* | MTP-IIA | H2 | 133 | 0,61 | 276 | 0,5 | 217 |
| MTP II/H2/Paenibacillus | pJB3Tc19 | MTP-IIA | H3 | 352 | 0,63 | 491 | 0,7 | 563 |
| MTP II/H2/Paenibacillus | pJBpleD* | MTP-IIA | H4 | 274 | 0,62 | 450 | 0,6 | 441 |
| MTP II/H3/Caulobacter | pJB3Tc19 | MTP-IIA | H5 | 195 | 0,48 | 465 | 0,4 | 404 |
| MTP II/H3/Caulobacter | pJBpleD* | MTP-IIA | H6 | 113 | 0,04 | nd | nd | 2755 |
| MTP II/H4/Bacillus | pJB3Tc19 | MTP-IIA | H7 | 246 | 0,52 | 423 | 0,6 | 473 |
| MTP II/H4/Bacillus | pJBpleD* | MTP-IIA | H8 | 205 | 0,55 | 467 | 0,4 | 372 |
| MTP II/H5/Xanthomonas | pJB3Tc19 | MTP-IIA | H9 | 180 | 0,53 | 429 | 0,4 | 339 |
| MTP II/H5/Xanthomonas | pJBpleD* | MTP-IIA | H10 | 361 | 0,55 | 443 | 0,8 | 652 |
| MTP II/H6/Agrobacterium | pJB3Tc19 | MTP-IIA | H11 | 2093 | 0,35 | 104 | 20,2 | 6036 |
| MTP II/H6/Agrobacterium | pJBpleD* | MTP-IIA | H12 | 563 | 0,34 | 277 | 2,0 | 1637 |
| MTP II/A7/Rhizobium | pJB3Tc19 | MTP-IIB | A1 | 382 | 0,35 | 170 | 2,3 | 1087 |
| MTP II/A7/Rhizobium | pJBpleD* | MTP-IIB | A2 | 334 | 0,53 | 458 | 0,7 | 630 |
| MTP II/A8/Microbacterium | pJB3Tc19 | MTP-IIB | A3 | 579 | 0,43 | 322 | 1,8 | 1339 |
| MTP II/A8/Microbacterium | pJBpleD* | MTP-IIB | A4 | 341 | 0,42 | 246 | 1,4 | 814 |
| MTP II/A9/Microbacterium | pJB3Tc19 | MTP-IIB | A5 | 2992 | 1,16 | 134 | 22,4 | 2580 |
| MTP II/A9/Microbacterium | pJBpleD* | MTP-IIB | A6 | 5932 | 1,31 | 241 | 24,7 | 4533 |
| MTP II/A10/Herbaspirillum | pJB3Tc19 | MTP-IIB | A7 | 4090 | 1,19 | 300 | 13,6 | 3440 |
| MTP II/A10/Herbaspirillum | pJBpleD* | MTP-IIB | A8 | 5793 | 1,23 | 173 | 33,5 | 4696 |
| MTP II/A11/Staphylococcus sp | pJB3Tc19 | MTP-IIB | A9 | 814 | 0,51 | 508 | 1,6 | 1605 |
| MTP II/A11/Staphylococcus sp | pJBpleD* | MTP-IIB | A10 | 553 | 0,51 | 516 | 1,1 | 1075 |
| MTP II/A12/Sphingomonas | pJB3Tc19 | MTP-IIB | A11 | 983 | 0,91 | 500 | 2,0 | 1082 |
| MTP II/A12/Sphingomonas | pJBpleD* | MTP-IIB | A12 | 633 | 1,01 | 200 | 3,2 | 629 |
| MTP II/B7/Herbaspirillum | pJB3Tc19 | MTP-IIB | B1 | 627 | 0,50 | 268 | 2,3 | 1242 |

| | | | | | | | | |
|---|---|---|---|---|---|---|---|---|
| MTP II/B7/Herbaspirillum | pJBpleD* | MTP-IIB | B2 | 799 | 0,61 | 214 | 3,7 | 1303 |
| MTP II/B8/Herbaspirillum | pJB3Tc19 | MTP-IIB | B3 | 757 | 0,64 | 575 | 1,3 | 1186 |
| MTP II/B8/Herbaspirillum | pJBpleD* | MTP-IIB | B4 | 633 | 0,65 | 162 | 3,9 | 975 |
| MTP II/B9/Bacillus | pJB3Tc19 | MTP-IIB | B5 | 688 | 0,45 | 538 | 1,3 | 1527 |
| MTP II/B9/Bacillus | pJBpleD* | MTP-IIB | B6 | 839 | 0,53 | 548 | 1,5 | 1583 |
| MTP II/B10/Microbacterium | pJB3Tc19 | MTP-IIB | B7 | 3992 | 0,67 | 249 | 16,0 | 5981 |
| MTP II/B10/Microbacterium | pJBpleD* | MTP-IIB | B8 | 3453 | 0,35 | 202 | 17,1 | 9932 |
| MTP II/B11/Microbacterium | pJB3Tc19 | MTP-IIB | B9 | 3824 | 0,75 | 361 | 10,6 | 5088 |
| MTP II/B11/Microbacterium | pJBpleD* | MTP-IIB | B10 | 3262 | 0,45 | 143 | 22,8 | 7215 |
| MTP II/B12/Klebsiella | pJB3Tc19 | MTP-IIB | B11 | 5478 | 0,63 | 504 | 10,9 | 8690 |
| MTP II/B12/Klebsiella | pJBpleD* | MTP-IIB | B12 | 5523 | 0,66 | 394 | 14,0 | 8309 |
| MTP II/C7/Microbacterium | pJB3Tc19 | MTP-IIB | C1 | 348 | 0,69 | 216 | 1,6 | 507 |
| MTP II/C7/Microbacterium | pJBpleD* | MTP-IIB | C2 | 1111 | 0,79 | 346 | 3,2 | 1402 |
| MTP II/C8/Microbacterium | pJB3Tc19 | MTP-IIB | C3 | 630 | 1,13 | 609 | 1,0 | 556 |
| MTP II/C8/Microbacterium | pJBpleD* | MTP-IIB | C4 | 654 | 1,11 | 684 | 1,0 | 589 |
| MTP II/C9/Microbacterium | pJB3Tc19 | MTP-IIB | C5 | 5582 | 0,61 | 466 | 12,0 | 9227 |
| MTP II/C9/Microbacterium | pJBpleD* | MTP-IIB | C6 | 4835 | 0,64 | 461 | 10,5 | 7588 |
| MTP II/C10/Microbacterium | pJB3Tc19 | MTP-IIB | C7 | 672 | 0,88 | 529 | 1,3 | 760 |
| MTP II/C10/Microbacterium | pJBpleD* | MTP-IIB | C8 | 594 | 0,87 | 524 | 1,1 | 684 |
| MTP II/C11/Microbacterium | pJB3Tc19 | MTP-IIB | C9 | 452 | 1,10 | 568 | 0,8 | 410 |
| MTP II/C11/Microbacterium | pJBpleD* | MTP-IIB | C10 | 679 | 1,06 | 499 | 1,4 | 642 |
| MTP II/C12/Pseudomonas | pJB3Tc19 | MTP-IIB | C11 | 558 | 0,92 | 539 | 1,0 | 609 |
| MTP II/C12/Pseudomonas | pJBpleD* | MTP-IIB | C12 | 512 | 1,01 | nd | nd | 509 |
| MTP II/D7/Bacillus | pJB3Tc19 | MTP-IIB | D1 | 3069 | 0,26 | 330 | 9,3 | 11818 |
| MTP II/D7/Bacillus | pJBpleD* | MTP-IIB | D2 | 2947 | 0,16 | 91 | 32,3 | 18677 |
| MTP II/D8/Brevundimonas | pJB3Tc19 | MTP-IIB | D3 | 283 | 1,17 | 675 | 0,4 | 241 |
| MTP II/D8/Brevundimonas | pJBpleD* | MTP-IIB | D4 | 279 | 1,09 | 593 | 0,5 | 257 |
| MTP II/D9/Microbacterium | pJB3Tc19 | MTP-IIB | D5 | 479 | 1,09 | 650 | 0,7 | 438 |
| MTP II/D9/Microbacterium | pJBpleD* | MTP-IIB | D6 | 516 | 1,05 | 633 | 0,8 | 491 |
| MTP II/D10/Rhodococcus | pJB3Tc19 | MTP-IIB | D7 | 560 | 0,63 | 568 | 1,0 | 882 |
| MTP II/D10/Rhodococcus | pJBpleD* | MTP-IIB | D8 | 578 | 0,65 | 566 | 1,0 | 883 |
| MTP II/D11/Pseudomonas | pJB3Tc19 | MTP-IIB | D9 | 378 | 0,43 | 156 | 2,4 | 872 |
| MTP II/D11/Pseudomonas | pJBpleD* | MTP-IIB | D10 | 507 | 0,31 | 141 | 3,6 | 1617 |

| | | | | | | | | |
|---|---|---|---|---|---|---|---|---|
| MTP II/D12/Brevundimonas | pJB3Tc19 | MTP-IIB | D11 | 765 | 0,55 | 603 | 1,3 | 1390 |
| MTP II/D12/Brevundimonas | pJBpleD* | MTP-IIB | D12 | 681 | 0,54 | 509 | 1,3 | 1266 |
| MTP II/E7/Rhodanobacter | pJB3Tc19 | MTP-IIB | E1 | 608 | 0,36 | 426 | 1,4 | 1669 |
| MTP II/E7/Rhodanobacter | pJBpleD* | MTP-IIB | E2 | 670 | 0,40 | 510 | 1,3 | 1655 |
| MTP II/E8/Burkholderia | pJB3Tc19 | MTP-IIB | E3 | 890 | 1,41 | 602 | 1,5 | 629 |
| MTP II/E8/Burkholderia | pJBpleD* | MTP-IIB | E4 | 624 | 1,43 | 572 | 1,1 | 437 |
| MTP II/E9/Bacillus | pJB3Tc19 | MTP-IIB | E5 | 495 | 0,57 | 631 | 0,8 | 862 |
| MTP II/E9/Bacillus | pJBpleD* | MTP-IIB | E6 | 600 | 0,57 | 653 | 0,9 | 1046 |
| MTP II/E10/Bacillus | pJB3Tc19 | MTP-IIB | E7 | 491 | 0,30 | 458 | 1,1 | 1636 |
| MTP II/E10/Bacillus | pJBpleD* | MTP-IIB | E8 | 734 | 0,69 | 771 | 1,0 | 1058 |
| MTP II/E11/Bacillus | pJB3Tc19 | MTP-IIB | E9 | 345 | 0,53 | 701 | 0,5 | 655 |
| MTP II/E11/Bacillus | pJBpleD* | MTP-IIB | E10 | 548 | 0,58 | 635 | 0,9 | 938 |
| MTP II/E12/Bacillus | pJB3Tc19 | MTP-IIB | E11 | 944 | 0,59 | 690 | 1,4 | 1614 |
| MTP II/E12/Bacillus | pJBpleD* | MTP-IIB | E12 | 672 | 0,58 | 514 | 1,3 | 1160 |
| MTP II/F7/Gluconacetobacter | pJB3Tc19 | MTP-IIB | F1 | 2447 | 0,47 | 444 | 5,5 | 5228 |
| MTP II/F7/Gluconacetobacter | pJBpleD* | MTP-IIB | F2 | 2132 | 0,42 | 474 | 4,5 | 5122 |
| MTP II/F8/Rahnella | pJB3Tc19 | MTP-IIB | F3 | 1099 | 0,24 | 100 | 11,0 | 4612 |
| MTP II/F8/Rahnella | pJBpleD* | MTP-IIB | F4 | 698 | 0,27 | 88 | 8,0 | 2609 |
| MTP II/F9/Rahnella | pJB3Tc19 | MTP-IIB | F5 | 710 | 0,27 | 114 | 6,2 | 2591 |
| MTP II/F9/Rahnella | pJBpleD* | MTP-IIB | F6 | 739 | 0,25 | 126 | 5,9 | 2924 |
| MTP II/F10/Rahnella | pJB3Tc19 | MTP-IIB | F7 | 986 | 0,25 | 110 | 9,0 | 4009 |
| MTP II/F10/Rahnella | pJBpleD* | MTP-IIB | F8 | 454 | 0,29 | 114 | 4,0 | 1583 |
| MTP II/F11/Rahnella | pJB3Tc19 | MTP-IIB | F9 | 1035 | 0,24 | 101 | 10,3 | 4329 |
| MTP II/F11/Rahnella | pJBpleD* | MTP-IIB | F10 | 677 | 0,26 | 118 | 5,8 | 2579 |
| MTP II/F12/Arthrobacter | pJB3Tc19 | MTP-IIB | F11 | 927 | 0,54 | 574 | 1,6 | 1723 |
| MTP II/F12/Arthrobacter | pJBpleD* | MTP-IIB | F12 | 642 | 0,57 | 492 | 1,3 | 1135 |
| MTP II/G7/Rhodococcus | pJB3Tc19 | MTP-IIB | G1 | 875 | 0,53 | 462 | 1,9 | 1646 |
| MTP II/G7/Rhodococcus | pJBpleD* | MTP-IIB | G2 | 809 | 0,57 | 625 | 1,3 | 1417 |
| MTP II/G8/Rhodococcus | pJB3Tc19 | MTP-IIB | G3 | 702 | 0,58 | 588 | 1,2 | 1219 |
| MTP II/G8/Rhodococcus | pJBpleD* | MTP-IIB | G4 | 875 | 0,58 | 644 | 1,4 | 1509 |
| MTP II/G9/Rhodococcus | pJB3Tc19 | MTP-IIB | G5 | 823 | 0,52 | 611 | 1,3 | 1579 |
| MTP II/G9/Rhodococcus | pJBpleD* | MTP-IIB | G6 | 1034 | 0,56 | 620 | 1,7 | 1859 |
| MTP II/G10/Nocardiopsis | pJB3Tc19 | MTP-IIB | G7 | 860 | 0,54 | 624 | 1,4 | 1606 |

| | | | | | | | | |
|---|---|---|---|---|---|---|---|---|
| MTP II/G10/Nocardiopsis | pJBpleD* | MTP-IIB | G8 | 628 | 0,55 | 639 | 1,0 | 1150 |
| MTP II/G11/Cellulosimicrobium | pJB3Tc19 | MTP-IIB | G9 | 535 | 0,73 | 617 | 0,9 | 729 |
| MTP II/G11/Cellulosimicrobium | pJBpleD* | MTP-IIB | G10 | 578 | 0,73 | 625 | 0,9 | 792 |
| MTP II/G12/Cellulosimicrobium | pJB3Tc19 | MTP-IIB | G11 | 1081 | 0,54 | 700 | 1,5 | 2005 |
| MTP II/G12/Cellulosimicrobium | pJBpleD* | MTP-IIB | G12 | 830 | 0,72 | 534 | 1,6 | 1145 |
| MTP II/H7/Xanthomonas | pJB3Tc19 | MTP-IIB | H1 | 2424 | 0,60 | 430 | 5,6 | 4069 |
| MTP II/H7/Xanthomonas | pJBpleD* | MTP-IIB | H2 | 1459 | 0,75 | 272 | 5,4 | 1943 |
| MTP II/H8/Xanthomonas | pJB3Tc19 | MTP-IIB | H3 | 2740 | 0,60 | 516 | 5,3 | 4591 |
| MTP II/H8/Xanthomonas | pJBpleD* | MTP-IIB | H4 | 909 | 0,63 | 296 | 3,1 | 1447 |
| MTP II/H9/Xanthomonas | pJB3Tc19 | MTP-IIB | H5 | 3403 | 0,65 | 451 | 7,5 | 5247 |
| MTP II/H9/Xanthomonas | pJBpleD* | MTP-IIB | H6 | 1747 | 0,74 | 397 | 4,4 | 2375 |
| MTP II/H10/Xanthomonas | pJB3Tc19 | MTP-IIB | H7 | 3117 | 0,59 | 568 | 5,5 | 5274 |
| MTP II/H10/Xanthomonas | pJBpleD* | MTP-IIB | H8 | 1382 | 0,52 | 340 | 4,1 | 2680 |
| MTP II/H11/Agrobacterium | pJB3Tc19 | MTP-IIB | H9 | 2746 | 0,23 | 93 | 29,5 | 11805 |
| MTP II/H11/Agrobacterium | pJBpleD* | MTP-IIB | H10 | 1638 | 0,31 | 104 | 15,8 | 5301 |
| MTP II/H12/Agrobacterium | pJB3Tc19 | MTP-IIB | H11 | 2144 | 0,37 | 35 | 61,6 | 5772 |
| MTP II/H12/Agrobacterium | pJBpleD* | MTP-IIB | H12 | 2557 | 0,29 | 28 | 90,9 | 8938 |
| Blank | | SPH | A1 | | | | | |
| Blank | | SPH | A2 | | | | | |
| Blank | | SPH | A3 | | | | | |
| Blank | | SPH | A4 | | | | | |
| Blank | | SPH | A5 | | | | | |
| Blank | | SPH | A6 | | | | | |
| Blank | | SPH | A7 | | | | | |
| Blank | | SPH | A8 | | | | | |
| Blank | | SPH | A9 | | | | | |
| Blank | | SPH | A10 | | | | | |
| Blank | | SPH | A11 | | | | | |
| Blank | | SPH | A12 | | | | | |
| Blank | | SPH | B1 | | | | | |
| Blank | | SPH | B2 | | | | | |
| Blank | | SPH | B3 | | | | | |
| Blank | | SPH | B4 | | | | | |

| | | | | | | | | |
|---|---|---|---|---|---|---|---|---|
| Blank | | SPH | B5 | | | | | |
| Blank | | SPH | B6 | | | | | |
| Blank | | SPH | B7 | | | | | |
| Blank | | SPH | B8 | | | | | |
| Blank | | SPH | B9 | | | | | |
| Blank | | SPH | B10 | | | | | |
| Blank | | SPH | B11 | | | | | |
| Blank | | SPH | B12 | | | | | |
| SphA1 | pJB3Tc19 | SPH | C1 | 417 | 0,04 | nd | nd | 11311 |
| SphA1 | pJBpleD* | SPH | C2 | 303 | 0,04 | nd | nd | 8092 |
| SphA3 | pJB3Tc19 | SPH | C3 | 301 | 0,04 | 2 | 164,3 | 7475 |
| SphA3 | pJBpleD* | SPH | C4 | 347 | 0,04 | nd | nd | 8659 |
| SphA4 | pJB3Tc19 | SPH | C5 | 1158 | 0,42 | 151 | 7,7 | 2779 |
| SphA4 | pJBpleD* | SPH | C6 | 377 | 0,04 | 9 | 42,2 | 9479 |
| SphA5 | pJB3Tc19 | SPH | C7 | 999 | 0,62 | 253 | 3,9 | 1605 |
| SphA5 | pJBpleD* | SPH | C8 | 608 | 0,96 | 63 | 9,6 | 631 |
| SphA8 | pJB3Tc19 | SPH | C9 | 393 | 0,04 | nd | nd | 10209 |
| SphA8 | pJBpleD* | SPH | C10 | 417 | 0,04 | 3 | 166,9 | 9817 |
| SphA9 | pJB3Tc19 | SPH | C11 | 388 | 0,04 | 6 | 66,5 | 9279 |
| SphA9 | pJBpleD* | SPH | C12 | 581 | 0,04 | nd | nd | 14885 |
| SphA11 | pJB3Tc19 | SPH | D1 | 346 | 0,05 | 2 | 185,3 | 7504 |
| SphA11 | pJBpleD* | SPH | D2 | 313 | 0,04 | 2 | 144,7 | 8496 |
| SphA12 | pJB3Tc19 | SPH | D3 | 245 | 0,04 | 5 | 53,6 | 6510 |
| SphA12 | pJBpleD* | SPH | D4 | 194 | 0,04 | 4 | 45,8 | 4792 |
| SphB1 | pJB3Tc19 | SPH | D5 | 206 | 0,04 | 13 | 15,4 | 4687 |
| SphB1 | pJBpleD* | SPH | D6 | 292 | 0,04 | 27 | 11,0 | 7380 |
| SphB2 | pJB3Tc19 | SPH | D7 | 329 | 0,06 | 9 | 37,1 | 5820 |
| SphB2 | pJBpleD* | SPH | D8 | 99 | 0,05 | 6 | 15,3 | 2146 |
| SphB5 | pJB3Tc19 | SPH | D9 | 412 | 0,04 | 3 | 143,6 | 10450 |
| SphB5 | pJBpleD* | SPH | D10 | 350 | 0,04 | 3 | 108,2 | 9301 |
| SphB6 | pJB3Tc19 | SPH | D11 | 1001 | 0,55 | 187 | 5,4 | 1807 |
| SphB6 | pJBpleD* | SPH | D12 | 1377 | 0,66 | 154 | 8,9 | 2086 |
| SphB7 | pJB3Tc19 | SPH | E1 | 1048 | 0,62 | 227 | 4,6 | 1688 |

| | | | | | | | | |
|---|---|---|---|---|---|---|---|---|
| SphB7 | pJBpleD* | SPH | E2 | 2319 | 0,31 | 81 | 28,7 | 7467 |
| SphB9 | pJB3Tc19 | SPH | E3 | 2913 | 0,56 | 91 | 31,8 | 5227 |
| SphB9 | pJBpleD* | SPH | E4 | 2027 | 0,18 | 32 | 63,6 | 11074 |
| SphC5 | pJB3Tc19 | SPH | E5 | 1029 | 0,57 | 301 | 3,4 | 1805 |
| SphC5 | pJBpleD* | SPH | E6 | 2410 | 1,73 | 866 | 2,8 | 1395 |
| CSph7 | pJB3Tc19 | SPH | E7 | 78 | 0,10 | 13 | 6,2 | 783 |
| SphC7 | pJBpleD* | SPH | E8 | 216 | 0,05 | 20 | 11,1 | 4375 |
| SphC8 | pJB3Tc19 | SPH | E9 | 899 | 0,72 | 174 | 5,2 | 1255 |
| SphC8 | pJBpleD* | SPH | E10 | 1127 | 0,50 | 108 | 10,4 | 2267 |
| SphC10 | pJB3Tc19 | SPH | E11 | 800 | 0,56 | 72 | 11,1 | 1423 |
| SphC10 | pJBpleD* | SPH | E12 | 4452 | 0,07 | 3 | 1483,9 | 65084 |
| SphD2 | pJB3Tc19 | SPH | F1 | 391 | 0,04 | 5 | 78,3 | 9986 |
| SphD2 | pJBpleD* | SPH | F2 | 241 | 0,05 | 22 | 11,2 | 5033 |
| SphD3 | pJB3Tc19 | SPH | F3 | 350 | 0,54 | 215 | 1,6 | 644 |
| SphD3 | pJBpleD* | SPH | F4 | 252 | 0,04 | 4 | 66,4 | 6534 |
| SphD4 | pJB3Tc19 | SPH | F5 | 334 | 0,79 | 137 | 2,4 | 420 |
| SphD4 | pJBpleD* | SPH | F6 | 132 | 1,55 | 213 | 0,6 | 85 |
| SphD5 | pJB3Tc19 | SPH | F7 | 210 | 0,04 | 3 | 79,9 | 5409 |
| SphD5 | pJBpleD* | SPH | F8 | 168 | 0,23 | 30 | 5,5 | 726 |
| SphD6 | pJB3Tc19 | SPH | F9 | 106 | 0,37 | 36 | 2,9 | 282 |
| SphD6 | pJBpleD* | SPH | F10 | 333 | 0,81 | 27 | 12,2 | 409 |
| SphD7 | pJB3Tc19 | SPH | F11 | 215 | 0,42 | 39 | 5,5 | 510 |
| SphD7 | pJBpleD* | SPH | F12 | 437 | 0,06 | 25 | 17,4 | 6924 |
| SphD8 | pJB3Tc19 | SPH | G1 | 514 | 0,04 | nd | nd | 13468 |
| SphD8 | pJBpleD* | SPH | G2 | 169 | 0,06 | nd | nd | 2654 |
| SphD9 | pJB3Tc19 | SPH | G3 | 1282 | 0,54 | 173 | 7,4 | 2367 |
| SphD9 | pJBpleD* | SPH | G4 | 1130 | 0,63 | 167 | 6,8 | 1790 |
| SphD10 | pJB3Tc19 | SPH | G5 | 295 | 0,70 | 300 | 1,0 | 419 |
| SphD10 | pJBpleD* | SPH | G6 | 2348 | 1,57 | 448 | 5,2 | 1497 |
| SphD11 | pJB3Tc19 | SPH | G7 | 200 | 0,37 | 69 | 2,9 | 536 |
| SphD11 | pJBpleD* | SPH | G8 | 223 | 0,05 | 34 | 6,6 | 4499 |
| SphE1 | pJB3Tc19 | SPH | G9 | 264 | 0,52 | 134 | 2,0 | 508 |
| SphE1 | pJBpleD* | SPH | G10 | 268 | 0,31 | 53 | 5,0 | 859 |

| | | | | | | | | |
|---|---|---|---|---|---|---|---|---|
| SphE2 | pJB3Tc19 | SPH | G11 | 361 | 0,86 | 215 | 1,7 | 418 |
| SphE2 | pJBpleD* | SPH | G12 | 428 | 0,75 | 153 | 2,8 | 568 |
| SphE3 | pJB3Tc19 | SPH | H1 | 911 | 0,48 | 148 | 6,2 | 1887 |
| SphE3 | pJBpleD* | SPH | H2 | 732 | 0,06 | 3 | 221,7 | 12464 |
| SphE5 | pJB3Tc19 | SPH | H3 | 175 | 0,61 | 245 | 0,7 | 285 |
| SphE5 | pJBpleD* | SPH | H4 | 227 | 0,78 | 290 | 0,8 | 290 |
| SphE6 | pJB3Tc19 | SPH | H5 | 425 | 0,66 | 310 | 1,4 | 640 |
| SphE6 | pJBpleD* | SPH | H6 | 499 | 1,81 | 827 | 0,6 | 276 |
| SphE8 | pJB3Tc19 | SPH | H7 | 921 | 0,04 | nd | nd | 23922 |
| SphE8 | pJBpleD* | SPH | H8 | 854 | 0,04 | 3 | 253,6 | 21666 |
| Blank | | | H9 | | | | | |
| Blank | | | H10 | | | | | |
| Blank | | | H11 | | | | | |
| Blank | | | H12 | | | | | |

*nd: Not detected

Table S4

| **Strains** | **Ratio TCC** | **Ratio TCC/TPC** | **Ratio TCC/OD** |
|---|---|---|---|
| Ret CFN42 | nd | nd | nd |
| Ret CFN42 ΔcelAB | nd | nd | nd |
| Ret CFN42 Δ363 | 0,02 | nd | 0,10 |
| Ret CFN42 ΔcelABΔ363 | nd | nd | nd |
| Sme 8530 | 1,32 | 0,98 | 1,17 |
| Sme 1021 | 0,85 | 0,83 | 0,79 |
| Sme 8530 sinI | 0,79 | 1,02 | 0,97 |
| Sme 8530 exoY | 0,73 | nd | 1,32 |
| Sme GR4 | nd | nd | nd |
| M.loti | 0,42 | 1,66 | 4,05 |
| Rle 8341 | nd | nd | nd |
| Rle UPM791 | nd | nd | nd |
| S. fredii | 0,36 | 0,53 | 0,46 |
| S. fredii NGR234 | nd | nd | nd |
| S. fredii USDA257 | nd | nd | nd |
| Pto DC3000 | 1,21 | 1,28 | 1,43 |
| Pto DC3000 ΔwssBC | 1,36 | 1,45 | 1,52 |
| Pto DC3000 Δalg8 | 0,92 | 1,53 | 1,04 |
| Pph 1448 | 2,12 | 6,69 | 3,22 |
| Psv | 1,53 | 6,48 | 4,74 |
| Pca | 1,07 | 1,69 | 1,23 |
| Pcc | 0,30 | 0,27 | 0,30 |
| Atu LBA1010 | nd | nd | nd |
| Mex PA1 | nd | nd | nd |
| Mex AM1 | 5,11 | 4,47 | 4,84 |
| Abr AZ39 | 0,57 | 1,68 | 0,97 |
| Kxy 7351 | 2,99 | 5,25 | 3,15 |
| A. acuatilis | 0,18 | 0,37 | 0,39 |
| Cupriavidus necator | 0,12 | 0,43 | 0,39 |
| Pkn B13 | 2,00 | 24,21 | 5,54 |
| Delftia acidovorans | 0,70 | 0,74 | 0,91 |
| Ain LnG9092 | nd | nd | nd |
| Ach denitrificans | nd | nd | nd |
| Sme 2011 (A. Becker) | 1,21 | 1,01 | 1,55 |
| Sme 8530W wfaB | 0,96 | 0,37 | 0,17 |
| GR-03 | 0,07 | 0,75 | 1,40 |
| GR-05 | 0,02 | 0,38 | 0,63 |
| GR-09 | 0,78 | 3,49 | 2,98 |
| GR-42 | 0,11 | 0,43 | 0,95 |
| GR-45 | nd | nd | nd |
| GR-56 | 0,02 | 0,09 | 0,12 |
| GR-60 | 0,02 | 0,19 | 0,45 |
| GR-64 | 0,80 | 3,03 | 2,97 |
| GR-84 | nd | nd | nd |
| GR-87 | 0,01 | 0,11 | 0,29 |
| Sme SR1 | 1,25 | 3,81 | 2,04 |
| Sme SR2 | 0,91 | 1,85 | 1,14 |
| Sme SR3 | 0,67 | 1,15 | 0,45 |

| | | | |
|---|---|---|---|
| Sme SR4 | 0,95 | 2,09 | 1,22 |
| Sme SR9 | 0,98 | 2,49 | 2,34 |
| Sme SR10 | 0,81 | 1,38 | 0,92 |
| Sme SR11 | 0,74 | 1,31 | 0,97 |
| Sme SR15 | 0,87 | 2,76 | 0,83 |
| Sme Cu9 | 1,01 | 1,49 | 1,19 |
| Sme Cu10 | 0,82 | 1,45 | 1,33 |
| Sfr B1 | 1,22 | 2,63 | 2,28 |
| Sfr B8 | 0,65 | 1,40 | 0,96 |
| Sfr B20 | 0,26 | 4,23 | 0,62 |
| Sfr B33 | 0,65 | 6,80 | 4,28 |
| Sfr HH2 | 0,21 | 0,90 | 2,52 |
| Sfr HH7 | 0,14 | 1,56 | 1,82 |
| Sfr HH28 | 0,72 | 11,72 | 1,02 |
| Sfr WH6 | 0,38 | 2,05 | 0,76 |
| Sfr WHG12 | 0,78 | 5,20 | 5,80 |
| Sfr WHG14 | 0,47 | nd | 3,97 |
| Sfr HW1 | 0,71 | 1,99 | 1,69 |
| Sfr HW14 | 0,91 | 1,12 | 1,25 |
| Sfr HW22 | 0,61 | 1,00 | 0,58 |
| Sfr WW10 | 0,55 | 5,86 | 3,93 |
| Sfr WWG11 | 0,61 | 1,05 | 1,42 |
| Sfr WWG35 | 0,75 | 6,54 | 6,51 |
| Sfr S1 | 0,52 | 1,29 | 1,07 |
| Sfr S9 | 0,17 | 1,08 | 3,82 |
| Sfr 16 | 0,60 | 2,10 | 6,69 |
| Sfr S25 | 0,09 | 0,82 | 1,51 |
| Sfr S50 | 1,07 | 0,92 | 0,71 |
| P.spp SJ04 | 1,93 | 8,59 | 3,74 |
| P.spp SJ08 | 1,18 | 1,06 | 2,51 |
| P.spp SJ13 | 2,11 | 2,47 | 2,06 |
| P.spp SJ16 | 0,84 | 1,03 | 1,51 |
| P.spp SJ17 | 0,47 | 0,26 | 0,64 |
| P.spp SJ07b | 0,60 | 1,20 | 2,27 |
| P.spp SJ28 | 0,75 | 1,37 | 2,31 |
| P.spp SJ45 | 0,80 | 1,28 | 2,48 |
| P.spp SJ48 | 0,59 | 0,96 | 0,80 |
| P.spp S5 | 1,54 | 1,68 | 1,45 |
| MTP I/A2/Pseudomonas | 1,59 | 1,47 | 1,76 |
| MTP I/A3/Bacillus | 0,61 | 0,46 | 1,65 |
| MTP I/A4/Bacillus | 1,38 | 0,67 | 1,11 |
| MTP I/A6/Pseudomonas | 6,45 | 17,32 | 9,82 |
| MTP I/A8/Pseudomonas | 0,64 | 0,78 | 0,83 |
| MTP I/A9/Pseudomonas | 1,39 | nd | 1,36 |
| MTP I/A10/Pseudomonas | 1,05 | 0,73 | 1,13 |
| MTP I/B1/Curtobacterium | 0,44 | 0,25 | 0,32 |
| MTP I/B2/Bacillus | 0,84 | 0,87 | 0,83 |
| MTP I/B3/Curtobacterium | 1,08 | nd | 1,17 |
| MTP I/B4/Curtobacterium | 2,09 | nd | 3,79 |
| MTP I/B5/Rahnella | 1,96 | nd | 2,20 |

| | | | |
|---|---|---|---|
| MTP I/B6/Arthrobacter | 0,92 | nd | 0,92 |
| MTP I/B7/Arthrobacter | 1,60 | nd | 1,65 |
| MTP I/B8/Arthrobacter | 0,81 | 2,38 | 1,04 |
| MTP I/B9/Arthrobacter | 0,76 | nd | 0,75 |
| MTP I/B10/Arthrobacter | 0,39 | nd | 0,44 |
| MTP I/B11/Arthrobacter | 1,10 | 2,69 | 0,53 |
| MTP I/B12/Bacillus | 1,16 | 1,10 | 1,08 |
| MTP I/C1/Arthrobacter | 1,63 | nd | 1,61 |
| MTP I/C2/Arthrobacter | 1,26 | nd | 1,80 |
| MTP I/C3/Arthrobacter | 18,86 | nd | 30,16 |
| MTP I/C4/Arthrobacter | 0,92 | 3,77 | 1,39 |
| MTP I/C5/Raoultella | 2,89 | nd | 7,64 |
| MTP I/C6/Pseudomonas | 1,08 | 1,13 | 1,09 |
| MTP I/C7/Kozakia | 0,17 | nd | 0,14 |
| MTP I/C8/Erwinia | 0,85 | 0,64 | 0,85 |
| MTP I/C9/Pseudomonas | 0,22 | 0,41 | 0,26 |
| MTP I/C10/Bacillus | 0,60 | nd | 0,56 |
| MTP I/C11/Arthrobacter | 1,45 | nd | 1,46 |
| MTP I/C12/Pseudomonas | 1,00 | nd | 0,98 |
| MTP I/D1/Bacillus | 0,80 | 1,18 | 1,29 |
| MTP I/D3/Agrobacterium | 0,63 | nd | 0,60 |
| MTP I/D4/Bacillus | 1,29 | nd | 1,48 |
| MTP I/D5/Micrococcus | 1,78 | 1,99 | 1,91 |
| MTP I/D6/Arthrobacter | 1,13 | 1,10 | 1,15 |
| MTP I/D7/Micrococcus | 0,42 | nd | 0,39 |
| MTP I/D8/Paracoccus | 0,10 | nd | 0,51 |
| MTP I/D9/Pseudomonas | 2,79 | nd | 1,00 |
| MTP I/D10/Pseudomonas | 1,33 | 3,06 | 1,34 |
| MTP I/D11/Burkholderia cepacia | 0,82 | 1,03 | 0,68 |
| MTP I/D12/Microbacterium | 1,02 | 2,32 | 1,38 |
| MTP I/E1/Dyella | 0,93 | 2,05 | 0,77 |
| MTP I/E2/Sphingomonas | 2,55 | 2,43 | 1,91 |
| MTP I/E3/Herbaspirillum | 4,89 | 7,15 | 7,16 |
| MTP I/E4/Sphingomonas | 1,20 | 1,78 | 1,68 |
| MTP I/E5/Herbaspirillum | 2,29 | 2,63 | 2,22 |
| MTP I/E6/Sphingomonas | 2,10 | nd | 2,93 |
| MTP I/E7/Sphingomonas | 1,37 | 0,28 | 1,43 |
| MTP I/E8/Herbaspirillum | 1,24 | 1,57 | 1,42 |
| MTP I/E9/Herbaspirillum | 0,35 | nd | 0,36 |
| MTP I/E10/Burkholderia cepacia | 1,57 | nd | 1,55 |
| MTP I/E11/Kaistobacter | 0,58 | nd | 0,66 |
| MTP I/F1/Sphingomonas | 0,93 | 0,56 | 0,79 |
| MTP I/F2/Rhodobacter | 1,72 | 3,32 | 2,73 |
| MTP I/F3/Rhodobacter | 0,57 | 0,69 | 0,56 |
| MTP I/F4/Arthrobacter | 1,78 | nd | 1,88 |
| MTP I/F5/Agrobacterium | 0,82 | nd | 0,84 |
| MTP I/F6/Ancylobacter | 1,23 | 9,35 | 3,12 |
| MTP I/F7/Microbacterium | 0,77 | 0,41 | 0,66 |
| MTP I/F8/Microbacterium | 2,02 | nd | 2,09 |
| MTP I/F9/Agrobacterium | 2,47 | nd | 2,58 |

| | | | |
|---|---|---|---|
| MTP I/F10/Arthrobacter | 1,75 | nd | 1,84 |
| MTP I/F11/Microbacterium | 0,80 | 0,76 | 0,74 |
| MTP I/F12/Microbacterium | 0,54 | 0,72 | 0,62 |
| MTP I/G1/Agrobacterium | 1,09 | nd | 2,78 |
| MTP I/G2/Sphingomonas | 4,04 | 4,97 | 4,72 |
| MTP I/G3/Paenibacillus | 0,62 | 1,94 | 0,61 |
| MTP I/G4/Paenibacillus | 0,72 | 0,81 | 0,96 |
| MTP I/G5/Agrobacterium | 0,80 | 1,20 | 1,00 |
| MTP I/G6/Pseudomonas | 1,18 | 1,04 | 1,15 |
| MTP I/G7/Paenibacillus | 0,89 | 0,28 | 0,86 |
| MTP I/G8/Paenibacillus | 0,09 | 0,11 | 0,73 |
| MTP I/G9/Shewanella | 0,88 | nd | 0,93 |
| MTP I/G10/Microbacterium | 0,94 | 0,99 | 0,94 |
| MTP I/G11/Arthrobacter | 1,03 | 0,86 | 0,86 |
| MTP I/G12/Agrobacterium | 15,67 | nd | 1,06 |
| MTP I/H2/Pseudomonas | 1,12 | nd | 1,14 |
| MTP I/H3/Xanthomonas | 1,68 | nd | 1,30 |
| MTP I/H4/Sinorhizobium | 2,21 | 1,35 | 2,23 |
| MTP I/H5/Rhodococcus | 0,80 | nd | 0,81 |
| MTP I/H6/Rhodococcus | 0,97 | nd | 1,07 |
| MTP I/H7/Paenibacillus | 0,27 | nd | 0,27 |
| MTP I/H8/Arthrobacter | 1,44 | nd | 1,38 |
| MTP I/H9/Rhodococcus | 0,42 | 0,60 | 0,66 |
| MTP I/H10/Arthrobacter | 0,48 | nd | 0,48 |
| MTP I/H11/Agrobacterium | 0,74 | nd | 0,38 |
| MTP I/H12/Agrobacterium | 0,70 | nd | 0,76 |
| MTP II/A2/Sphingomonas | 6,51 | 11,03 | 9,60 |
| MTP II/A3/Rhizobium | 2,23 | 2,33 | 2,10 |
| MTP II/A4/Microbacterium | 1,22 | 1,11 | 1,22 |
| MTP II/A5/Ancylobacter | 0,54 | 0,84 | 0,60 |
| MTP II/A6/Pseudomonas | 0,27 | 0,68 | 0,38 |
| MTP II/A7/Rhizobium | 2,98 | 3,52 | 2,34 |
| MTP II/A8/Microbacterium | 0,83 | 0,32 | 0,89 |
| MTP II/A9/Microbacterium | 0,20 | 0,21 | 0,23 |
| MTP II/A10/Herbaspirillum | 0,88 | 0,97 | 0,85 |
| MTP II/A11/Staphylococcus sp | 1,63 | 1,81 | 1,80 |
| MTP II/A12/Sphingomonas | 1,20 | 2,11 | 1,53 |
| MTP II/B1/Herbaspirillum | 1,00 | nd | 1,10 |
| MTP II/B2/Sphingomonas | 0,03 | 0,50 | 0,16 |
| MTP II/B3/Sphingomonas | 1,21 | 1,21 | 1,18 |
| MTP II/B4/Rhizobium | 0,29 | 0,30 | 0,30 |
| MTP II/B5/Paenibacillus | 0,66 | 0,69 | 0,73 |
| MTP II/B6/Paenibacillus | 1,92 | 1,91 | 1,95 |
| MTP II/B7/Herbaspirillum | 1,48 | 1,40 | 1,54 |
| MTP II/B8/Herbaspirillum | 0,73 | 0,67 | 0,42 |
| MTP II/B9/Bacillus | 0,96 | 0,85 | 0,98 |
| MTP II/B10/Microbacterium | 0,04 | 0,09 | 0,11 |
| MTP II/B11/Microbacterium | 0,31 | 0,25 | 0,19 |
| MTP II/B12/Klebsiella | 3,52 | 5,94 | 4,02 |
| MTP II/C1/Microbacterium | 1,77 | 2,83 | 1,82 |

| | | | |
|---|---|---|---|
| MTP II/C2/Microbacterium | 1,17 | 1,21 | 1,20 |
| MTP II/C3/Microbacterium | 0,72 | 0,52 | 0,65 |
| MTP II/C4/Microbacterium | 1,36 | 1,29 | 1,20 |
| MTP II/C5/Microbacterium | 1,14 | 1,04 | 1,02 |
| MTP II/C6/Microbacterium | 1,10 | 1,12 | 1,11 |
| MTP II/C7/Microbacterium | 0,46 | 0,33 | 0,31 |
| MTP II/C8/Microbacterium | 0,62 | 0,66 | 0,56 |
| MTP II/C9/Microbacterium | 0,95 | 0,96 | 0,97 |
| MTP II/C10/Microbacterium | 0,90 | 0,89 | 0,90 |
| MTP II/C11/Microbacterium | 0,39 | 0,16 | 0,22 |
| MTP II/C12/Pseudomonas | 0,57 | 0,40 | 0,61 |
| MTP II/D1/Bacillus | 0,24 | 0,23 | 0,23 |
| MTP II/D2/Bacillus | 1,51 | 1,43 | 1,44 |
| MTP II/D3/Rhodococcus | 2,58 | 2,69 | 2,28 |
| MTP II/D4/Pseudomonas | 0,98 | 0,91 | 0,94 |
| MTP II/D5/Bacillus | 2,00 | 2,00 | 1,80 |
| MTP II/D6/Bacillus | 9,10 | 7,84 | 8,22 |
| MTP II/D7/Bacillus | 0,78 | 0,85 | 0,78 |
| MTP II/D8/Brevundimonas | 0,58 | nd | 6,82 |
| MTP II/D9/Microbacterium | 0,83 | 0,75 | 0,79 |
| MTP II/D10/Rhodococcus | 2,01 | 1,95 | 1,93 |
| MTP II/D11/Pseudomonas | 0,27 | 0,10 | 0,27 |
| MTP II/D12/Brevundimonas | 0,87 | 0,32 | 0,58 |
| MTP II/E1/Burkholderia | 0,59 | 0,77 | 0,61 |
| MTP II/E2/Microbacterium | 1,98 | 1,10 | 1,76 |
| MTP II/E3/Microbacterium | 1,42 | 2,46 | 1,36 |
| MTP II/E4/Microbacterium | 0,68 | 0,67 | 0,67 |
| MTP II/E5/Microbacterium | 0,64 | 1,61 | 0,58 |
| MTP II/E6/Burkholderia | 1,28 | 1,60 | 1,05 |
| MTP II/E7/Rhodanobacter | 0,84 | 2,98 | 0,82 |
| MTP II/E8/Burkholderia | 1,22 | 1,19 | 1,04 |
| MTP II/E9/Bacillus | 0,87 | 1,07 | 1,66 |
| MTP II/E10/Bacillus | 0,85 | 2,15 | 1,42 |
| MTP II/E11/Bacillus | 1,01 | 1,29 | 0,96 |
| MTP II/E12/Bacillus | 3,20 | 1,99 | 2,77 |
| MTP II/F1/Agrobacterium | 1,04 | 0,93 | 1,06 |
| MTP II/F2/Arthrobacter | 0,87 | 0,88 | 0,82 |
| MTP II/F4/Burkholderia | 0,88 | 0,89 | 0,90 |
| MTP II/F5/Paenibacillus | 1,50 | 1,71 | 1,57 |
| MTP II/F6/Gluconacetobacter | 0,92 | nd | 0,84 |
| MTP II/F7/Gluconacetobacter | 0,96 | 3,47 | 1,58 |
| MTP II/F8/Rahnella | 0,99 | 1,12 | 1,07 |
| MTP II/F9/Rahnella | 1,08 | 1,11 | 1,12 |
| MTP II/F10/Rahnella | 1,03 | 1,04 | 1,00 |
| MTP II/F11/Rahnella | 1,34 | 1,49 | 1,86 |
| MTP II/F12/Arthrobacter | 0,89 | 1,05 | 0,91 |
| MTP II/G1/Beijerinckia indica | 1,10 | 0,92 | 0,99 |
| MTP II/G2/Bacillus | 0,70 | 0,74 | 0,69 |
| MTP II/G3/Bacillus | 1,21 | 1,17 | 1,21 |
| MTP II/G4/Bacillus | 1,49 | 0,89 | 0,65 |

| | | | |
|---|---|---|---|
| MTP II/G5/Bacillus | 1,59 | 1,76 | 1,43 |
| MTP II/G6/Arthrobacter | 0,71 | 0,95 | 0,72 |
| MTP II/G7/Rhodococcus | 0,87 | 0,82 | 0,98 |
| MTP II/G8/Rhodococcus | 0,64 | 0,72 | 0,57 |
| MTP II/G9/Rhodococcus | 1,04 | 0,95 | 1,13 |
| MTP II/G10/Nocardiopsis | 0,46 | 0,44 | 0,39 |
| MTP II/G11/Cellulosimicrobium | 0,65 | 0,56 | 0,60 |
| MTP II/G12/Cellulosimicrobium | 0,69 | 0,81 | 0,66 |
| MTP II/H1/Sphingomonas | 0,93 | 0,68 | 0,86 |
| MTP II/H2/Paenibacillus | 1,25 | 1,14 | 1,24 |
| MTP II/H3/Caulobacter | 1,26 | 1,24 | 1,18 |
| MTP II/H4/Bacillus | 0,73 | 0,71 | 0,72 |
| MTP II/H5/Xanthomonas | 1,08 | 1,07 | 1,09 |
| MTP II/H6/Agrobacterium | 0,77 | 1,01 | 0,57 |
| MTP II/H7/Xanthomonas | 0,60 | 0,95 | 0,48 |
| MTP II/H8/Xanthomonas | 0,33 | 0,58 | 0,32 |
| MTP II/H9/Xanthomonas | 0,51 | 0,58 | 0,45 |
| MTP II/H10/Xanthomonas | 0,44 | 0,74 | 0,51 |
| MTP II/H11/Agrobacterium | 0,60 | 0,53 | 0,45 |
| MTP II/H12/Agrobacterium | 1,19 | 1,47 | 1,55 |
| SphA1 | 0,73 | nd | 0,72 |
| SphA3 | 1,15 | nd | 1,16 |
| SphA4 | 0,33 | 5,50 | 3,41 |
| SphA5 | 0,61 | 2,44 | 0,39 |
| SphA8 | 1,06 | nd | 0,96 |
| SphA9 | 1,50 | nd | 1,60 |
| SphA11 | 0,91 | 0,78 | 1,13 |
| SphA12 | 0,79 | 0,86 | 0,74 |
| SphB1 | 1,42 | 0,71 | 1,57 |
| SphB2 | 0,30 | 0,41 | 0,37 |
| SphB5 | 0,85 | 0,75 | 0,89 |
| SphB6 | 1,38 | 1,66 | 1,15 |
| SphB7 | 2,21 | 6,21 | 4,42 |
| SphB9 | 0,70 | 2,00 | 2,12 |
| SphC5 | 2,34 | 0,81 | 0,77 |
| SphC7 | 2,76 | 1,80 | 5,59 |
| SphC8 | 1,25 | 2,02 | 1,81 |
| SphC10 | 5,56 | 133,3 | 45,74 |
| SphD2 | 0,62 | 0,14 | 0,50 |
| SphD3 | 0,72 | 40,81 | 10,14 |
| SphD4 | 0,40 | 0,25 | 0,20 |
| SphD5 | 0,80 | 0,07 | 0,13 |
| SphD6 | 3,16 | 4,18 | 1,45 |
| SphD7 | 2,03 | 3,15 | 13,57 |
| SphD8 | 0,33 | nd | 0,20 |
| SphD9 | 0,88 | 0,91 | 0,76 |
| SphD10 | 7,96 | 5,33 | 3,57 |
| SphD11 | 1,11 | 2,30 | 8,39 |
| SphE1 | 1,01 | 2,55 | 1,69 |
| SphE2 | 1,19 | 1,67 | 1,36 |

| | | | |
|---|---|---|---|
| SphE3 | 0,80 | 36,05 | 6,60 |
| SphE5 | 1,29 | 1,09 | 1,02 |
| SphE6 | 1,17 | 0,44 | 0,43 |
| SphE8 | 0,93 | nd | 0,91 |

*nd: Not determined
